# Engineered extracellular matrices reveal stiffness-mediated chemoresistance in patient-derived pancreatic cancer organoids

**DOI:** 10.1101/2022.04.22.488943

**Authors:** Bauer L. LeSavage, Aidan E. Gilchrist, Brad A. Krajina, Kasper Karlsson, Amber R. Smith, Kremena Karagyozova, Katarina C. Klett, Christina Curtis, Calvin J. Kuo, Sarah C. Heilshorn

## Abstract

Pancreatic ductal adenocarcinoma (PDAC) is characterized by its fibrotic and stiff extracellular matrix (ECM); however, the role that altered cell-ECM signaling may play in driving PDAC phenotype has historically been difficult to dissect. Here, we design an engineered matrix that recapitulates key hallmarks of the tumor ECM and show that patient-derived PDAC organoids develop gemcitabine chemoresistance when cultured within high stiffness matrices mechanically matched to *in vivo* tumors. Using genetic barcoding, we find that while matrix-specific clonal selection occurs, cellular heterogeneity is not the main driver of chemoresistance. Instead, stiffness-induced chemoresistance occurs due to the development of a plastic cancer stem cell phenotype – mediated by hyaluronan mechanosignaling – with increased expression of drug efflux transporters. Moreover, PDAC chemoresistance is reversible following transfer from high to low stiffness matrices, suggesting that mechanotherapeutics targeting the fibrotic ECM may sensitize chemoresistant tumors. Overall, we demonstrate the power of engineered matrices and patient-derived organoids to elucidate how ECM properties influence human disease pathophysiology.

## Introduction

Pancreatic ductal adenocarcinoma (PDAC) is a lethal disease with an overall 5-year survival rate less than 7%^1,2^. Frontline chemotherapeutics to treat PDAC are often ineffective due to intrinsic or acquired resistance^3^, and no approved chemotherapies have substantially extended PDAC patient survival since the regulatory approval of gemcitabine in 1996^4,5^. Therefore, understanding the mechanisms of how PDAC tumors develop and retain chemoresistance is critical to advancing impactful treatment strategies.

Clinically, PDAC chemoresistance has been independently correlated with both the presence of highly plastic cancer stem cells (CSCs)^6,7^ and extreme fibrotic remodeling of the extracellular matrix (ECM) (i.e. desmoplasia)^8–10^. CSCs are a stem-like population with enhanced capacity for tumor initiation, metastasis, and chemoresistance, whose presence has been linked with decreased patient survival^11^. The PDAC ECM is characterized by its high stiffness and dense stroma, which worsens throughout disease progression^8–10^. In PDAC, these matrix properties and the associated high interstitial fluid pressures are largely thought to impact chemoresistance by acting as a physical barrier that limits drug delivery to the tumor^12,13^. Promisingly, recent PDAC studies have shown that co-administration of chemotherapies and ECM-depleting factors can significantly improve survival rates in mice^12,14^. However, these therapies based on animal models have yet to show efficacy in human PDAC trials^15–17^, suggesting the ECM may be playing an additional role in driving PDAC chemoresistance.

As these fibrotic ECM properties are known to directly influence cancer cell phenotypes in other tumor types (notably breast cancer)^18–21^, we hypothesized that the PDAC ECM may drive CSC enrichment and/or chemoresistance through direct cell-ECM signaling. Unfortunately, animal models are not ideal for studying reciprocal cell-ECM interactions as they are costly, low-throughput, and offer limited experimental control over the matrix microenvironment. Alternatively, human cancer organoids offer a relatively cost-effective and representative model of patient-specific tumors and their matrix^22–25^. However, traditional methods for culturing cancer organoids *in vitro* rely on animal-derived ECMs such as basement membrane extract (e.g. Matrigel/Cultrex) that are ill-defined and non-tunable^22,26^, which limits the ability to uncover mechanistic links between ECM properties and chemoresistance. Notably, Matrigel-alternatives are currently being explored for both healthy and cancer organoid models^27–31^, yet have not been applied to understanding matrix-mediated PDAC chemoresistance.

Here, we engineer a defined and tunable 3D matrix that mimics key biochemical and mechanical properties of the *in vivo* PDAC ECM and supports long-term culture and passaging of primary human patient-derived PDAC organoids *in vitro*. Using our engineered matrix, we identify previously untested causal relationships between PDAC chemoresistance and the ECM, which opens new directions of drug development for effective patient therapy.

## Results

### 3D extracellular matrix stiffness drives PDAC organoid chemoresistance

The *in vivo* PDAC ECM is characterized by the increased deposition of several matrix components including fibronectin and hyaluronan (HA)^8–10^ (**Fig. 1a**), a linear polysaccharide commonly found in connective and epithelial tissues. RNA-seq analysis of 179 PDAC and 171 normal pancreas tissues collected from The Cancer Genome Atlas and Genome-Tissue Expression Project^32^ confirmed increased expression of fibronectin (*FN1*) and HA synthesis genes (*HAS1, HAS2, HAS3, UGDH*) in the tumor microenvironment (**Fig. 1b**). Increased matrix deposition also correlated with increased tumor stiffness (∼2900 Pa) compared to normal pancreas tissue (∼900 Pa), as measured by bulk shear rheology on freshly resected human samples (**Fig. 1c**).

**Figure 1.**
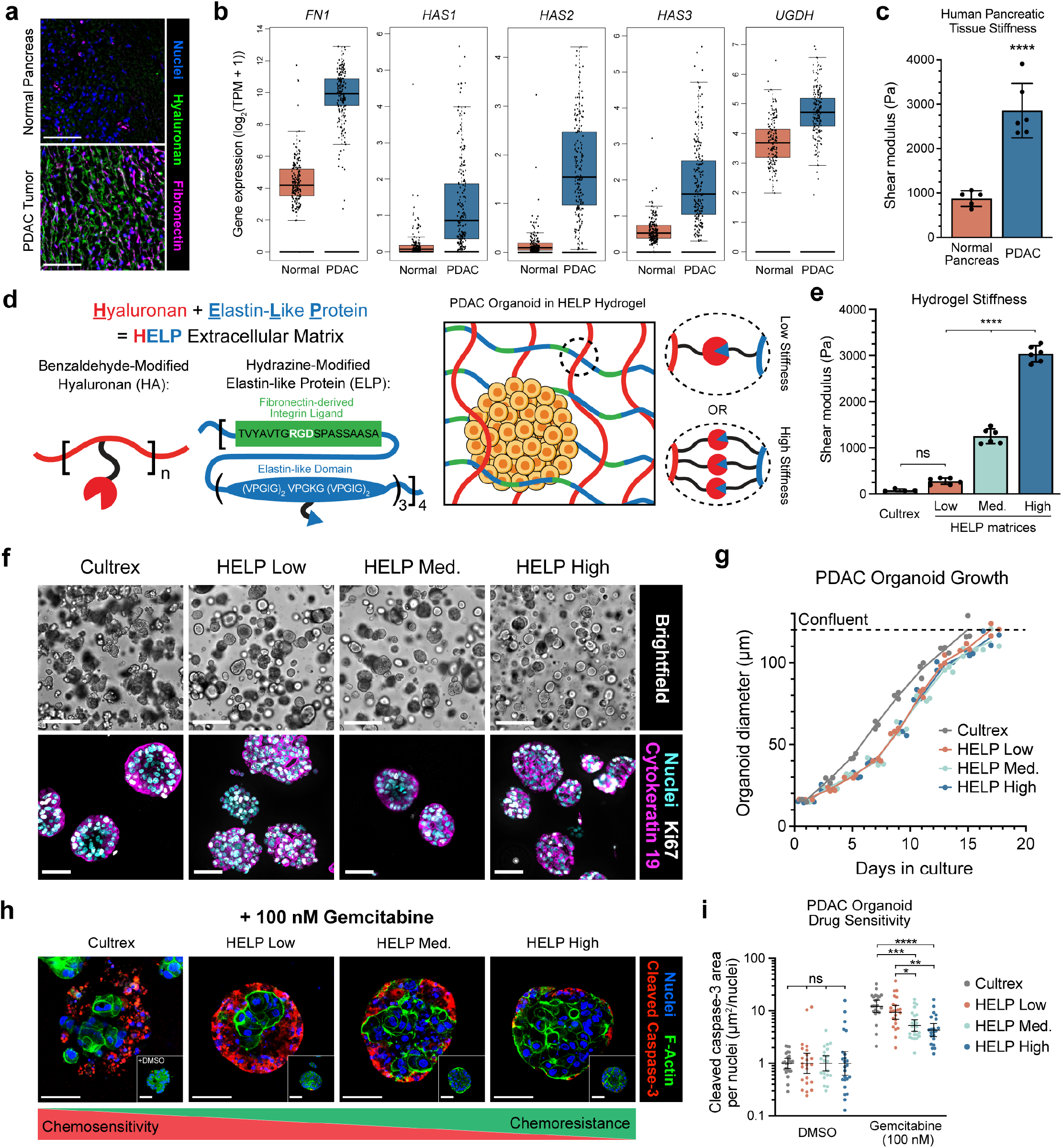
3D extracellular matrix stiffness drives PDAC organoid chemoresistance. **a**, Representative immunofluorescence (IF) images of human PDAC and normal adjacent pancreatic tissue stained for extracellular matrix components. Scale bar, 100 μm. **b**, Bulk RNA-sequencing analysis of PDAC and normal pancreatic tissue samples for fibronectin (*FN1*) and hyaluronan associated genes (*HAS1, HAS2, HAS3, UGDH*; N=179 for PDAC, N=171 for Normal). **c**, Stiffness measurements of fresh, surgically-resected human PDAC and normal pancreatic tissue (N=5-6, mean ± SD, unpaired two-tailed Student’s t-test, ****p<0.0001). **d**, Schematic of engineered HELP matrix comprising chemically-modified hyaluronan and elastin-like protein. **e**, Stiffness measurements of Cultrex and HELP matrices (N=4-6, mean ± SD, ordinary one-way ANOVA with Tukey multiple comparisons correction, ****p<0.0001, ns = not significant). Low, Medium, or High labels refer to relative stiffness values. **f**, Representative brightfield (BF; top) and IF (bottom) images of PDAC organoids cultured within Cultrex and HELP matrices. BF scale bar, 250 μm; IF scale bar, 50 μm. **g**, Quantification of PDAC organoid diameter during culture within Cultrex and HELP matrices. Each data point represents the average organoid diameter from one hydrogel at the given timepoint (N=3 replicate hydrogels, Cultrex: n=77-164; HELP Low: n=84-206; HELP Medium: n=93-284; HELP High: n=85-223 organoids per individual hydrogel per timepoint). No statistical difference was measured across HELP matrices of varying stiffness at each timepoint using ordinary one-way ANOVA with Tukey multiple comparisons correction. Dashed line represents approximate organoid diameter where cultures are considered confluent. **h**, Representative IF images of PDAC organoids cultured within Cultrex and HELP matrices and treated with DMSO control (inset) or 100 nM gemcitabine for three days following organoid formation (cleaved caspase-3 = apoptosis marker). Scale bars, 50 μm. **i**, Quantification of cleaved caspase-3 signal normalized to DMSO control. Each data point represents the average cleaved caspase-3 area per nuclei from one confocal z-stack containing several organoids (data compiled from N=3 replicate hydrogels, n=5-8 z-stacks per hydrogel, geometric mean ± 95% confidence interval, ordinary one-way ANOVA with Tukey multiple comparisons correction, *p<0.05, **p<0.01, ***p<0.001, ****p<0.0001, ns = not significant).

To model these biochemical components and mechanical features of the PDAC ECM *in vitro*, we designed a defined and tunable 3D matrix comprised of HA and elastin-like protein (ELP), which we term HELP^28^ (**Fig. 1d**). ELP is a recombinant protein consisting of repeating elastin- and fibronectin-derived sequences that include an RGD integrin-binding motif^33^. By tuning the polymer concentration and chemical crosslink density between benzaldehyde and hydrazine groups on the HA and ELP, respectively, we created a physiologically-relevant range of HELP matrices that span from the low stiffness of Cultrex hydrogel controls to the high stiffness of PDAC tumor tissue (**Fig. 1e, Supplementary Fig. 1**,**2**), while maintaining identical HA content and RGD concentration (**Supplementary Table 1**,**2**). We termed these formulations HELP Low, HELP Medium, and HELP High, corresponding to their stiffness.

For all hydrogel encapsulations, patient-derived PDAC organoids were dissociated into single cell suspensions and encapsulated within either Cultrex or HELP matrices of varying stiffness. Organoids showed robust expansion in all matrices and stained positive for cytokeratin 19 (ductal lineage) and Ki67 (proliferation) (**Fig. 1f**). PDAC organoids encapsulated in HELP matrices had an initial delay in proliferation compared to Cultrex hydrogels, yet similar overall diameter growth rate over 7-14 days (**Fig. 1g, Supplementary Fig. 3a**,**b**).

To test how matrix stiffness impacts PDAC drug sensitivity, we treated ∼75-μm diameter PDAC organoids with 100 nM gemcitabine, a nucleoside analog commonly prescribed for clinical treatment of PDAC tumors^5^, for 3 days. We found organoids cultured in stiff HELP High matrices had lower expression of apoptotic marker cleaved caspase-3 and were more resistant to gemcitabine treatment compared to softer HELP Low and Cultrex matrices (**Fig. 1h,i**).

To explore the possibility that this difference in drug response was due to differences in cell proliferation rate, we performed cell cycle analysis on PDAC organoids grown within Cultrex, HELP Low, and HELP High matrices. In agreement with Ki67 and organoid diameter data, we saw no significant difference in percentage of proliferative cells in s-phase (**Supplementary Fig. 3-5**). Additionally, all hydrogels had similar diffusion of macromolecules ranging in size from 10-250 kDa, ensuring no differences in gemcitabine (263 Da) delivery to organoids grown in different matrices (**Supplementary Fig. 6)**. From these data, we concluded that matrix stiffness is a key driver of PDAC organoid chemoresistance in our model.

### Prolonged exposure to high stiffness mediates PDAC organoid chemoresistance

To explore the dynamic onset of PDAC chemoresistance over long-term exposure to high stiffness, we cultured PDAC organoids in HELP Low or HELP High matrices and tested their drug sensitivity on a subset of previously untreated cells following one and four passages. Specifically, cells were treated with gemcitabine following the formation of ∼75-μm diameter organoids (**Fig. 2a**). Interestingly, after four passages within HELP High, PDAC organoids showed increased resistance to gemcitabine compared to treatment during passage one, resulting in a doubling of the IC_50_ value (**Fig. 2b,c**). Expansion of PDAC organoids in soft HELP Low and Cultrex matrices for four passages did not result in a statistically significant difference in gemcitabine IC_50_ (**Fig. 2b,c, Supplementary Fig. 7**).

**Figure 2.**
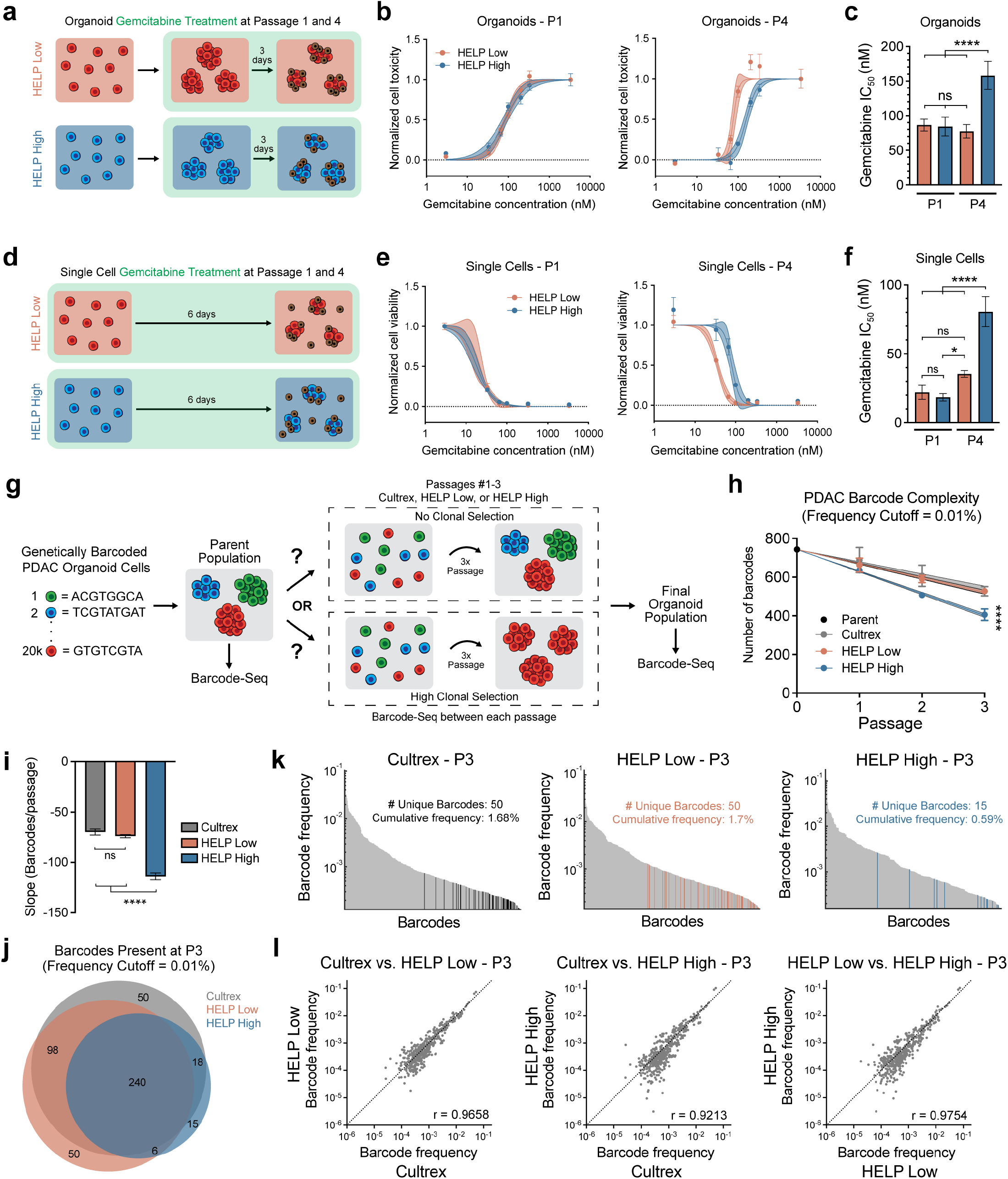
Prolonged exposure to high stiffness mediates PDAC organoid chemoresistance. **a**, Schematic of organoid drug treatment protocol, where PDAC cultures are treated with gemcitabine for three days following formation of ∼75-μm diameter multicellular organoids. **b**, Organoid-level gemcitabine dose-response curves for PDAC organoids expanded within HELP Low and HELP High matrices for either one (left) or four (right) passages. **c**, Gemcitabine IC_50_ values calculated from nonlinear fit of dose-response curves shown in **b. d**, Schematic of single cell drug treatment protocol, where PDAC cultures are treated with gemcitabine for six days during log-phase growth of single cells. **e**, Single cell-level gemcitabine dose-response curves for PDAC organoids expanded within HELP Low and HELP High matrices for either one (left) or four (right) passages. **f**, Gemcitabine IC_50_ values calculated from nonlinear fit of dose-response curves shown in **e**. In **b** and **e**, each data point represents the mean ± SEM (N=4, solid center line is nonlinear least squares regression of data; shaded region represents 95% confidence bands of nonlinear fit; data are normalized to positive controls (DMSO for single cells; 3333 nM gemcitabine for organoids). In **c** and **f**, each bar represents the mean ± SEM (N=3-4, ordinary one-way ANOVA with Tukey multiple comparisons correction, *p<0.05, ****p<0.0001, ns = not significant). **g**, Schematic summarizing protocol and potential outcomes of genetically barcoded PDAC organoid expansion within Cultrex and HELP matrices. **h**, Number of barcodes present within PDAC organoid populations expanded within Cultrex and HELP matrices for one to three passages (N=3, mean ± 95% confidence interval, center line represents linear fit for each matrix over three passages, shaded region represents 95% confidence interval of linear fit). Statistical analysis was performed using an ordinary one-way ANOVA with Tukey multiple comparisons correction between data points at passage three (****p<0.0001 for comparison between HELP High and Cultrex/HELP Low, comparison between HELP Low and Cultrex was not significantly different). The initial Parent population was the same for all matrices (N=1). **i**, Slope of linear fit from **h** for each matrix (N=3, mean ± 95% confidence interval, ordinary one-way ANOVA with Tukey multiple comparisons correction, ****p<0.0001, ns = not significant). **j**, Venn diagram of barcodes at passage three across indicated matrices. **k**, Frequencies of barcodes present in the indicated matrix at passage three. Barcodes are listed in rank order and colored bars correspond to barcodes unique to only the indicated matrix. The number of these unique barcodes and their cumulative frequency are reported for each matrix. **l**, Correlation plots comparing barcode frequencies across Cultrex and HELP at passage three. Pearson r values are reported for each pairing. Dashed line represents perfect correlation. In **j**-**l**, only barcodes with a frequency >0.01% across all three biological replicates at passage three were included.

Furthermore, to test whether drug response was dependent on the stage of organoid formation, we treated a subset of PDAC cells with gemcitabine during log-phase growth of single cells following either one or four passages (**Fig. 2d**). Following four passages in HELP High matrices, PDAC single cells adopted the same stiffness-mediated chemoresistance as seen with multicellular organoid drug treatment, while cells expanded in HELP Low or Cultrex matrices did not significantly increase their IC_50_ value (**Fig. 2e,f, Supplementary Fig. 7**). These results show that stiffness-mediated chemoresistance is not strictly an emergent phenotype of multicellular organoids, suggesting that PDAC is being influenced by matrix stiffness on the single cell-level.

### Prolonged exposure to high stiffness mediates PDAC clonal heterogeneity

PDAC tumors are characterized by their significant intratumor heterogeneity and the existence of clonal populations that may uniquely respond to drug treatments^34^. Given that organoid models can retain cellular heterogeneity *in vitro*^35–38^ and that our chemoresistant phenotype is preserved on the single cell-level, we hypothesized that stiffness-mediated chemoresistance could be due to the selection of inherently resistant subclones within our organoid population. To explore this hypothesis, we cultured genetically barcoded (via lentiviral transduction)^39^ PDAC organoids within Cultrex, HELP Low, or HELP High matrices for three passages (**Fig. 2g**). Following each passage, a subset of organoids was collected for barcode sequencing to track the relative frequencies of each clone over time. Our results show PDAC organoids cultured within HELP High matrices had steeper clonal selection over three passages compared to softer HELP Low and Cultrex matrices (**Fig. 2h,i, Supplementary Fig. 8a**). To mitigate noise, a barcode frequency cutoff of 0.01% was applied, which removed <2.5% of reads for each replicate (**Supplementary Fig. 8b**).

To identify whether the enriched clones in HELP High were unique across matrices, we quantified the overlap of barcodes and their relative frequencies following three passages. We found each matrix enriched a subset of barcodes unique to only that matrix (**Fig. 2j**) and that several clones preferred expanding in either a low or high stiffness environment, suggesting distinct cancer subclones exhibit different fitness in response to matrix properties (**Supplementary Fig. 9**,**10**).

Despite this preference for specific matrices by some subclones, the majority of subclone barcodes in HELP High were also present in HELP Low and Cultrex matrices (88% and 92% overlap, respectively, **Fig. 2j, Supplementary Fig. 11**). Moreover, the frequency of the 15 barcodes unique to HELP High were relatively low, and their cumulative frequency accounted for <1% of the total population (**Fig. 2k**), suggesting these clones have not sufficiently expanded to drive broad changes in gemcitabine IC_50_. In addition, the frequency of clones in HELP High was highly correlated with HELP Low and Cultrex matrices (**Fig. 2l**). Overall, these data suggest the HELP High matrices did not enrich for an inherently chemoresistant population and that PDAC organoids may instead be altering their phenotype in response to matrix stiffness.

### PDAC cancer stem cell phenotype is enriched in high stiffness matrices

CSCs are a population of malignant cells within a tumor that exhibit a de-differentiated phenotype and increased chemoresistance through a broad range of mechanisms^11^. In PDAC, CSCs are often identified by the expression of several cell-surface markers, including CD44, ABCG2, CD9, CD24, and CD133, each of which has been previously associated with poor prognosis in pancreatic cancer^6,40–44^. To corroborate the potential correlation between these CSC markers and PDAC patient survival, we evaluated data from The Cancer Genome Atlas and quantified overall survival for patients stratified by upper and lower quartile gene expression of each CSC marker (**Fig. 3a**)^45^. We found decreased survival was associated with increased marker expression in this cohort for several CSC markers. Notably, while *ABCG2* did not follow this trend in this cohort, *ABCG2* was highly upregulated in PDAC tumors compared to normal pancreatic tissue in the same dataset (**Supplementary Fig. 12**).

**Figure 3.**
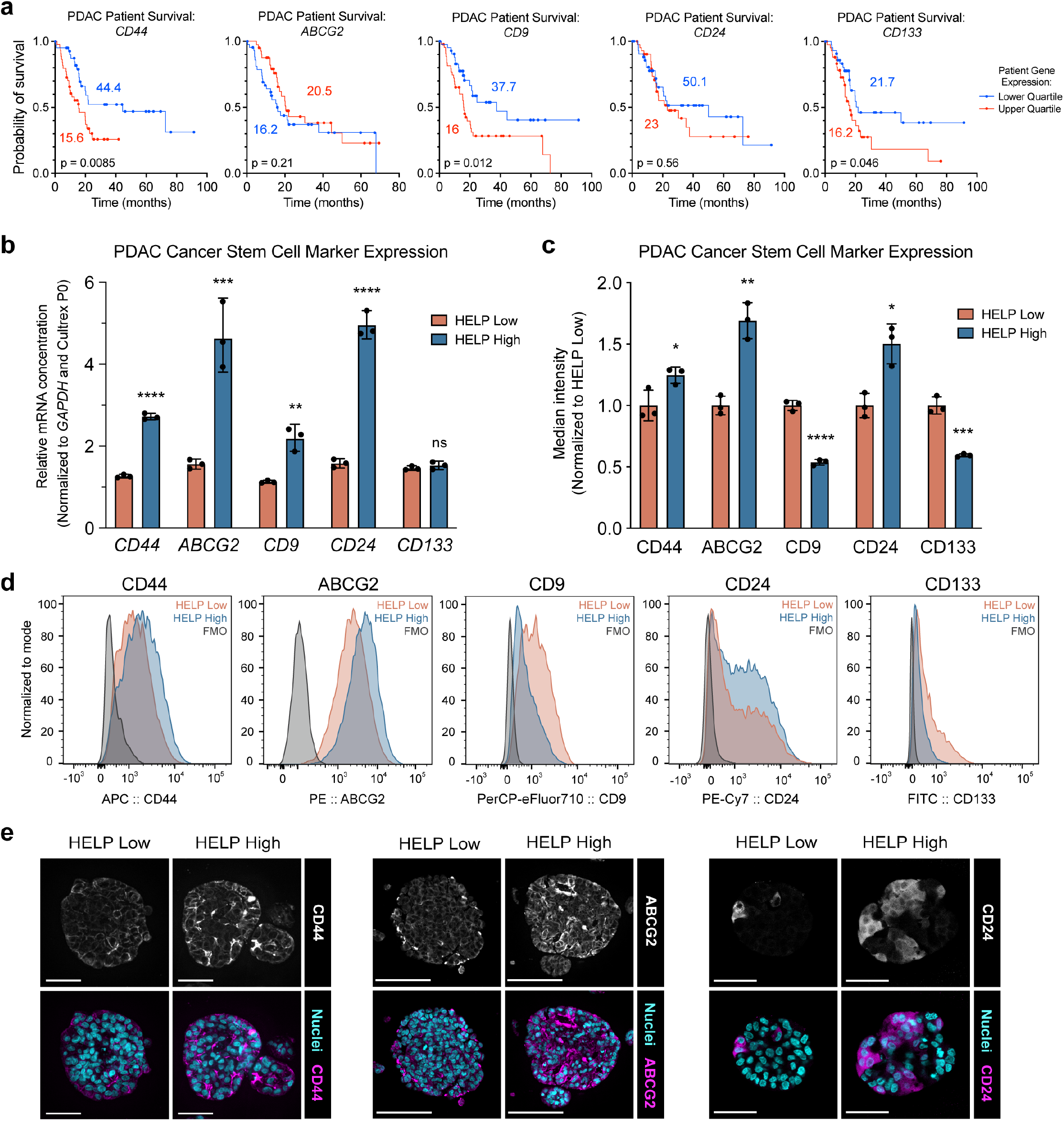
PDAC cancer stem cell phenotype is enriched in high stiffness matrices. **a**, Kaplan-Meier survival curves of PDAC patients in The Cancer Genome Atlas repository stratified by expression of the indicated CSC marker (lower quartile, blue, n=44; upper quartile, red, n=44; log-rank Mantel-Cox test). Filled circles represent censored events. **b**, qPCR quantification of mRNA-level CSC marker expression in PDAC organoids expanded within HELP Low or HELP High for four passages (N=3, mean ± 95% confidence interval). Data are normalized to *GAPDH* gene expression and respective marker expression in the PDAC organoid parent population cultured within Cultrex prior to expansion in HELP (i.e. Cultrex P0). **c**, Median intensity of CSC marker expression measured via flow cytometry of PDAC organoids expanded within HELP Low or HELP High for four passages (N=3, mean ± SD). Data are normalized to HELP Low. In **b** and **c**, statistical analysis was performed using an unpaired two-tailed Student’s t-test comparing HELP Low and HELP High for the indicated marker (*p<0.05, **p<0.01, ***p<0.001, ****p<0.0001, ns = not significant). **d**, Representative flow cytometry analysis of CSC marker expression in PDAC organoids expanded within HELP Low and HELP High (representative of N=3 replicates). Fluorescence minus one (FMO) controls are included for each marker (grey). **e**, Representative IF images of CD44 (left; scale bar, 50 μm), ABCG2 (middle; scale bar, 100 μm), and CD24 (right; scale bar, 50 μm) in PDAC organoids expanded within HELP Low and HELP High.

To explore the hypothesis that increased matrix stiffness was promoting a CSC-like phenotype within our *in vitro* model, PDAC organoids were expanded within Cultrex, HELP Low, or HELP High matrices for four passages. PDAC organoids cultured in HELP High matrices had increased mRNA expression of all tested CSC markers except *CD133*, compared to HELP Low and Cultrex (**Fig. 3b, Supplementary Fig. 13a**). Protein-level expression of CSC markers measured by flow cytometry and immunostaining showed a similar increase in overall expression of CD44, ABCG2, and CD24 in organoids grown within HELP High compared to HELP Low (**Fig. 3c-e, Supplementary Fig. 13b-d**).

### Drug efflux transporter expression mediates PDAC organoid chemoresistance

Altered expression of cell-surface drug transporters is a common mechanism utilized by CSCs to resist chemotherapy^46,47^. Therefore, understanding the environmental cues that mediate the development and maintenance of drug transporter expression could potentially lead to new treatment strategies to sensitize patient tumors. In particular, the ATP-binding cassette (ABC) family of cell-surface transporters are expressed across many tissues and cell types and are known for regulating efflux of a variety of endogenous and foreign substrates, including chemotherapeutics^48^. Due to the significant increase in ABCG2 transporter expression in HELP High matrices we observed while evaluating expression of CSC markers, we hypothesized the expression of other similar drug pumps may be upregulated in high stiffness matrices.

We found that several ABC-family drug efflux transporters commonly associated with PDAC and/or gemcitabine resistance^49–54^ (*ABCG2, ABCC3/4/5*) were upregulated in organoids in stiff HELP High matrices compared to softer HELP Low and Cultrex matrices (**Fig. 4a, Supplementary Fig. 14a**). Notably, solute carrier (SLC) family concentrative and equilibrative transporters that have been associated with gemcitabine influx^3^ (*SLC28A1, SLC28A3, SLC29A1, SLC29A2*) were either not influenced by matrix stiffness or showed a modest increase in HELP High (**Fig. 4b, Supplementary Fig. 14b**).

**Figure 4.**
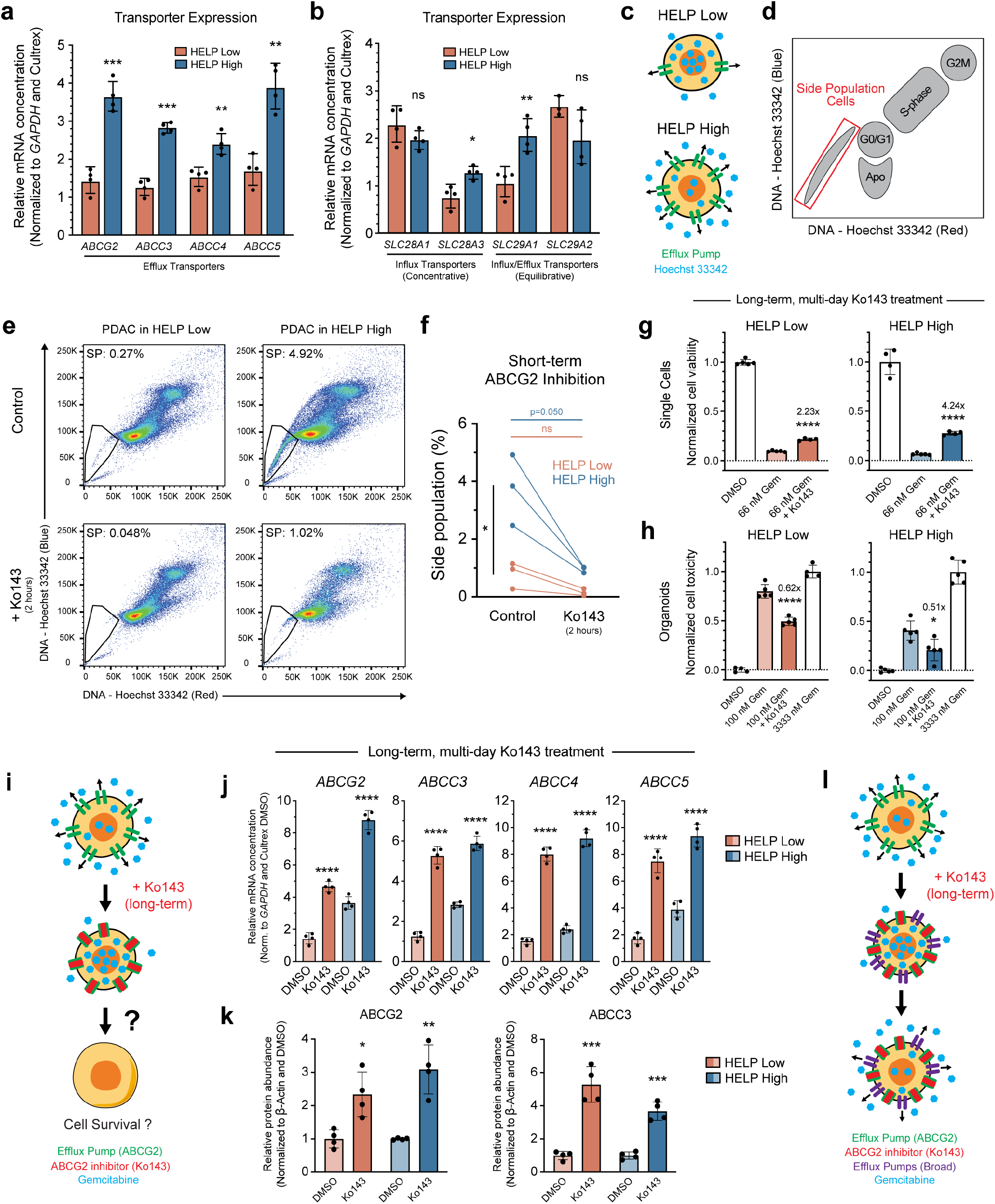
Drug efflux transporter expression mediates PDAC organoid chemoresistance. **a**,**b**, qPCR quantification of mRNA-level ATP-binding cassette (ABC) family (**a**) and solute carrier (SLC) family (**b**) drug transporter expression in PDAC organoids expanded within HELP Low or HELP High for four passages (N=4, mean ± 95% confidence interval, two-tailed Student’s t-test comparing HELP Low and HELP High, *p<0.05, **p<0.01, ***p<0.001, ns = not significant). Data are normalized to *GAPDH* gene expression and respective marker expression in PDAC organoids expanded within Cultrex. **c**, Illustrated summary of drug transporter expression in HELP Low or HELP High matrices and predicted efflux of Hoechst fluorescent dye. **d**, Schematic of side population (SP) results via flow cytometry analysis. Cells with higher efflux transporter activity (i.e. SP, outlined in red) have decreased Hoechst (DNA) signal. Schematic adapted from Petriz^59^. **e**, Representative SP analysis via flow cytometry of PDAC organoids expanded within HELP Low (left) or HELP High (right) for four passages and treated with Hoechst (top) or Hoechst + Ko143 (ABCG2 efflux transporter inhibitor; bottom) for two hours (representative of N=3 replicates). SP is outlined in black and percentage of cells in SP is reported for each sample. **f**, Quantification of SP cells from the full dataset of flow cytometry data represented in **e** (N=3). Data points represent one biological replicate and data points connected by a line are from the same parent population of cells. Statistical analysis comparing control vs. Ko143 for each matrix was performed using a paired t-test (color-matched statistical bars, p=0.050, ns = not significant). Statistical analysis comparing percentage of SP cells in control samples across matrices was performed using a two-tailed Student’s t-test (black statistical bars, *p<0.05). **g**, PDAC viability following treatment with DMSO (control, normalized to 1), 66 nM gemcitabine (Gem), or 66 nM gemcitabine + 20 μM Ko143 (N=4-5, mean ± SD). **h**, PDAC toxicity following treatment with DMSO (control, normalized to 0), 100 nM gemcitabine, 100 nM gemcitabine + 20 μM Ko143, or 3333 nM gemcitabine (positive control, normalized to 1) (N=4-5, mean ± SD). In **g** and **h**, PDAC organoids were expanded for four passages in HELP Low (left) or HELP High (right) prior to gemcitabine (+ Ko143) treatment for six days on single cells during log-phase growth (**g**) or for three days following formation of ∼75-μm diameter multicellular organoids (**h**). Statistical analysis comparing experimental gemcitabine treatment and gemcitabine + Ko143 treatment was performed using an unpaired two-tailed Student’s t-test (*p<0.05, ****p<0.0001), and the fold change between these two conditions is reported for each comparison. **i**, Illustrated summary of results from **g** and **h. j**,**k**, qPCR (**j**) and Western blot (**k**) quantification of mRNA and protein-level ABC-family drug efflux transporter expression in PDAC organoids expanded within HELP Low or HELP High for four passages and treated with either DMSO or 20 μM Ko143 (N=4, qPCR: mean ± 95% confidence interval, Western: mean ± SD). Statistical analysis comparing DMSO vs. Ko143 treatment for each matrix was performed using an unpaired two-tailed Student’s t-test (*p<0.05, **p<0.01, ***p<0.001, ****p<0.0001). qPCR data are normalized to *GAPDH* gene expression and respective marker expression of Cultrex DMSO samples. Western data are normalized to β-actin expression and DMSO samples for each marker and matrix. PDAC cells were treated with Ko143 throughout single cell log-phase growth for six days. **l**, Illustrated summary of results from **j** and **k**.

To measure the functional activity of drug efflux transporters, we performed a side population (SP) assay via flow cytometry, which is used to identify healthy stem cell and CSC populations across several tissue types^55^. Following expansion of PDAC organoids for four passages in HELP Low, HELP High, or Cultrex matrices, organoids were extracted from the hydrogels, dissociated into single cells, and treated with Hoechst 33342, a live cell-permeable, fluorescent DNA stain. As several ABC-family transporters, including ABCG2, have been shown to readily efflux Hoechst, cells with higher efflux pump activity should show decreased fluorescent signal when measured via flow cytometry^55,56^ (**Fig. 4c,d**).

In agreement with our mRNA expression data, we found organoids cultured in HELP High had a significantly larger SP (3.74% ± 1.23%, mean ± SD) compared to HELP Low (0.79% ± 0.46%) (**Fig. 4e,f, Supplementary Fig. 15**,**16**). To test the specific role of ABCG2 in mediating this response, we performed the same SP assay with the addition of Ko143, a small-molecule inhibitor with strong affinity to the ABCG2 transporter^57,58^. Treatment with Ko143 for two hours led to a significant decrease in SP across organoids cultured in both HELP High (0.95% ± 0.11%) and Low (0.15% ± 0.12%) matrices (**Fig. 4e,f, Supplementary Fig. 15**,**16**), highlighting ABCG2 as a highly active efflux transporter for PDAC organoids. Similar treatment with verapamil, a less-specific ABC-family transport inhibitor^51,55^, also led to a decrease in SP (**Supplementary Fig. 15-18**). Interestingly, organoids cultured within Cultrex had a relatively high percentage of SP cells and a similar decrease in SP following treatment with Ko143 and verapamil (**Supplementary Fig. 17**,**18**), suggesting other uncontrolled matrix properties or soluble cues within animal-derived Cultrex hydrogels may also be contributing to overall efflux activity^22,26^.

Due to their ability to block drug efflux pumps that may enable chemoresistance in cancer cells, inhibitors like Ko143 and verapamil have been explored as a potential treatment strategy to sensitize tumors to chemotherapy^46,48,58^. Therefore, we hypothesized that co-administration of gemcitabine and Ko143 *in vitro* would lead to increased drug sensitivity of PDAC organoids and overall reduced cell survival.

Surprisingly, we found long-term addition of Ko143 throughout our 6-day single cell and 3-day organoid drug treatment protocols decreased sensitivity to gemcitabine and promoted PDAC survival across HELP matrices, with a stronger effect seen in HELP High (**Fig. 4g-i, Supplementary Fig. 19a**). In agreement with SP data, organoids within Cultrex showed a similar response to long-term Ko143 treatment as those expanded within HELP High (**Supplementary Fig. 19b**,**c**). Importantly, Ko143 treatment alone did not increase cell viability compared to DMSO control (**Supplementary Fig. 19d**).

To explore how long-term Ko143 treatment resulted in increased PDAC organoid chemoresistance, we measured the expression of several drug transporters. Interestingly, we found the addition of Ko143 alone throughout organoid formation led to broad mRNA- and protein-level upregulation of efflux transporters compared to DMSO controls (**Fig. 4j,k, Supplementary Fig. 20a-e**). Similar results were seen for SLC-family concentrative and equilibrative transporters, albeit to a lesser extent than ABC-family efflux pumps (**Supplementary Fig. 20f**). Overall, our data suggest PDAC cells compensate for efflux pump inhibition by driving the expression of broad efflux transporters, resulting in increased chemoresistance (**Fig. 4l**).

### Hyaluronan mechanosignaling mediates PDAC chemoresistance

We have so far described how PDAC organoids dynamically adapt their phenotype in response to increased matrix stiffness by increasing expression of ABCG2, leading to decreased chemosensitivity. To further elucidate the mechanism of this mechanosignaling, we explored the role of specific matrix ligands in driving PDAC organoid chemoresistance by leveraging the tunability of our engineered matrix.

The PDAC ECM is a dynamic milieu comprised of several polymer components including fibronectin and HA^60^, which mediate cancer cell-ECM interactions through integrin and CD44 signaling, respectively. Furthermore, integrins and CD44 are known to transduce mechanosignaling^61,62^. In particular, the fibronectin mimicking RGD ligand has been implicated in mediating both cancer progression and drug sensitivity in several cancer types^63^. To explore the role of RGD signaling in our system, we modified the amino acid sequence of our recombinant ELP component to present a scrambled, non-integrin-binding R<u>DG</u> motif^28,33^, resulting in identical HELP matrices with 0 mM RGD (**Fig. 5a**). We found PDAC organoids grew robustly from single cell encapsulation in HELP R<u>DG</u> Low and HELP R<u>DG</u> High matrices without the RGD ligand (**Fig. 5b**). Several CSC markers, including *CD44* and *ABCG2*, were similarly upregulated in HELP R<u>DG</u> High compared to HELP R<u>DG</u> Low after four passages (**Fig. 5c, Supplementary Fig. 21a**).

**Figure 5.**
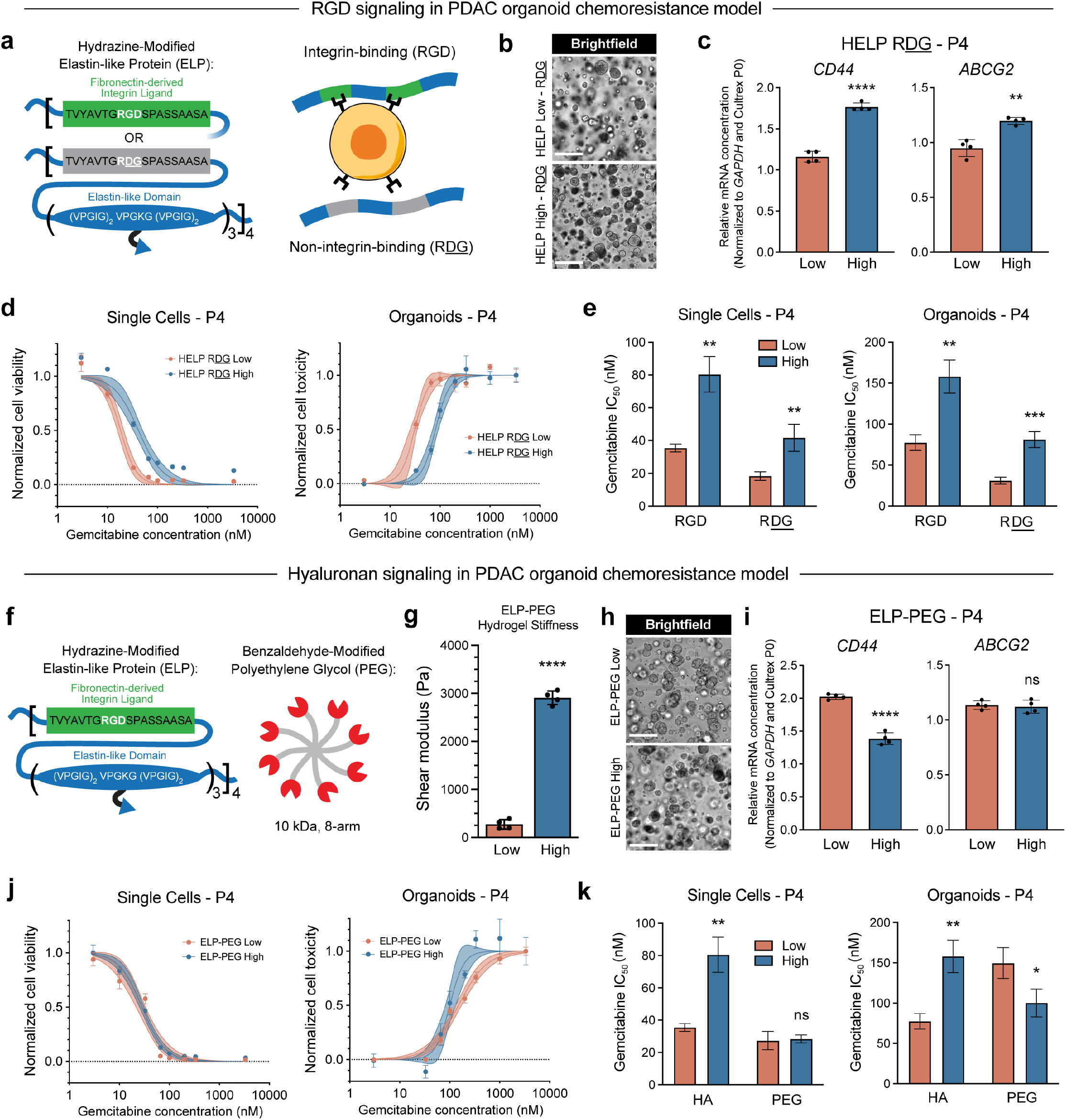
Influence of RGD and hyaluronan mechanosignaling on PDAC chemoresistance. **a**, Schematic of recombinant elastin-like protein (ELP) component of HELP, which can be engineered to display a non-interactive, scrambled R<u>DG</u> sequence, resulting in a HELP matrix without RGD, but with identical mechanical properties and HA concentration. **b**, Representative BF images of PDAC organoids expanded within HELP R<u>DG</u> Low (top) or HELP R<u>DG</u> High (bottom) for four passages. Scale bar, 250 μm. **c**, qPCR quantification of mRNA-level CSC marker expression in PDAC organoids expanded within HELP R<u>DG</u> Low or High for four passages (N=4, mean ± 95% confidence interval). **d**, Single cell-(left) and organoid-level (right) gemcitabine dose-response curves for PDAC organoids expanded within HELP R<u>DG</u> Low or High for four passages. **e**, Gemcitabine IC_50_ values calculated from nonlinear fit of dose-response curves shown in **d** for single cell (left) and organoid (right) drug treatment in HELP R<u>DG</u> compared to HELP containing RGD. **f**, Schematic of ELP-polyethylene glycol (PEG) matrix, where HA has been replaced with inert PEG, resulting in a HELP matrix without HA, but with identical mechanical properties and RGD concentration. **g**, Stiffness measurements of ELP-PEG matrices stiffness-matched to HELP Low and HELP High (N=4, mean ± SD, unpaired two-tailed Student’s t-test, ****p<0.0001). **h**, Representative BF images of PDAC organoids expanded within ELP-PEG Low (top) or ELP-PEG High (bottom) for four passages. Scale bar, 250 μm. **i**, qPCR quantification of mRNA-level CSC marker expression in PDAC organoids expanded within ELP-PEG Low or High for four passages (N=4, mean ± 95% confidence interval). **j**, Single cell-(left) and organoid-level (right) gemcitabine dose-response curves for PDAC organoids expanded within ELP-PEG Low or High for four passages. **k**, Gemcitabine IC_50_ values calculated from nonlinear fit of dose-response curves shown in **j** for single cell (left) and organoid (right) drug treatment in ELP-PEG compared to HELP containing HA. In **c** and **i**, statistical analysis comparing marker expression in Low vs. High matrices was performed using an unpaired two-tailed Student’s t-test (**p<0.01, ****p<0.0001, ns = not significant). All data are normalized to *GAPDH* gene expression and respective marker expression in the PDAC organoid parent population cultured within Cultrex prior to expansion in HELP R<u>DG</u> or ELP-PEG (i.e. Cultrex P0). In **d** and **j**, each data point represents the mean ± SEM (N=4, solid center line is nonlinear least squares regression of data; shaded region represents 95% confidence bands of nonlinear fit; data are normalized to positive controls (DMSO for single cells, 3333 nM gemcitabine for organoids). In **e** and **k**, each bar represents the mean ± SEM (N=4, unpaired two-tailed Student’s t-test between Low and High for each matrix variation, *p<0.05, **p<0.01 ***p<0.001, ns = not significant).

Furthermore, while HELP R<u>DG</u> High still promoted a chemoresistant phenotype compared to HELP R<u>DG</u> Low, the overall gemcitabine IC_50_ value decreased across both matrices compared to HELP matrices with RGD (**Fig. 5d,e**). These results reveal that RGD signaling was not necessary for organoid expansion nor the establishment of increased chemoresistance in high stiffness matrices, suggesting that RGD-integrin interactions are not the primary mechanism of stiffness-mediated PDAC chemoresistance.

CD44-mediated HA signaling has also been linked to altered drug response in other cancer types^64^. To explore the effect of HA on PDAC organoids in our model, we modified our HELP matrix by replacing HA with an inert 8-arm polyethylene glycol (PEG) polymer to fabricate matrices we term ELP-PEG that are stiffness-matched to HELP Low and High (**Fig. 5f,g, Supplementary Fig. 22**). We found PDAC organoids expanded robustly over four passages in ELP-PEG Low and High matrices despite removal of HA (**Fig. 5h**).

Interestingly, upregulation of several CSC makers, including *CD44* and *ABCG2*, was lost in PDAC organoids expanded within ELP-PEG High compared to ELP-PEG Low matrices following four passages (**Fig. 5i, Supplementary Fig. 21b**). Similarly, the stiffness-mediated chemoresistant phenotype was lost within ELP-PEG High compared to ELP-PEG Low matrices, unlike the increased chemoresistance measured in HELP High matrices with HA (**Fig. 5j,k**). Taken together, these data suggest that HA mechanosignaling plays a significant mechanistic role in PDAC CSC enrichment and acquired chemoresistance via ABCG2 upregulation in our model.

### Stiffness-mediated PDAC chemoresistance is reversible

Recent work has shown degradation of HA in the PDAC tumor microenvironment can improve survival in mouse models, with the hypothesis that it leads to increased drug delivery to the tumor^12^. Here, we identified that PDAC organoids can dynamically alter their phenotype and drug sensitivity in response to HA-mechanosignaling in a high stiffness matrix, even when drug transport through high and low stiffness matrices is similar. This stiffness-induced chemoresistance led us to hypothesize that reversing matrix stiffness may re-sensitize PDAC organoids to gemcitabine treatment.

To test this hypothesis, we expanded PDAC organoids in HELP Low or HELP High for a total of seven passages and performed drug treatment during log-phase growth of single cells or following organoid formation on a subset of cells at passages one, four, and seven. On the fourth passage, a subset of cells from each matrix were switched to the opposite stiffness matrix for the remaining three passages (**Fig. 6a**). Interestingly, in the passage immediately following the switch from one matrix to another, PDAC single cells and organoids retained short-term mechanical memory of their initial matrix stiffness (**Fig. 6b,c, Supplementary Fig. 23a**). For example, despite being cultured within a softer HELP Low matrix after the switch at passage four, PDAC organoids exhibited a similar gemcitabine IC_50_ value as organoids continuously maintained within HELP High. However, this mechanical memory was lost following an additional three passages within the opposite stiffness matrix (**Fig. 6b-e**). Over-expression of CSC markers, including *CD44* and *ABCG2*, was also reversible following several passages within the opposite stiffness matrix (**Fig. 6f, Supplementary Fig. 23b**).

**Figure 6.**
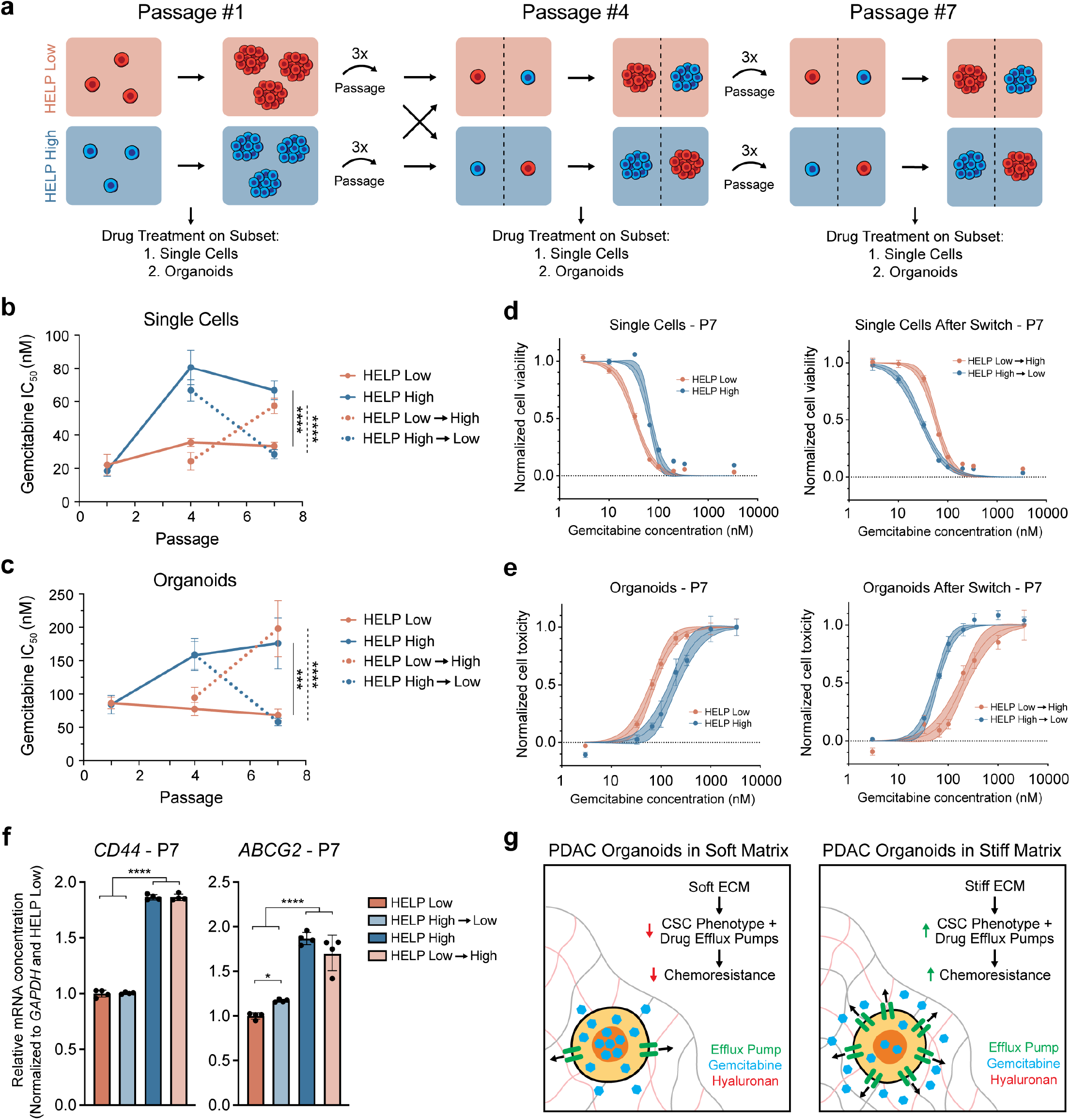
Stiffness-mediated PDAC chemoresistance is reversible. **a**, Schematic of PDAC organoid expansion protocol for testing reversibility of stiffness-mediated chemoresistance. **b**,**c**, Summary of gemcitabine IC_50_ values for PDAC organoids expanded within HELP Low or HELP High matrices according to the protocol in **a**. PDAC cells were treated with gemcitabine throughout log-phase single cell expansion (**b**) and following multicellular organoid formation (**c**) for six and three days, respectively. Each data point represents the mean ± 95% confidence interval (N=4). Statistical analysis comparing IC_50_ values at passage 7 for all samples was performed using an ordinary one-way ANOVA with Tukey multiple comparisons correction (***p<0.001, ****p<0.0001, solid line compares non-switched HELP matrices, dashed line compares switched HELP matrices). **d**,**e**, Single cell-(**d**) and organoid-level (**e**) gemcitabine dose-response curves for PDAC organoids expanded within HELP Low or HELP High for seven passages without (left) or with (right) a switch to the opposite matrix stiffness at passage four. Each data point represents the mean ± SEM (N=4, solid center line is nonlinear least squares regression of data, shaded region represents 95% confidence bands of nonlinear fit, data are normalized to positive controls (DMSO for single cells, 3333 nM gemcitabine for organoids). **f**, qPCR quantification of mRNA-level CSC marker expression in PDAC organoids expanded within HELP Low or HELP High for seven passages, with or without a switch to the opposite matrix stiffness at passage four (N=4, mean ± 95% confidence interval, ordinary one-way ANOVA with Tukey multiple comparisons correction, *p<0.05, ****p<0.0001, unlabeled comparisons are not significantly different). All data are normalized to *GAPDH* gene expression and respective marker expression in HELP Low at passage seven. **g**, Illustration summarizing mechanism of stiffness-mediated chemoresistance in this PDAC organoid model.

## Discussion

Our data show that patient-derived PDAC organoids dynamically and reversibly respond to increased matrix stiffness through HA mechanosignaling by adopting a CSC-like phenotype resulting in increased chemoresistance via over-expression of drug efflux transporters (**Fig. 6g**). Importantly, these findings would not have been possible with heterogeneous, non-tunable animal-derived matrices (e.g. Cultrex/Matrigel), highlighting the need for tunable engineered matrices to uncover the role of ECM properties in driving human cancer organoid phenotype and drug response.

We demonstrated that PDAC organoids reverse their phenotype and become sensitive to gemcitabine treatment when transferred from high to low matrix stiffness. These results suggest treatment of PDAC tumors with mechanotherapeutics that target matrix stiffness and/or cell-ECM interactions in combination with anti-cancer agents may improve tumor drug sensitivity and lead to improved patient outcomes. Notably, several mechanotherapeutics targeting cell-ECM signaling pathways are currently being explored in pancreatic cancer^65^, and in the context of our work, highlight the importance of considering the influence of ECM signaling when developing anti-cancer treatments. Future work should also explore how the 3D ECM influences resistance to other chemotherapies and mediates the epithelial-mesenchymal transition in PDAC, as shown in previous 2D models^66^.

Although stiffness-mediated chemoresistance was not a result of clonal enrichment of chemoresistant populations, our single cell barcoding experiments showed that individual PDAC clones exhibit unique fitness when cultured in soft vs. stiff matrices. This result highlights that matrix properties can drive selection of cancer subclones, which should be considered when culturing patient-derived organoids in any model system and warrants future investigation.

As inhibitors of drug efflux transporters have clinical potential to sensitize tumors to chemotherapies, we hypothesized treatment with drug transporter inhibitors in our organoid model would lead to decreased cell survival. Instead, we found long-term exposure to Ko143 led to increased cancer cell survival and chemoresistance, mediated through broad upregulation of drug efflux transporters. To our knowledge this is the first report identifying the extreme plasticity of drug efflux pump expression in response to a blocking agent and suggests a potential mechanism for why these inhibitors have been ineffective in sensitizing PDAC tumors in the clinic^48^. Therefore, future strategies for sensitizing chemoresistant PDAC tumors may require combination therapies that target the source of altered transporter expression, such as the ECM properties identified here.

## Materials and Methods

### Material synthesis and characterization

#### Elastin-like protein expression

Elastin-like protein (ELP) was produced as previously reported^33^. Briefly, BL21(DE3)pLysS *Escherichia coli* (Invitrogen C606003) were transformed with a pET15b plasmid encoding the ELP sequence containing either the RGD or R<u>DG</u> motif. The bacteria were cultured to an OD_600_ of 0.8 and ELP expression was induced by 1 mM isopropyl β-D-1-thiogalactopyranoside (IPTG; Thermo Fisher BP1755) under the control of a T7 promoter. Following a 7-hour incubation at 32 °C, the bacteria were pelleted and lysed through three freeze-thaw cycles in the presence of TEN buffer (10 mM Tris, 100 mM NaCl, 1 mM ethylenediaminetetraacetic acid (EDTA), pH 8.0) supplemented with 10 µM DNAse I (Sigma DN25) and 1 mM phenylmethanesulfonyl fluoride (PMSF; MP Biomedicals 195381) protease inhibitor. ELPs were purified from cell lysate by successive thermal cycling and centrifugation steps followed by dialysis against Milli-Q water for 3 days at 4 °C with regular water changes (molecular weight cutoff: 10,000 Da; Spectrum 132576). The final ELP product was lyophilized and stored at -20 °C. Typical yields from 12 L of bacterial culture are between 300-1000 mg. ELP purity and molecular weight were confirmed using Western blot. See Supplementary Table 2 for full ELP amino acid sequences.

#### Synthesis of hydrazine-modified elastin-like protein

ELP was modified with hydrazine as previously reported^28^ and produced in batches ranging from 300-1000 mg. The following protocol is written for a 1 g batch and was scaled as appropriate. Lyophilized ELP was added to a 100 mL round bottom flask and fully dissolved in 13.66 mL anhydrous dimethyl sulfoxide (DMSO; Sigma 276855) to a concentration of 7.3 wt% at room temperature (RT). Once dissolved, an equal volume of anhydrous dimethylformamide (DMF; Sigma 227056) was added to the same reaction vessel. In a separate 20 mL glass vial, tri-Boc hydrazinoacetic acid (303 mg, 0.775 mmol, 2.1 eq:ELP amines; Sigma 68972) and hexafluorophosphate azabenzotriazole tetramethyl uronium (HATU; 281 mg, 0.738 mmol, 2 eq:ELP amines; Sigma 445460) were dissolved in 13.66 mL DMF. Immediately after, 4-methylmorpholine (203 uL, 0.92 g/mL, 1.8 mmol, 5 eq: ELP amines; Sigma M56557) was added to the vial and allowed to stir for 10 min at RT. After 10 min, the activated tri-Boc hydrazinoacetic acid mixture was added to the ELP reaction vessel and allowed to react for 24 hours at RT. The following day, the Boc-protected ELP-hydrazine product was precipitated by addition to ice-cold diethyl ether (∼120 mL; Fisher E138), collected via centrifugation at 18,000 x g for 25 min, and dried under a stream of nitrogen gas. To calculate the degree of boc-hydrazinoacetic acid functionalization, the modified ELP polymers were dissolved in DMSO-d6 (Thermo Fisher 320770075) at 10 mg/mL for proton nuclear magnetic resonance spectroscopy (^1^H NMR; 500 MHz, Varian Inova). The degree of hydrazine modification was estimated using the signal from the Boc groups. The aromatic protons of the tyrosine residues in the amino acid sequence of ELP served as a reference signal and the modification calculation is summarized in Supplementary Fig. 1. ^1^H NMR (500 MHz, DMSO-d6) *δ* ppm 7.00 and 6.62 (2H each, d, tyrosine amino acid); 1.46 and 1.39 (27H, Boc groups). The Boc protecting groups were removed via acid-mediated deprotection by dissolving the modified ELPs to a final concentration of 3.3 wt% in 30 mL of a 1:1 mixture of dichloromethane (DCM; Sigma DX0835-3) and trifluoroacetic acid (TFA; Sigma T6508) supplemented with 5% v/v tri-isopropylsilane (Sigma 233781). After stirring for 4 hours at RT, deprotected ELP-hydrazine was precipitated by addition to ice-cold diethyl ether (∼90 mL), dried under a stream of nitrogen gas, and dissolved in ice-cold Milli-Q water overnight. The solution was dialyzed against Milli-Q water for 3 days at 4 °C with regular water changes (molecular weight cutoff: 10,000 Da), sterile-filtered, and lyophilized to afford the product as a white solid (∼700 mg, ∼70% yield), which was stored at -20 °C. ^1^H NMR was used as above to ensure removal of Boc protecting groups.

#### Synthesis of benzaldehyde-modified hyaluronan

The synthesis of hyaluronan modified with a benzaldehyde (HA-BZA) crosslinking group can be divided into two main steps. First, the carboxylic acid groups on hyaluronan are amidated with propargylamine via a 1-ethyl-3-(3-dimethylaminopropyl)carbodiimide hydrochloride (EDC) carbodiimide crosslinking reaction to form an HA-alkyne intermediate polymer. Second, using copper-catalyzed azide-alkyne click chemistry, we modify the alkyne groups on the HA with a small molecule azidobenzaldehyde to form the final HA-BZA polymer. To fabricate an engineered matrix with varying stiffness, we created two versions of HA with a low (∼7%) or high (∼34%) modification of alkyne/BZA, represented as a percentage of the total available carboxylic groups on the HA polymer.

HA-alkyne was prepared in batches of 200-1000 mg. Linear HA (100 kDa, sodium salt, LifeCore Biomedical HA100K) was dissolved in 2-(N-Morpholino)ethanesulfonic acid (MES) buffer (0.2 M MES hydrate (Sigma M2933), 0.15 M NaCl in Milli-Q water; pH 4.5) to a concentration of 1 wt% in a round bottom flask. Once the HA had fully dissolved, propargylamine (0.8 or 6.0 equiv:HA carboxylic acid groups for 7% or 34% modification, respectively; Sigma P50900) was added to the reaction vessel and the pH was immediately adjusted to 6 using NaOH. N-hydroxysuccinimide (NHS; 0.8 or 6.0 eq:HA carboxylic acid groups for 7% or 34% modification, respectively; Thermo Fisher 24500) and EDC (0.8 or 6.0 eq:HA carboxylic acid groups for 7% or 34% modification, respectively; Thermo Fisher 22980) were added sequentially to the reaction vessel. The reaction was allowed to stir at RT for 4 or 24 hours for 7% or 34% modification, respectively. The final solution was dialyzed against Milli-Q water for 3 days at 4 °C with regular water changes (molecular weight cutoff: 10,000 Da), sterile-filtered, and lyophilized to afford the product (∼900 mg, ∼90% yield), which was stored at -20 °C.

The small molecule precursor azidobenzaldehyde was synthesized and analyzed as described previously^67^ with slight modification. Specifically, the carboxylic acid group on 4-formylbenzoic acid is amidated with a 3-azidopropylamine via an EDC/NHS carbodiimide crosslinking reaction. Azidobenzaldehyde was prepared in 1-2 g batches. The following protocol was written for a 1 g batch and scaled as appropriate. First, 4-formylbenzoic acid (6.66 mmol, 1.0 eq; Sigma 124915) was dissolved in 30 mL extra dry DCM (Thermo Fisher 610300010) in a round bottom reaction vessel. NHS (0.843 g, 7.33 mmol, 1.1 eq) and EDC (1.41 g, 7.33 mmol, 1.1 eq) were added sequentially to the reaction vessel. The reaction was covered and allowed to stir at RT for 24 hours. Next, the reaction was diluted in 30 mL DCM (Thermo Fisher D151-4) and washed with water (2 × 60 mL) and brine (2 × 60 mL). The organic layer was dried over magnesium sulfate, filtered, and concentrated via rotary evaporation to afford the intermediate, NHS-benzaldehyde, as a white powder (975 mg, 3.94 mmol, 97.5% yield), which was subsequently stored at -20 °C. Next, the NHS-benzaldehyde intermediate (1 g, 1.0 eq) was dissolved in 15 mL dry DCM in a round bottom flask. 3-azidopropylamine (608 mg, 1.5 eq; Click Chemistry Tools AZ115) and diisopropyl ethyl amine (DIPEA; 2.12 mL, 3.0 eq; Sigma D125806) were added sequentially to the reaction vessel. The reaction was capped, protected from light, and allowed to stir for 24 hours at RT. The reaction mixture was diluted in 15 mL DCM and washed with brine (2 × 60 mL). The organic layer was dried over magnesium sulfate, filtered, and concentrated via rotary evaporation to a minimal volume of DCM such that the product remained in solution. The final azidobenzaldehyde product was purified via silica flash chromatography, eluting with a 10:1 DCM:acetone mobile phase into ∼15 mL fractions. The fractions containing the azidobenzaldehyde product were identified via TLC, collected into a single vessel, and concentrated via rotary evaporation to afford a light-yellow oil that solidified to a white/yellow powder upon standing at -20 °C (475 mg, 2.04 mmol, 47.5% yield). The product was subsequently stored at -20 °C for long-term storage. ^1^H NMR (500 MHz, CDCl3) *δ* ppm 10.09 (s, 1H, aldehyde), 7.97 (d, J = 8.5 Hz, 2H, aromatic), 7.93 (d, J = 8.3 Hz, 2H, aromatic), 6.50 (br. s, 1H, amide), 3.60 (q, J = 6.5 Hz, 2H, propyl), 3.49 (t, J = 6.3 Hz, 2H, propyl), 1.95 (quin, J = 6.5 Hz, 2H, propyl).

HA-BZA was prepared in batches of 200-1000 mg. Lyophilized HA-alkyne was dissolved to a final concentration of 1 wt% in a 10x phosphate buffered saline solution (10x PBS; 81 mM sodium phosphate dibasic, 19 mM sodium phosphate monobasic, 60 mM sodium chloride in Milli-Q water; pH 7.4) supplemented with 1 mg/mL beta-cyclodextrin (Sigma C4767). Once the HA-alkyne had fully dissolved, the solution was degassed by bubbling with nitrogen (N_2_) for 30 min. Next, N_2_ degassed solutions of sodium ascorbate (4.52 mM, 0.18 eq:HA carboxylic acid groups, Sigma A7631) and copper (II) sulfate pentahydrate (0.24 mM, 0.0096 eq:HA carboxylic acid groups, Sigma 209198) dissolved in Milli-Q water were sequentially added to the HA-alkyne reaction vessel via a syringe. Finally, azidobenzaldehyde (0.6 or 2.0 eq:alkyne groups for 7% or 34% modification, respectively) dissolved in a minimal amount of anhydrous DMSO (∼300 mg/mL; Sigma 276855) was added to the reaction vessel and the final solution was degassed for an additional 15 min. Following 24 hours of stirring at RT, an equal volume of 50 mM EDTA in Milli-Q water (pH 7.0; Thermo Fisher O2793) was added to the reaction to chelate remaining copper for 1 hour. The final solution was dialyzed against Milli-Q water for 3 days at 4 °C with regular water changes (molecular weight cutoff: 10,000 Da), sterile-filtered, and lyophilized to afford the product (∼750 mg, ∼75% yield), which was stored at -20 °C. ^1^H NMR (500 MHz, Varian Inova) was used to calculate degree of modification of BZA onto the HA-alkyne polymer by measuring the protons on the benzene ring, triazole linkage, and aldehyde group. The acetyl group on the HA was used as a reference. HA-BZA was dissolved in deuterated water (D_2_O; Thermo Fisher 351430075) at 10 mg/mL for NMR analysis. ^1^H NMR (500 MHz, D_2_O) *δ* ppm 9.9 (1H, s, aldehyde); 7.93 and 7.82 (2H each; d; benzene ring); 7.9 (1H, s, triazole link); 1.8 (3H, HA acetyl group, reference).

#### Synthesis of benzaldehyde-modified polyethylene glycol

Polyethylene glycol (PEG)-alkyne (8-arm, 10 kDa; Creative PEGWorks PSB-886) was modified with a BZA group using the same copper click reaction protocol for HA-BZA with the following alterations. PEG-BZA was produced in 600 mg (480 µmol alkyne) batches. The small molecule azidobenzaldehyde (960 µmol, 2.0 eq:alkyne groups) was dissolved in a minimal amount of anhydrous DMSO (∼300 mg/mL) and added to the reaction. The final product was dialyzed against Milli-Q water for 3 days at 4 °C with regular water changes (molecular weight cutoff: 3500 Da; Spectrum 132592), sterile-filtered, and lyophilized to afford the product (500 mg, 83% yield), which was stored at -20 °C. ^1^H NMR (500 MHz, Varian Inova) was used calculate degree of modification of BZA onto the PEG polymer. PEG-BZA was dissolved in D_2_O at 10 mg/mL for NMR analysis. ^1^H NMR (500 MHz, D_2_O) *δ* ppm 9.9 (1H, s, aldehyde); 7.93 and 7.82 (2H each; d; benzene ring); 7.9 (1H, s, triazole link); 3.56 (105H per arm; PEG monomer).

#### HELP, ELP-PEG, and Cultrex matrix formation

For all experiments, HELP matrices were prepared by dissolving lyophilized HA and ELP components in a 10x phosphate buffered saline solution (10x PBS; 81 mM sodium phosphate dibasic, 19 mM sodium phosphate monobasic, 60 mM sodium chloride in Milli-Q water; pH 7.4) to a concentration double their final hydrogel concentration, allowing the formation of matrices upon a 1:1 mixture of stock HA-benzaldehyde and ELP-hydrazine solutions. Specifically, the appropriate volume of 2x stock HA solution was added to the corresponding well/mold and an equal volume of 2x stock ELP solution was pipetted directly onto the HA and immediately mixed using the same pipette tip. Both the HA and ELP solutions and the well-plate containing the hydrogels was kept on ice during the mixing process. Following mixing, the matrices were immediately incubated at RT for 10 mins (HELP Low and Medium) or 5 mins (HELP High), followed by another incubation at 37 °C for 10 mins (HELP Low and Medium) or 5 mins (HELP High) to ensure complete gelation. Following gelation, matrices were submerged in PBS or cell culture medium depending on the application. ELP-PEG matrices were formed in the same way as HELP matrices, with slight modification: HA-benzaldehyde was replaced with PEG-benzaldehyde and both the PEG and ELP components were dissolved in 1x PBS. Cultrex solutions were stored at -80 °C and thawed on ice prior to use. The appropriate volume of Cultrex solution was pipetted into the corresponding well/mold without any dilution or modification and allowed to incubate at 37 °C for 10 mins prior to submersion in PBS or cell culture medium depending on the application. See Supplementary Table 1 for detailed final formulations for each matrix.

#### Hydrogel rheological characterization

Mechanical characterization was performed on all matrices using a stress-controlled AR-G2 rheometer (TA Instruments) and a cone-plate geometry (20 mm diameter, 1° cone angle, 28 μm gap between the geometry and stage). For HELP matrices, the 2x HA and 2x ELP stock solutions were combined and mixed on the center of the rheometer stage using a pipette tip to form a 48 μL hydrogel. The geometry head was immediately lowered onto the sample, and the crosslinking reaction proceeded under 1% oscillatory strain and 1 rad/s angular frequency for 15 min at 23 °C, followed by a temperature ramp to 37 °C at 2 °C/min, and another time sweep under 1% oscillatory strain and 1 rad/s angular frequency for 10 min at 37 °C. ELP-PEG matrices were formed on the rheometer stage, followed by an initial time sweep at 4 °C for 5 min, a time sweep at 23 °C for 10 min, a ramp to 37 °C at 2 °C/min, and finally a time sweep at 37 °C for 10 min, all under 1% oscillatory strain and 1 rad/s angular frequency. For measuring the mechanics of Cultrex matrices, 48 μL of solution was added to the rheometer stage. The geometry head was immediately lowered onto the sample and the crosslinking reaction proceeded under 1% oscillatory strain and 1 rad/s angular frequency for 5 min at 23 °C followed by a time sweep under the same conditions for 5 min at 37 °C. Finally, for HELP, ELP-PEG, and Cultrex matrices, a frequency sweep from 0.1 to 100 rad/s was performed under 1% oscillatory strain at 37 °C. The final shear modulus for all matrices was derived from the linear region of the frequency sweep at 1 rad/s angular frequency.

#### Fluorescence recovery after photobleaching (FRAP)

The diffusivity within HELP and Cultrex matrices was measured using FRAP analysis based on a previously published procedure^68^. Briefly, HELP and Cultrex matrices (30 μL) were prepared within a well of a clear-bottom, half-area, black 96-well plate (Greiner Bio-One 675090) following the procedures above. Following hydrogel gelation, matrices were submerged in 165 μL of 4 mg/mL solutions of fluorescein isothiocyanate (FITC)-labeled dextrans (Sigma) of varying molecular weights (10k, 20k, 40k, 70k, 150k, 250k Da) in PBS and allowed to incubate for 24 hours at 37 °C in a humidified incubator. Fluorescent images were taken using a confocal microscope (Leica SPE) where a 100 × 100 μm area in each matrix was photobleached using a 488 nm laser at 100% intensity for 1 min. FITC-dextran recovery into the photobleached region was monitored over 4 min (1 frame per second). Images were analyzed using a previously published technique with an accompanying open source MATLAB code “frap_analysis”^69^. Four independent hydrogels were analyzed for each matrix type and for each FITC-dextran molecular weight.

### Human tissue analysis

#### Human tissue collection

Human patient-derived primary PDAC and normal adjacent pancreas tissue were obtained from the Stanford Tissue Bank from patients undergoing surgical resection at Stanford University Medical Center (SUMC). Procedures for generation of human organoid lines from patient tissue samples were approved by the SUMC Institutional Review Board (IRB) and performed under protocol #28908. The organoid line used in this study was derived from a treatment-naïve PDAC tumor. Histological staining and mechanical testing of human tissue were deemed by the IRB to not involve human subjects as defined in U.S. federal regulation 45 CFR 46.102(f) or 21 CFR 50.3(g), as the patient samples were de-identified prior to acquisition. All samples were confirmed to be tumor or normal adjacent by pathological assessment at SUMC. Written informed consent for research was obtained from donors prior to tissue acquisition.

#### Human tissue rheological characterization

Patient-derived normal and PDAC pancreas tissue was collected from surgical-resection and placed in advanced Dulbecco’s Modified Eagle Medium/Ham’s F-12 (ADMEM/F12; Gibco 12634010) supplemented with 10% fetal bovine serum (FBS; Sigma F0804) and 1% Penicillin/Streptomycin/Glutamine (PSQ; Gibco 10378016). Mechanical measurements of tissue were collected between 1-4 hours following surgical resection. Normal and PDAC tissues were cut using a biopsy punch and/or razor blade into 8 mm diameter and 2-4 mm thick sections. Mechanical characterization was performed using a stress-controlled AR-G2 rheometer (TA Instruments) using a parallel plate geometry (8 mm). The rheometer stage and geometry head were affixed with a thin section of sandpaper to prevent tissue slipping during measurements. To perform the measurements, the geometry head was lowered onto the tissue section, and once the normal force reached a value of 0.1-0.2 N, a frequency sweep was immediately performed. The final shear modulus was derived from the linear region of the frequency sweep from the average of 5 data points from 0.6 – 1.5 rad/s angular frequency. All measurements were collected at RT.

#### Immunohistochemistry

Tissue sections were fixed in 4% paraformaldehyde (PFA; Electron Microscopy Sciences 15700) in PBS for 48 hours at 4 °C, washed with PBS (3 × 15 min), paraffin embedded using step-wise dehydration, sectioned (∼5 μm), and affixed to histology slides. Slides were then deparaffinized by two washes in fresh xylene (5 min each), followed by two washes in 100% ethanol (5 min each), one wash in 95% ethanol (1 min), one wash in 70% ethanol (1 min), and were finally submerged in RT Milli-Q water. Heat-induced antigen retrieval was performed via steamer for 30 min while slides were submerged in epitope retrieval solution (IHC World IW-1100). Next, slides were allowed to cool for 20 min before using a hydrophobic pen to isolate the tissue section of interest. Samples were permeabilized with 0.1% Triton X-100 (Thermo Fisher A16046) in PBS (PBST; 2 × 15 min) and blocked with 10% goat serum (Gibco 16210-072) in PBST for 90 min, all at RT. Primary antibodies were diluted in sterile-filtered 0.05% Triton X-100, 0.1% bovine serum albumin (BSA; Sigma A9418), and 0.1% Tween-20 (Thermo Fisher AAJ20605AP) in PBS and incubated with the samples overnight at 4 °C. The following day, the samples were washed with PBST (3 × 15 min) and incubated with corresponding fluorescently-tagged secondary antibodies and 4′,6-diamidino-2-phenylindole (DAPI; 5 mg/mL stock, 1:2000) in the same antibody dilution solution for 2 hours, all at RT. Samples were again washed with PBST (3 × 15 min) and mounted to No. 1 glass cover slips with ProLong Gold Antifade Reagent (Cell Signaling 9071) for 48 hours at RT. Stained samples were imaged using an epifluorescent microscope (Leica Microsystems, THUNDER Imager 3D Cell Culture) and brightness/contrast was adjusted equally across comparative samples using ImageJ (NIH, v.2.1.0/1.53c). See Supplementary Table 3 and 4 for information about primary and secondary antibodies and their dilutions.

### Cell culture and analysis

#### PDAC organoid culture

PDAC organoids consisting of only epithelial cancer cells were derived from surgically-resected patient tissue using previously established methods^70^. For all experiments, PDAC organoids were used between an overall passage of 6 and 15. Where indicated in the main text and figure captions, PDAC organoids were expanded within Cultrex, HELP, or ELP-PEG matrices for at least four consecutive passages prior to performing drug treatment or cell analysis. These passage counts refer to the passage count throughout that specific experiment and not the overall passage count of the PDAC organoids.

PDAC organoid culture within Engelbreth-Holm-Swarm (EHS) matrices, specifically Cultrex Reduced Growth-Factor Basement Membrane Extract, Type 2 (Biotechne R&D Systems 353301002), served as maintenance cultures throughout this study. Organoids were expanded within 40 µL Cultrex hydrogels immobilized in a custom 7 mm diameter silicone mold affixed to a glass cover slip within a 24 well plate as previously reported^33^. Organoids were passaged in Cultrex matrices every 7-10 days upon reaching confluency. To passage organoids, Cultrex hydrogels were first dissociated in 5 mM EDTA on ice for 30-45 min and pelleted at 500 x g for 5 min. Cell pellets were resuspended in 1 mL TrypLE (2 × 3 min incubations at 37 °C, with a pipette mixing in between; Gibco 12604013) to dissociate organoids into a single cell suspension. The solutions were quenched in 40% FBS in PBS and cells were pelleted at 500 x g for 5 min prior to being resuspended in complete WENR medium (see below for full formulation) supplemented with 2.5 µM CHIR99021 (Cayman Chemical 13122) and 10 µM Y27632 hydrochloride (Cayman Chemical 10005583), filtered through a 40 µm cell strainer, and counted. The desired number of cells were then pelleted at 500 x g for 5 min and resuspended in fresh, ice-cold Cultrex solution at 700 cells/µL. Cultrex hydrogels were formed as described above before adding 0.7 mL of fresh, pre-warmed complete WENR medium supplemented with 2.5 µM CHIR99021 and 10 µM Y27632 to each well of a 24-well plate containing one 40 µL hydrogel. After 3 days, medium was changed to complete WENR medium without supplements and medium was changed every 2 days thereafter. Organoids were cultured at 37 °C in a humidified incubator with 5% CO_2_. PDAC organoids were regularly tested for mycoplasma contamination using a MycoAlert Mycoplasma Detection Kit (Lonza LT07-318) and a Lucetta Single Tube Luminometer (Lonza AAL-1001).

To form cell-laden HELP matrices, PDAC organoids were released from Cultrex hydrogels and dissociated into a single cell suspension as described above. The desired number of cells for the given experiment were pelleted at 500 x g for 5 min and resuspended in a 2x stock solution of ELP-hydrazine. HELP matrix formation with cells was performed as described above before adding 0.7 mL of fresh, pre-warmed complete WENR medium supplemented with 2.5 µM CHIR99021 and 10 µM Y27632 to each well of a 24-well plate containing one 40 µL hydrogel. After 3 days, medium was changed to complete WENR medium without supplements and medium was changed every 2-3 days thereafter. If relevant to the given experiment, PDAC organoids were passaged in HELP matrices every 10-14 days upon reaching confluency. To passage organoids, HELP matrices were removed from the silicone molds using a spatula and transferred to a 1.5 mL epi tube containing an equal volume of HELP dissociation solution (PBS supplemented with 200 U/mL elastase (GoldBio E-240-1), 2000 U/mL hyaluronidase (Sigma H3506), 3 mM EDTA, and 20% v/v complete WENR media). This mixture was incubated for 45 min at 37 °C and pipette mixed every 15 min to aid in hydrogel degradation. Samples were then pelleted, dissociated with TrypLE, resuspended in media, and counted as described above for passaging organoids in Cultrex matrices. PDAC organoids were encapsulated in HELP matrices as single cells at 1000 cells/µL or 750 cells/µL for routine passaging or for all comparative experiments, respectively.

The formation of cell-laden ELP-PEG matrices was the same as HELP matrices, with some slight alterations as described above in the “matrix formation” methods section. To passage in ELP-PEG matrices, ELP-PEG hydrogels were collected from culture into a 1.5 mL epi tube containing an equal volume of ELP-PEG dissociation solution (PBS supplemented with 400 U/mL elastase (GoldBio E-240-1), 3 mM EDTA, and 20% v/v complete WENR media) and incubated for 1 hour at 37 °C with regular pipette mixing. The remaining steps to fully release the organoids from ELP-PEG matrices was identical to the HELP protocols above.

#### PDAC organoid media generation

Complete WENR medium used throughout these studies was comprised of a 1:1 mixture of ADMEM/F12 base medium and LWRN cell (ATCC CRL3276) conditioned medium. To prepare the conditioned medium, LWRN cells were expanded for 1-2 days in T150 tissue culture flasks in maintenance medium (DMEM (Gibco 11960044) supplemented with 10% FBS and 1% PSQ). Upon reaching 80% confluence, cells were split 1:4. After 1 day, growth medium was supplemented with 500 µg/ml G418 (Gibco 10131-035) and Hygromycin B (Invitrogen 10687010) selection factors to ensure selection of stable clones containing the expression vectors for Wnt3A, R-spondin 3, and Noggin. Cells were passaged twice more in selection medium and split into several T150 flasks with maintenance medium. Upon reaching 80% confluency, cells were washed with PBS and 25 mL of collection medium (ADMEM/F12 supplemented with 10% FBS and 1% PSQ) was added to the cells. After 24 hours conditioned medium was collected from all flasks, spun down to remove cell debris (2000 x g for 5 min), filtered, and combined into a single container. Fresh collection medium was added to each flask and the conditioned medium was collected in the same manner for up to four collections. To make the complete WENR medium, LWRN conditioned media was combined 1:1 with ADMEM/F12 and supplemented with 1 mM HEPES (Gibco 15630106), 1% GlutaMax (Invitrogen 35050061), 10 mM nicotinamide (Sigma N0636), 1 mM N-acetylcysteine (Sigma A9165), 2% B-27 Supplement without Vitamin A (Invitrogen 12587010), 0.5 µM A83-01 (Cayman Chemical 9001799), 1% PSQ, 10 nM Gastrin I (Cayman Chemical 24457), 10 µM SB-202190 (Cayman Chemical 10010399), 50 ng/mL recombinant human EGF (Peprotech AF-100-15), and 100 µg/mL Normocin (Invivogen ANT-NR-1).

#### Brightfield imaging and organoid diameter quantification

PDAC organoids previously expanded in Cultrex matrices were dissociated into a single cell suspension and encapsulated in 20 μL Cultrex or HELP matrices at a density of 750 cells/μL as above. Organoids were imaged every other day from Day 1 until reaching confluence (Day 15: Cultrex; Day 17: HELP) via phase contrast (Leica Microsystems, THUNDER Imager 3D Cell Culture) using a 10x objective. At least 8 non-overlapping images were taken of N=3 replicate hydrogels for each matrix type resulting in the following number of organoids measured per individual hydrogel per timepoint: Cultrex: n=77-164; HELP Low: n=84-206; HELP Medium: n=93-284; HELP High: n=85-223. Organoid diameter was measured by manually drawing a line over each organoid using ImageJ (NIH, v.2.1.0/1.53c).

#### Gemcitabine drug and ABCG2 inhibitor treatment

To assess PDAC organoid gemcitabine sensitivity through activation of caspase-3, organoids were encapsulated as single cells within the indicated matrices at an initial density of 750 cells/µL. PDAC organoids were grown to an average diameter of ∼75 µm prior to being treated with either 0.1% DMSO (control) or 100 nM gemcitabine hydrochloride (DMSO; Sigma G6423) supplemented in complete WENR media for 3 days. Within the 3 days, one media change with fresh DMSO or gemcitabine was performed 24 hours after the initial drug treatment. Following the drug treatment, organoids were fixed and stained for cleaved caspase-3 expression following the whole mount immunofluorescence protocol below. DAPI and phalloidin were used to stain nuclei and F-actin, respectively. Stained samples for each matrix were imaged using a confocal microscope (Leica SPE) and a 20x objective for a 550 × 550 µm field of view; care was taken to image samples under the same imaging parameters. 5-8 non-overlapping z-stacks (z-range: 100-300 µm; image intervals: 10 µm) were taken per hydrogel per condition. Three biological replicate hydrogels were analyzed for each condition, yielding a range of 20-24 total z-stacks each containing several organoids. Quantification of cleaved caspase-3 signal was performed using ImageJ (NIH, v.2.1.0/1.53c) and all images were blinded to the researcher during analysis. The same analysis pipeline using identical tool/plugin parameters was performed on all samples in the experiment. Briefly, for each image, Li’s minimum cross entropy thresholding method was performed to create a binary mask of positive cleaved caspase-3 signal and the *measure* function was used to find the total cleaved caspase-3 area in µm-squared. For the same image, the *find maxima* and *analyze particles* tools were used to count the number of nuclei. For each z-stack, the total cleaved caspase-3 area was divided by the total nuclei count to calculate the area of cleaved caspase-3 signal per nuclei.

For generation of IC_50_ curves, PDAC organoids were cultured and treated with gemcitabine at one of two stages of organoid culture: (1) drug treatment throughout the log-phase expansion of single cells into multicellular organoids or (2) drug treatment upon formation of ∼75-μm diameter multicellular organoids. For these experiments, Cultrex, HELP, and ELP-PEG matrices (25 μL) were prepared within a clear-bottom, half-area, black 96-well plate (Greiner Bio-One 675090) following the procedures above. For each experiment, a total of 3-5 replicate hydrogels were cast for each drug concentration within a given condition.

For organoid drug treatment, PDAC organoids were seeded as single cells in their respective matrices at a density of 750 cells/μL and were grown to an average diameter of ∼75 µm prior to being treated with DMSO (0.1%), 3, 33, 66, 100, 200, 333, 1000, or 3333 nM gemcitabine for 3 days. Within the 3 days, one media change with fresh DMSO or gemcitabine was performed 24 hours after the initial drug treatment. Following drug treatment, cell toxicity was measured for each sample using a CytoTox Glo Assay (Promega G9290) and a LUMIstar Omega microplate luminometer (BMG Labtech). Toxicity measurements were normalized to DMSO (i.e. 0) and 3333 nM gemcitabine (i.e. 1). A least squares nonlinear regression method (“[Inhibitor] vs. response -Variable slope (four parameters)”) was used to calculate the best fit and IC_50_ value for each experimental group using GraphPad Prism software (v.9.3.1).

For single cell drug treatment, PDAC organoids were seeded as single cells in their respective matrices at a density of 750 cells/μL and were grown for 2 (Cultrex) or 4 (HELP and ELP-PEG) days prior to being treated with DMSO (0.1%), 3, 10, 33, 66, 100, 200, 333, or 3333 nM gemcitabine for 6 days. Media supplemented with DMSO or gemcitabine was replaced every other day. Following drug treatment, cell viability was measured for each sample using a Cell Titer Glo 3D Assay (Promega G9681) and a LUMIstar Omega microplate luminometer (BMG Labtech). Viability measurements were normalized to DMSO (i.e. 1). The best fit and IC_50_ values were calculated as described above.

To test the role of ABCG2 inhibition on PDAC organoid gemcitabine sensitivity, PDAC organoids were encapsulated in Cultrex or HELP matrices and treated with drug/inhibitor following the *single cell* and *organoid* treatment protocols described above. Specifically, for *single cell* treatment, cells were treated with either 0.2% DMSO, 33/66 nM gemcitabine, or 33/66 nM gemcitabine + 20 μM Ko143 (Cayman Chemical 15215). For *organoid* treatment, organoids were treated with either 0.2% DMSO, 100 nM gemcitabine, 100 nM gemcitabine + 20 μM Ko143, or 3333 nM gemcitabine (positive control). PDAC viability upon treatment of 20 μM Ko143 alone was measured following the *single cell* treatment protocol.

#### Quantitative polymerase chain reaction (qPCR)

For qPCR analysis, PDAC organoids cultured within 40 μL Cultrex, HELP, or ELP-PEG matrices were dissociated for 20-30 min at 37 °C in 40 μL of HELP or ELP-PEG dissociation solution. Cultrex matrices were dissociated in HELP dissociation solution to expose all cells to the same treatment. Next, the samples were immediately resuspended in 500 μL of TRIzol reagent (Invitrogen 15596018) and frozen at -80 °C until use. mRNA was purified from lysates using a phenol-chloroform extraction. First, samples were disrupted via probe sonication (Heilscher UP50H, 50% amplitude (25 watts), 30kHz frequency, 0.5 cycle), transferred to a phase lock gel (Quantabio 5PRIME 2302830), and supplemented with 100 μL of chloroform (Sigma CX1055). Samples were then centrifuged at 15,300 x g at 4° C for 15 min and the top, aqueous phase was transferred to a clean 1.5 mL epi-tube. Samples were precipitated with 1 wash of isopropyl alcohol followed by two washes of 70% ethanol with centrifugation steps between each wash (18,500 x g at 4 °C for 10 min). After decanting the final ethanol wash, samples were dried and resuspended in 15-30 μL of nuclease free water. mRNA concentrations were measured via NanoDrop (Thermo Scientific) and a consistent amount of mRNA across all samples (100-1000 ng) was reverse transcribed using a High-Capacity cDNA Reverse Transcription Kit (Applied Biosystems 4368814). qPCR was performed on 6.6 μL of diluted cDNA per gene target mixed with 0.9 μL of 5 μM forward and reverse primer pair solution and 7.5 μL of Fast SYBR Green Master Mix (Applied Biosystems 4385612). Samples were run on a StepOnePlus Real Time PCR System (Applied Biosystems). CT values were calculated using the StepOnePlus software (v.2.3) and analyzed by the ΔCT method. Statistical analysis was performed prior to transforming to a natural scale as ΔCT values approximate a normal distribution. mRNA expression data throughout the paper is reported as a geometric mean with asymmetric 95% confidence intervals derived from the non-transformed data. See Supplementary Table 5 for information about qPCR primers.

In Figure 4j, modeling the *single-cell* gemcitabine drug treatment protocol, PDAC cells were cultured for two days in Cultrex or 4 days in HELP matrices prior to the addition of either 0.1% DMSO or 20 μM Ko143 to complete WENR medium. Upon the addition of Ko143, cell culture medium was changed every 24 hours for the next 6 days, and Ko143 was diluted fresh in the medium prior to addition to the organoids. Cells were collected for qPCR analysis on Day 8 (Cultrex) or Day 10 (HELP).

#### Western blot

PDAC organoids previously cultured in Cultrex, HELP Low, or HELP High matrices were encapsulated within 40 μL Cultrex or HELP as described above at an initial concentration of 750 cells/μL. Modeling the *single cell* gemcitabine drug treatment protocol, PDAC cells were cultured for 2 days in Cultrex or 4 days in HELP matrices prior to the addition of either 0.1% DMSO or 20 μM Ko143 to complete WENR medium. Upon the addition of Ko143, cell culture medium was changed every 24 hours for the next 6 days, and Ko143 was diluted fresh in the medium prior to addition to the organoids. After 6 days, to release the organoids from the matrices, each hydrogel was dissociated for 45 min at 37 °C in 40 μL of HELP dissociation solution. Cultrex matrices were also dissociated using the same solution to expose all cells to the same treatment. Samples were pipette mixed throughout to aid in matrix dissociation. Next, samples were resuspended in 120 μL of RIPA lysis buffer solution supplemented with 1 mM PMSF and protease inhibitor tablets (Roche 11873580001), disrupted via sonication (Heilscher UP50H, 50% amplitude (25 watts), 30kHz frequency, 0.5 cycle), centrifuged at 18,000 x g for 10 min at 4 °C to clear any debris, and frozen at -80 °C until use. Samples were then thawed on ice and 20 μL of lysate for each replicate was combined with 5 μL of 5x Laemmli buffer (50% v/v glycerol, 10 wt% sodium dodecyl sulfate, 0.05 wt% bromophenol blue, 300 mM Tris pH 6.8) supplemented with fresh 0.5 M dithiothreitol prior to incubating at 96 °C for 10 min to denature the protein. Pre-cast polyacrylamide gels (Bio-Rad 4561043) were loaded with 25 μL sample per well and 10 μL of protein ladder (Bio-Rad 1610375). The gel was run at 140 V for 90-120 min depending on the protein of interest. The separated proteins were transferred to a methanol activated polyvinylidene fluoride (PVDF) membrane (Invitrogen LC2005) via wet transfer for 1 hour at 100 V. At this point, the membranes were cut to enable staining of different molecular weight proteins from the same run. Next, the membrane was blocked for 1 hour in 5 wt% milk in tris buffered saline (20x stock: 3 M NaCl, 750 mM Tris hydrochloride, pH 7.2) supplemented with 0.2% v/v Triton X-100 (TBST). Primary antibodies were diluted in TBST and incubated overnight while rocking at 4 °C. The following day, membranes were washed in TBST (3 × 10 min) prior to addition of the horseradish peroxidase-conjugated secondary antibody diluted in TBST for 1 hour at RT. The membrane was washed in TBST (4 × 10 min) and developed using either the SuperSignal West Pico or Femto Chemiluminescent Substrate (Thermo Fisher 34580, 34094), and imaged using a ChemiDoc MP gel imaging system (Bio-Rad). Densitometry analysis was performed on a preliminary blot for each sample stained for β-Actin (loading control) using Image Lab software (Bio-Rad, v.6.0.0), and the results were used to normalize the amount of sample loaded for all Western blot data included in the manuscript. The same densitometry tools were used to quantify the blots in Supplementary Figure 20. All individual replicates (N=4) per matrix type were kept separate throughout matrix dissociation and Western blot analysis. See Supplementary Table 3 and 4 for information about primary and secondary antibodies and their dilutions.

#### Immunofluorescence imaging

PDAC organoids cultured within HELP and Cultrex matrices were prepared for immunostaining using two different methods: (1) paraffin embedding and sectioning or (2) whole mount *in situ* 3D staining and imaging.

For paraffin embedded samples, HELP and Cultrex matrices containing PDAC organoids were washed with 1 mL PBS and fixed with 1 mL 4% PFA and 0.1% glutaraldehyde (Thermo Fisher BP25481) in PBS for 48 hours at 4 °C. Samples were then washed with 200 mM glycine in PBS (1 × 15 min) to quench any remaining glutaraldehyde followed by washes with PBS (2 × 10 min). Subsequent paraffin embedding, sectioning, staining, and mounting were performed as described above for human tissue sections. Stained samples were imaged using an epifluorescent microscope (Leica Microsystems, THUNDER Imager 3D Cell Culture) and care was taken to scale intensity values equally across all images when comparing protein expression across samples using ImageJ (NIH, v.2.1.0/1.53c). See Supplementary Table 3 and 4 for information about primary and secondary antibodies and their dilutions.

For whole mount 3D samples, HELP and Cultrex matrices containing PDAC organoids were washed with 1 mL PBS and fixed with 1 mL 4% PFA and 0.1% glutaraldehyde in PBS for 30 min at 37 °C. Samples were then washed with 200 mM glycine in PBS (1 × 15 min) to quench any remaining glutaraldehyde followed by washes with PBS (2 × 10 min). Next, the samples were permeabilized with 1 mL PBST for 1 hour at RT while rocking and subsequently blocked with 1 mL 10% goat serum in PBST for 4 hours at RT. Primary antibodies were diluted in sterile-filtered 0.05% Triton X-100, 0.1% BSA, and 0.1% Tween-20 in PBS and 400 uL of primary antibody solution was added to each sample to incubate overnight at 4 °C. The following day, the samples were washed with 1 mL PBST (3 × 15 min) at RT and incubated with 400 uL of the corresponding fluorescently-tagged secondary antibodies, DAPI (5 mg/mL stock, 1:2000), and/or tetramethylrhodamine (TRITC)-phalloidin (100 ug/mL in DMSO stock, 1:500) in the same antibody dilution solution overnight at 4 °C. Samples were again washed with PBST (3 × 15 min) and mounted to No. 1 glass cover slips with ProLong Gold Antifade Reagent for 48 hours at RT. Stained samples were imaged using a confocal microscope (Leica SPE) and care was taken to scale intensity values equally across all images using ImageJ (NIH, v.2.1.0/1.53c) when comparing protein expression across samples. See Supplementary Table 3 and 4 for information about primary and secondary antibodies and their dilutions.

#### Organoid barcoding and analysis

Genetically-barcoded PDAC organoids were generated as previously described^39^. For this work, genetically-barcoded PDAC organoids were expanded within Cultrex, HELP Low, or HELP High matrices for a total of three passages from the initial parent population established within Cultrex. For each passage, organoids were expanded in their respective matrices until reaching confluence and organoids were released from the matrices and dissociated into a single cell solution as described above (i.e. Cultrex and HELP matrices were dissociated using the same dissociation solutions and protocols). 1 million cells were collected for DNA barcode sequencing from the initial parent population and between each passage from N=3 independent biological replicates from each matrix type. These samples were washed with PBS, pelleted, and frozen at -80 °C until use. DNA extraction was performed using either a QIAamp DNA Micro Kit or DNeasy Blood and Tissue Kit (Qiagen 56304/69504) and dsDNA was measured using Qubit Fluorometer (Invitrogen Q33238) and Qubit dsDNA HS Assay Kit (Invitrogen Q32851).

Barcode sequencing and analysis was performed similarly to previous work^39^. Briefly, the barcode region was amplified by PCR using 2X KAPA Hifi PCR Master Mix (Roche Sequencing Solutions KK2601) with a minimum of 200 ng of DNA input per sample. The PCR product was purified using Ampure magnetic beads (Beckman Coulter A63881), and then subjected to a second PCR reaction where sequencing adaptors with sample specific indexes were added. The PCR products of all samples were then combined and purified using a Qiagen PCR purification kit. To remove primer dimers, the purified product was run on a 2% EX gel (Thermo Fischer G401002) and the 200 bp band was isolated and gel purified using a Qiaquick gel extraction kit (Qiagen 28706). The purified product was bioanalyzed and confirmed for a 204 bp peak prior to sequencing with 150 bp paired-end reads on an Illumina MiSeq.

For extraction of cell barcodes from sequencing fastq files, only reads exactly matching the sequence around the 30 base-pair barcode were retained to ensure read quality. Then, reads from all samples were combined into a single file, and UMI-tools^71^ was used to merge similar barcode reads (i.e. reads where the difference between barcodes was more likely to depend on sequencing and PCR errors, as compared to being a separate barcode). Each such group of combined barcodes was called a read group. Barcode reads for individual samples were then matched to the file containing all read groups, and associated barcode reads and counts for each read group were summed up per sample. For each sample, the number of reads per barcode were normalized by the total read count to calculate the barcode frequency. To mitigate noise, a frequency cutoff of 0.01% was applied to all samples. For analysis of barcodes present at passage 3, a barcode was only included if it met the frequency cutoff in all three biological replicates for a given matrix. Data analysis was conducted using MATLAB (v.9.11.0.1769968). In Figure 4l, some barcodes may have average frequencies <0.01% because the specific clone had an average frequency >0.01% in at least one matrix type. In Supplementary Figure 9 and 10, barcode frequencies at passage three within each matrix were normalized by its initial frequency in the Parent population.

#### Side population assay and cell cycle analysis

PDAC organoids expanded within Cultrex or HELP matrices were collected and dissociated into single cells as above. Single cells were immediately resuspended in warm complete WENR media supplemented with 5% FBS and 10 μM Y27632 (Cayman Chemical 10005583) solubilized in water at a density of 10^6^ cells/mL. Hoechst 33342 (Thermo Fisher H3570) was added to all conditions at a concentration of 5 µg/mL, and Ko143 (200 nM; Cayman Chemical 15215) or Verapamil (50 µM; Sigma V4629) solubilized in water were added to inhibition samples. Ko143 was first solubilized in DMSO at 10 μM and then diluted in PBS to 4 μM before being added to the media at a final concentration of 200nM. Samples were immediately moved to 37 °C for 2 hours, with gentle mixing every 15 min to reduce cell settling. After treatment, cell pellets were collected (5 min, 500 x g) and resuspended in ice-cold flow buffer (PBS + 5% FBS) with 7-AAD viability dye (5:100; Thermo Fisher, 00-6993-50) and immediately used for flow cytometry on a BD FACS Symphony A5. The flow rate was kept to below 200 events per second and a minimum of 50,000 live, single cell events were recorded for each sample. Side population was detected using a 428/31 band-pass filter (Hoechst-Blue) and a 670/30 band-pass and 635 long-pass filter (Hoechst-Red); excitation with a UV (305 nm) laser. Verapamil samples were used to confirm gating strategy for identification of the side population. DNA content for cell cycle analysis was detected using a 428/31 band-pass filter and excited with a UV (305 nm) laser. Cells in s-phase were identified as 2n<DNA<4n. All flow cytometry analysis was performed with FlowJo software (v.10.7.1).

#### Flow cytometry on cancer stem cell markers

PDAC organoids expanded within Cultrex or HELP matrices were collected and dissociated into single cells as above. Single cells were immediately resuspended in ice-cold flow buffer (PBS + 5% FBS) with Fc receptor block (20:100; Thermo Fisher 14-9161-73). After 20 min, samples were stained for surface protein expression for 30 min on ice. Samples were washed in ice-cold flow buffer and pelleted (5 min, 500 x g). Samples were resuspended in flow buffer with 7-AAD viability dye (5:100; Thermo Fisher 00-6993-50) and immediately used for flow cytometry on a BD FACS Symphony A5. A minimum of 50,000 live, single cell events were recorded for each sample. Compensation was set using UltraComp eBeads Compensation Beads (Thermo Fisher 01-2222-41) and unstained cells. Fluorescence minus one (FMO) controls were used to set gating strategies. All antibody stains were titrated prior to use to identify optimal concentration for separation of negative and positive populations. See Supplementary Table 3 for antibodies and dilutions. All flow cytometry analysis was performed with FlowJo software (v.10.7.1). All values are reported as the median intensity of the live population.

### Other

#### Kaplan Meier survival curve analysis

Analysis of PDAC patient survival was performed using a previously published online tool^45^ collecting data from The Cancer Genome Atlas (TCGA) dataset. In this analysis, patients were stratified by the upper and lower quartile expression of the given marker and overall survival was plotted. “Pancreatic ductal adenocarcinoma (n=177)” was selected as the cancer type. The analysis was not restricted by subtype or cellular content. Follow up threshold was set to “all” and “Censore at threshold” was selected. The raw data output was plotted and analyzed using GraphPad Prism v.9.3.1 software. The log-rank (Mantel-Cox) test was used for statistical analysis.

#### Tumor vs normal RNA-sequencing analysis

Analysis of RNA-sequencing (RNA-seq) data was performed using a previously published online tool^32^ collecting data from TCGA and The Genotype Tissue Expression Project (GTEx). The pancreatic adenocarcinoma (“PAAD”) dataset was used. The data was plotted on a log scale (log_2_(TPM + 1)) with a jitter size of 0.4. “Match TCGA normal and GTEx data” was selected.

#### Statistical analysis

Statistical analyses for this study were performed using GraphPad Prism v.9.3.1 software. Details of specific statistical methods and p-value results are included within the figure captions. For all studies, ns = not significant (p>0.05), * = p<0.05, ** = p<0.01, *** = p<0.001, **** = p<0.0001.

## Acknowledgments

We thank Thomas Lozanoski, Erinn Rankin, Suchitra Natarajan, Julien Roth, and Christopher Madl for insightful conversations and editing of the manuscript. We are grateful to the patients who donated tissue samples used for this research. We thank the Stanford Tissue bank for their assistance in procuring patient tissue samples. We thank Zhicheng Ma for help with barcode DNA sequencing. We thank Pauline Chu at the Stanford Human Pathology/Histology Service Center for assistance with tissue and organoid paraffin embedding and sectioning. B.L.L. acknowledges financial support from the Siebel Scholars Program and Stanford Bio-X Bowes Graduate Fellowship. B.A.K. acknowledges financial support from the Stanford Bio-X Bowes Graduate Fellowship. K.K. acknowledges financial support from the Swedish Research Council (2018-00454). K.C.K. acknowledges financial support from the National Science Foundation Graduate Research Fellowship Program. This work was supported by funding from the National Institutes of Health (R01 EB027171 to S.C.H.; R01CA2515143, U01CA217851, and U54CA224081 to C.J.K.; U01CA217851 and DP1CA238296 to C.C.), Stand Up to Cancer (C.J.K.), and the National Science Foundation (CBET 2033302 to S.C.H.). Flow cytometry data was collected on BD FACS Symphony A5 in the Shared FACS Facility obtained using NIH S10 Shared Instrument Grant (1S10OD026831-01). Part of this work was performed at the Stanford Nano Shared Facilities (SNSF), supported by the National Science Foundation under award ECCS-2026822.

## Competing Interests

S.C.H. has a patent application pending related to the engineered matrix. C.C. is an advisor and holds equity in Grail, Ravel, and DeepCell, and is an advisor to Genentech and NanoString. The rest of the authors have no competing interests.

## Data Availability

All data needed to evaluate the conclusions in the paper are present in the manuscript and/or the Supplementary Information. Raw data related to this paper are available for research purposes upon reasonable request.

## Author contributions

B.L.L., A.E.G., B.A.K., A.R.S., S.C.H. designed the research; B.L.L., A.E.G., B.A.K., K. Karagyozova, K. Klett conducted experiments; B.L.L., A.E.G., K. Karlsson analyzed data; K. Karlsson, A.R.S., C.C., C.J.K. provided organoids and reagents; B.L.L. wrote the manuscript; B.L.L., A.E.G., B.A.K., K. Karlsson, A.R.S., K. Karagyozova, K. Klett, C.C., C.J.K., S.C.H. edited and approved the final manuscript.

