## Supplementary Information for "Engineered extracellular matrices reveal stiffness-mediated chemoresistance in patient-derived pancreatic cancer organoids"

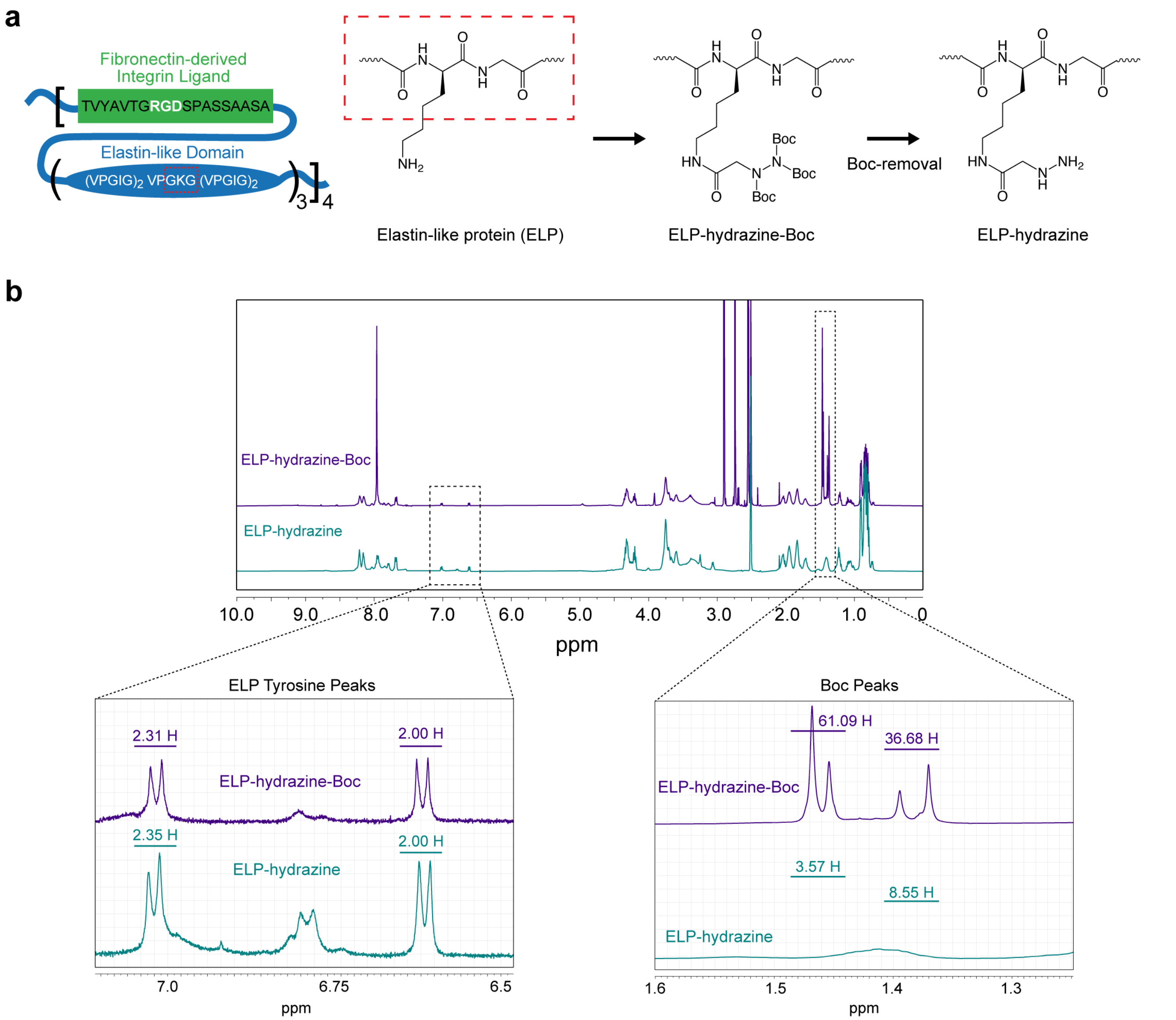


**c**

$$\frac{Hydrazine\ groups}{ELP} = \frac{Boc\ peak\ H}{Tyrosine\ peak\ H} * \frac{4\ H}{tyrosine} * \frac{4\ tyrosine}{ELP} * \frac{1\ Boc}{9\ H} * \frac{1\ hydrazine}{3\ Boc}$$

$$\frac{Hydrazine\ groups}{ELP} = \frac{97.77\ H}{4.31\ H} * \frac{4\ H}{tyrosine} * \frac{4\ tyrosine}{ELP} * \frac{1\ Boc}{9\ H} * \frac{1\ hydrazine}{3\ Boc} = 13.44$$

$$\frac{13.44}{14\ possible\ hydrazine\ groups} = \sim 96\%\ modification\ of\ ELP$$

**Supplementary Figure 1. Elastin-like protein (ELP) modification with a hydrazine functional group.** **a**, Schematic of ELP amino acid sequence and step-wise chemical modification with a hydrazine group. The hydrazine is conjugated to the lysine amino acid within the elastin-like domain of ELP. **b**, Representative proton nuclear magnetic resonance ($^1$H NMR; $D_2O$ solvent) spectra of final ELP-hydrazine polymer (green) and intermediate ELP-hydrazine-Boc (purple). The tert-butyloxycarbonyl (Boc) protecting group is completely removed from the hydrazine ($\delta$ = 1.46 and 1.39 ppm) in the final ELP-hydrazine polymer. Tyrosine peaks ($\delta$ = 6.62, d, 2H and $\delta$ = 7.00, d, 2H) from the ELP serve as a baseline for normalizing $^1$H NMR signal. **c**, Calculation of ELP hydrazine modification.

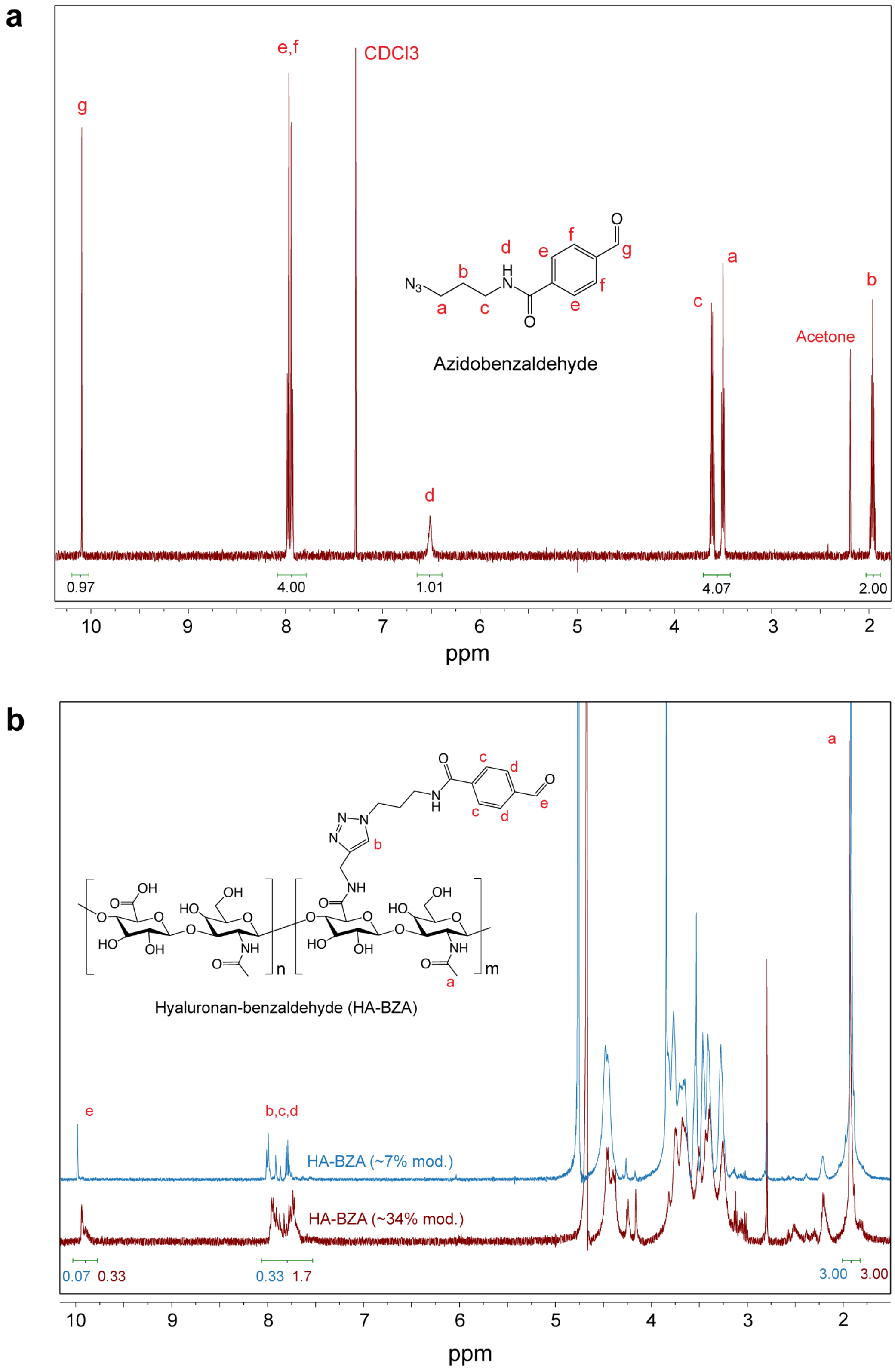


**Supplementary Figure 2. Hyaluronan (HA) modification with a benzaldehyde functional group. a**, Representative $^{1}$H NMR spectrum (CDCl3 solvent) of small molecule precursor azidobenzaldehyde. **b**, Representative $^{1}$H NMR spectra ($D_2O$ solvent) of either ~7% or ~34% modified HA-benzaldehyde polymer.

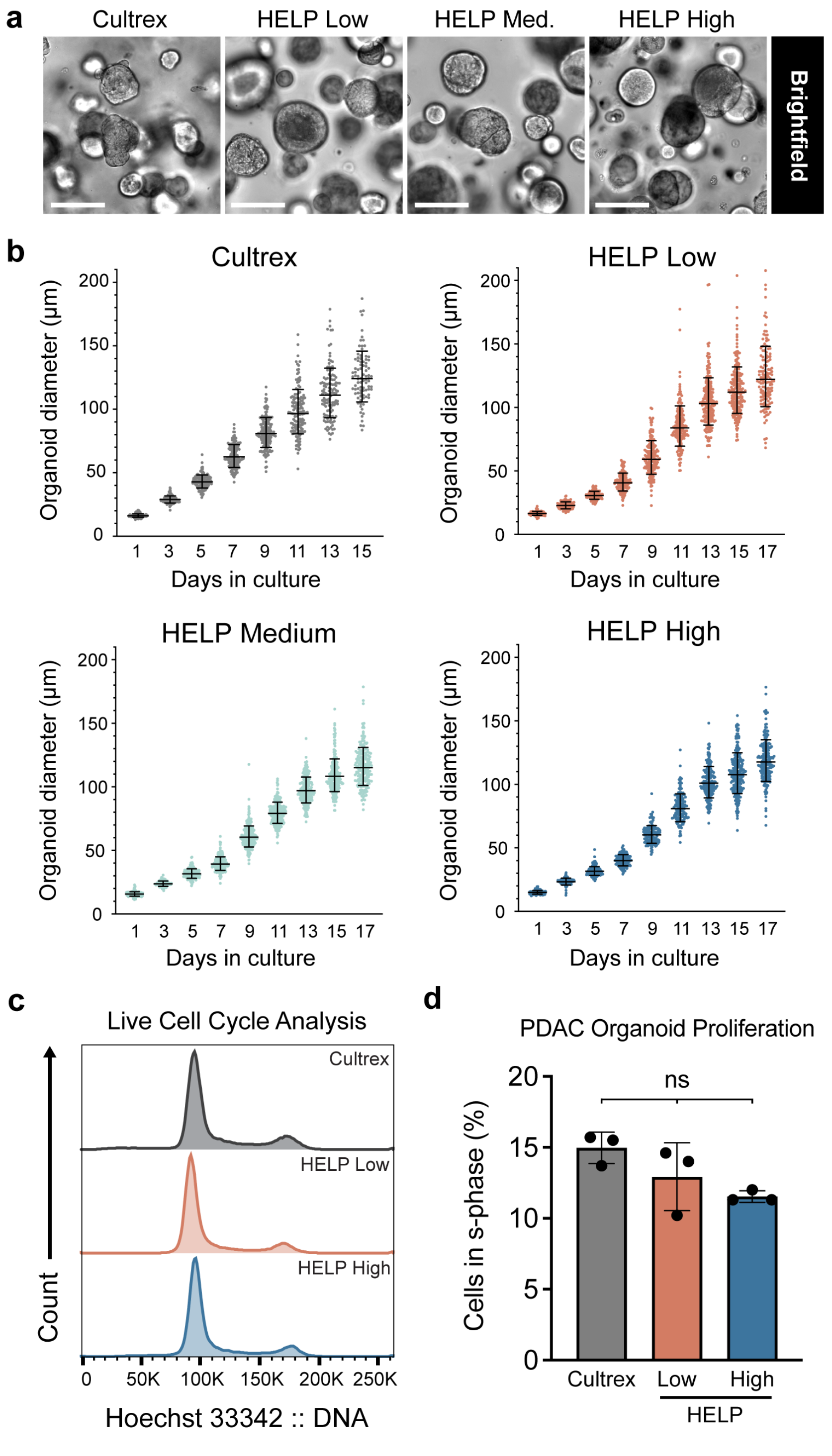


**Supplementary Figure 3. PDAC organoids show similar proliferation within Cultrex and HELP matrices of varying stiffness. a**, Representative brightfield images of PDAC organoids expanded for one passage within Cultrex and HELP matrices. Scale bar, 100 µm. **b**, Quantification of PDAC organoid diameter during culture within Cultrex and HELP matrices for up to 17 days. Each data point represents a diameter measurement from a single organoid (Cultrex: n=77-164; HELP Low: n=84-206; HELP Medium: n=93-284; HELP High: n=85-223). Data is compiled from N=3 matrices for each timepoint. **c**, Representative histogram of DNA content of PDAC organoids expanded for four passages within Cultrex and HELP matrices. **d**, Proportion of PDAC cells in s-phase following four passages within Cultrex (d8) and HELP (d10) matrices (N=3, mean ± SD, ordinary one-way ANOVA with Tukey multiple comparisons correction, ns = not significant).

Gating Strategy - Live Single Cells

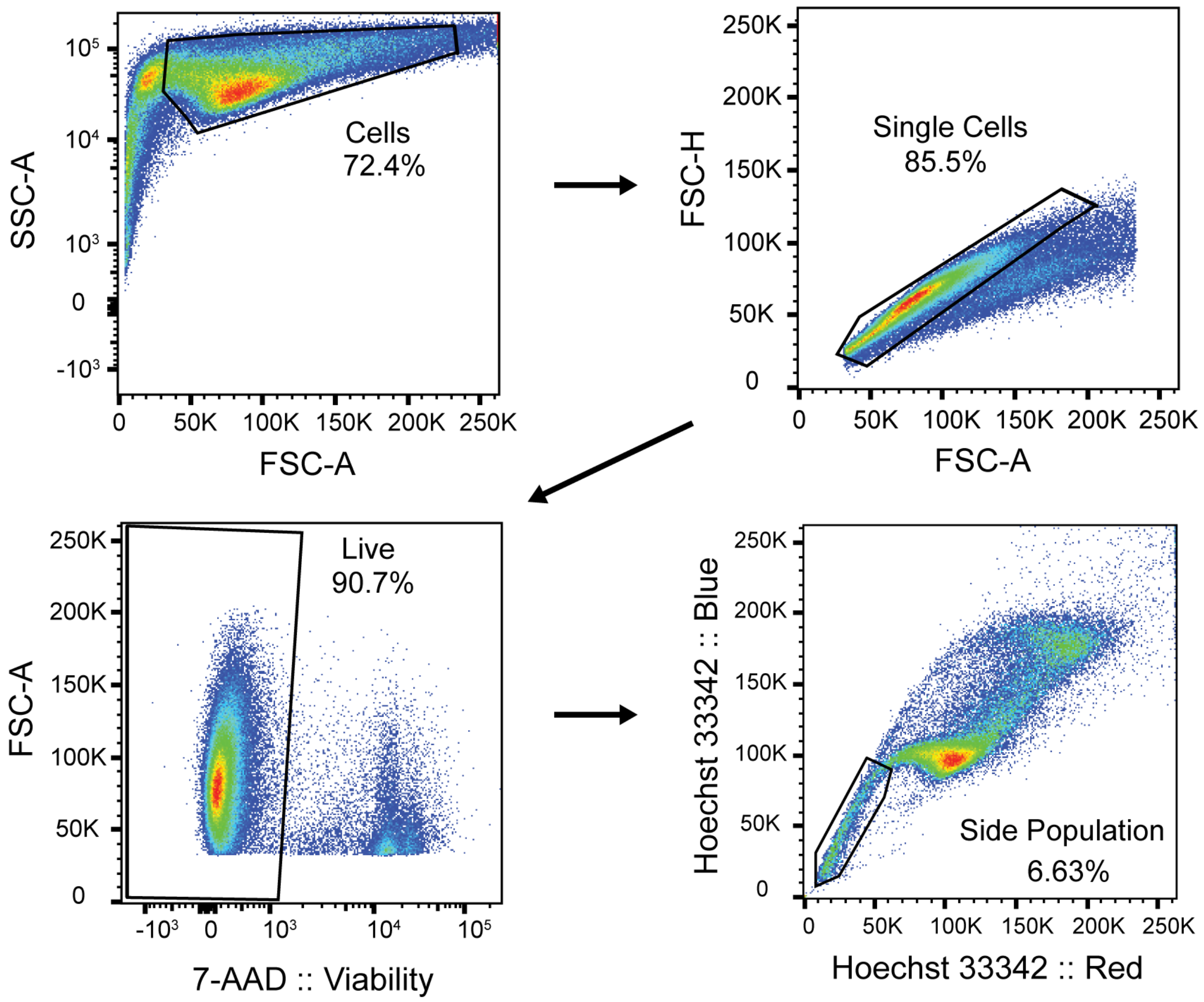


**Supplementary Figure 4. Gating strategy for flow cytometry analyses.** Gating strategy for live single cells used for all flow cytometry experiments. For side population experiments, samples treated with verapamil to inhibit drug efflux transporters were used to confirm gating strategy for identification of the side population.

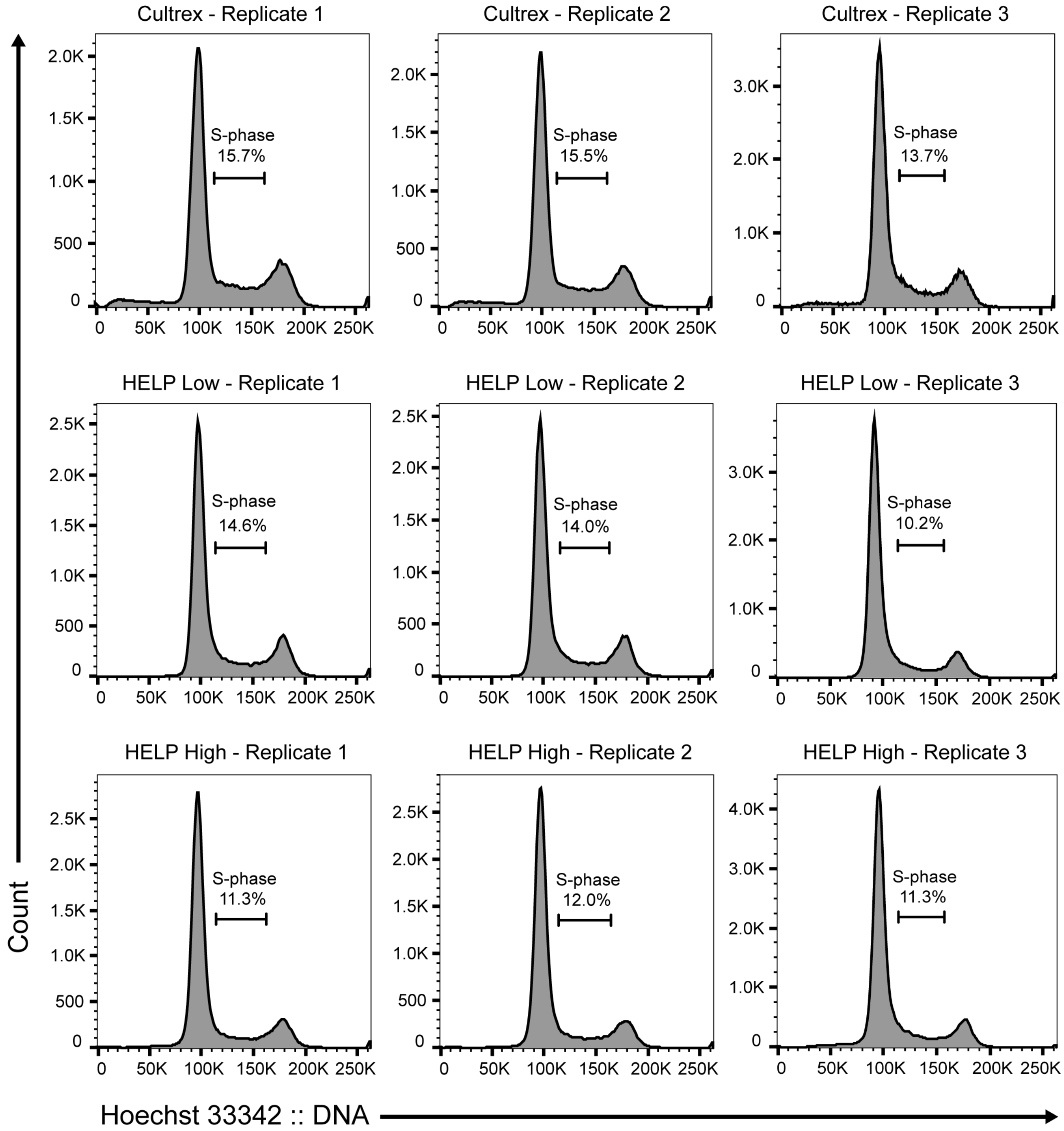


**Supplementary Figure 5. PDAC organoids show similar proportion of cells in s-phase when expanded within Cultrex and HELP matrices.** Replicate histograms of DNA content of PDAC cells expanded for four passages within Cultrex (d8, top), HELP Low (d10, middle) or HELP High (d10, bottom) matrices. Percentage of actively proliferating cells in s-phase is denoted for each sample. All data is gated for live, single cells.

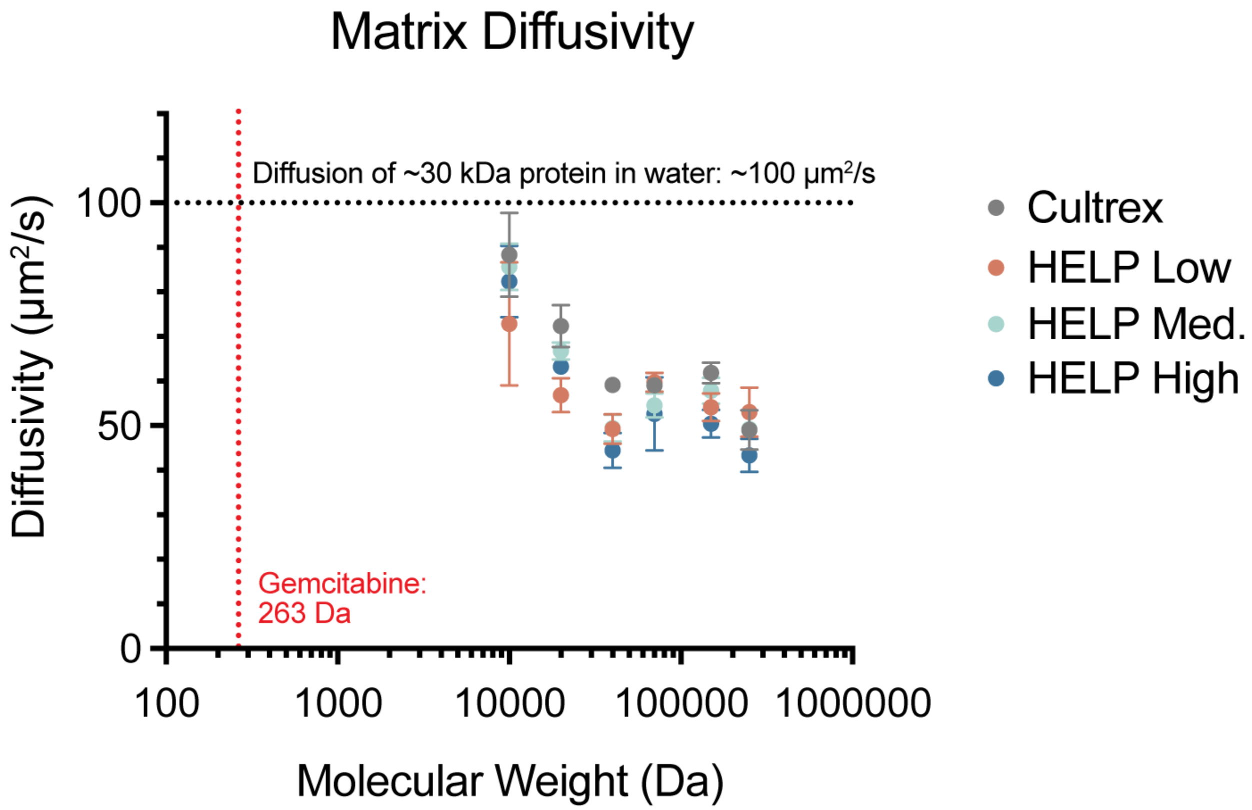


**Supplementary Figure 6. Diffusive transport properties are similar across Cultrex and HELP hydrogels of varying stiffness.** Diffusivity of soluble FITC-Dextran macromolecules of varying molecular weight within Cultrex and HELP matrices. (N=3-4, mean ± 95% confidence interval). Molecular weight of gemcitabine drug is denoted with a red dashed line. The approximate diffusivity of a ~30 kDa protein in water was calculated using the Stokes-Einstein equation[1] and is denoted with a black dashed line.

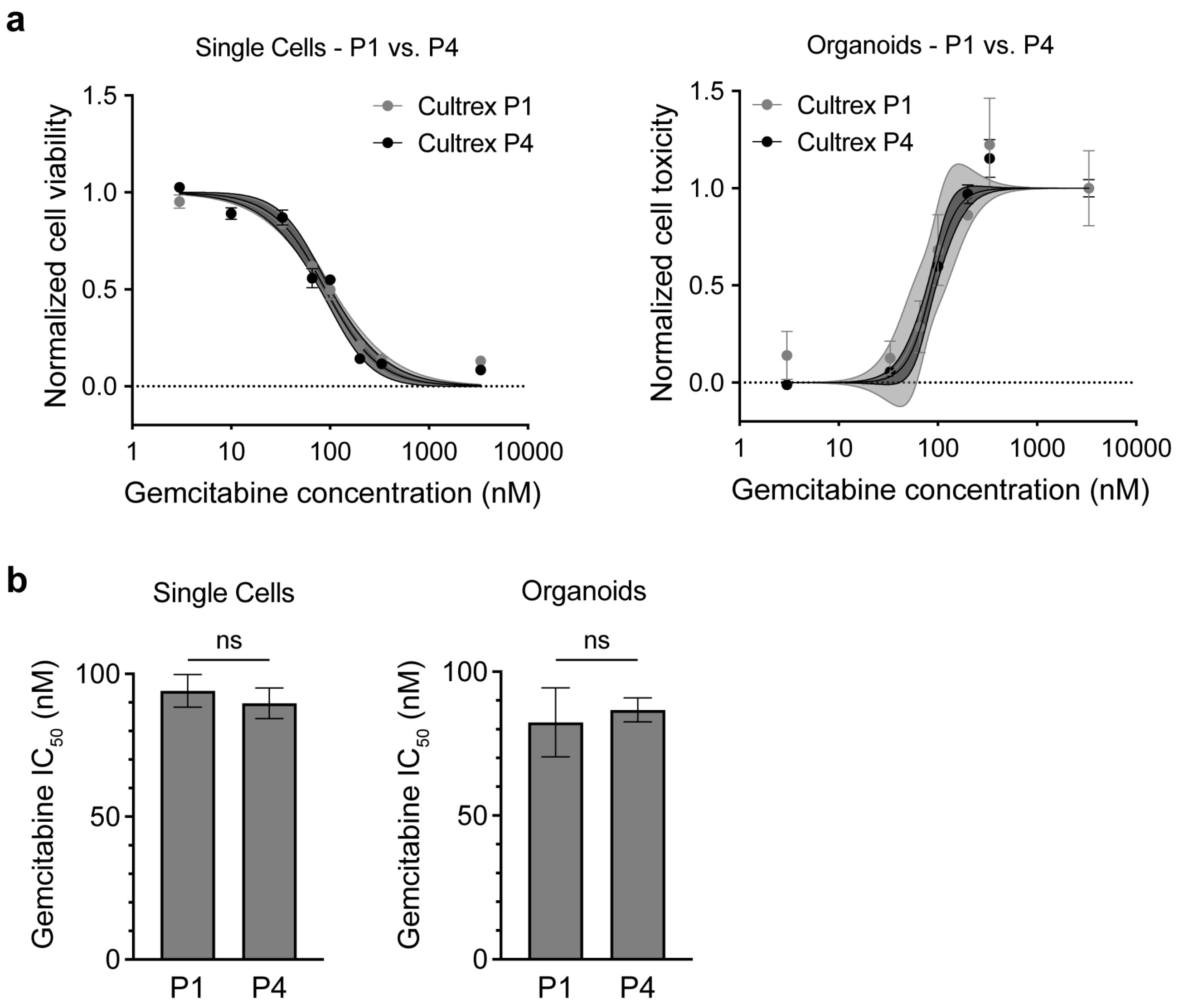


**Supplementary Figure 7. PDAC organoids expanded within Cultrex maintain stable drug sensitivity across passages. a**, Gemcitabine dose-response curves for PDAC organoids expanded within Cultrex matrices for either one (P1) or four (P4) passages. PDAC cells were treated with gemcitabine either throughout log-phase single cell expansion (left) or following organoid formation (right) (N=4, data points are mean ± SEM, solid center line is nonlinear least squares regression of data; shaded region represents 95% confidence bands of nonlinear fit; data are normalized to positive controls (DMSO for single cells; 3333 nM gemcitabine for organoids). **b**, Gemcitabine $IC_{50}$ values calculated from nonlinear fit of dose-response curves from **a** (N=4, mean ± SEM, unpaired two-tailed Student's t-test, ns = not significant).

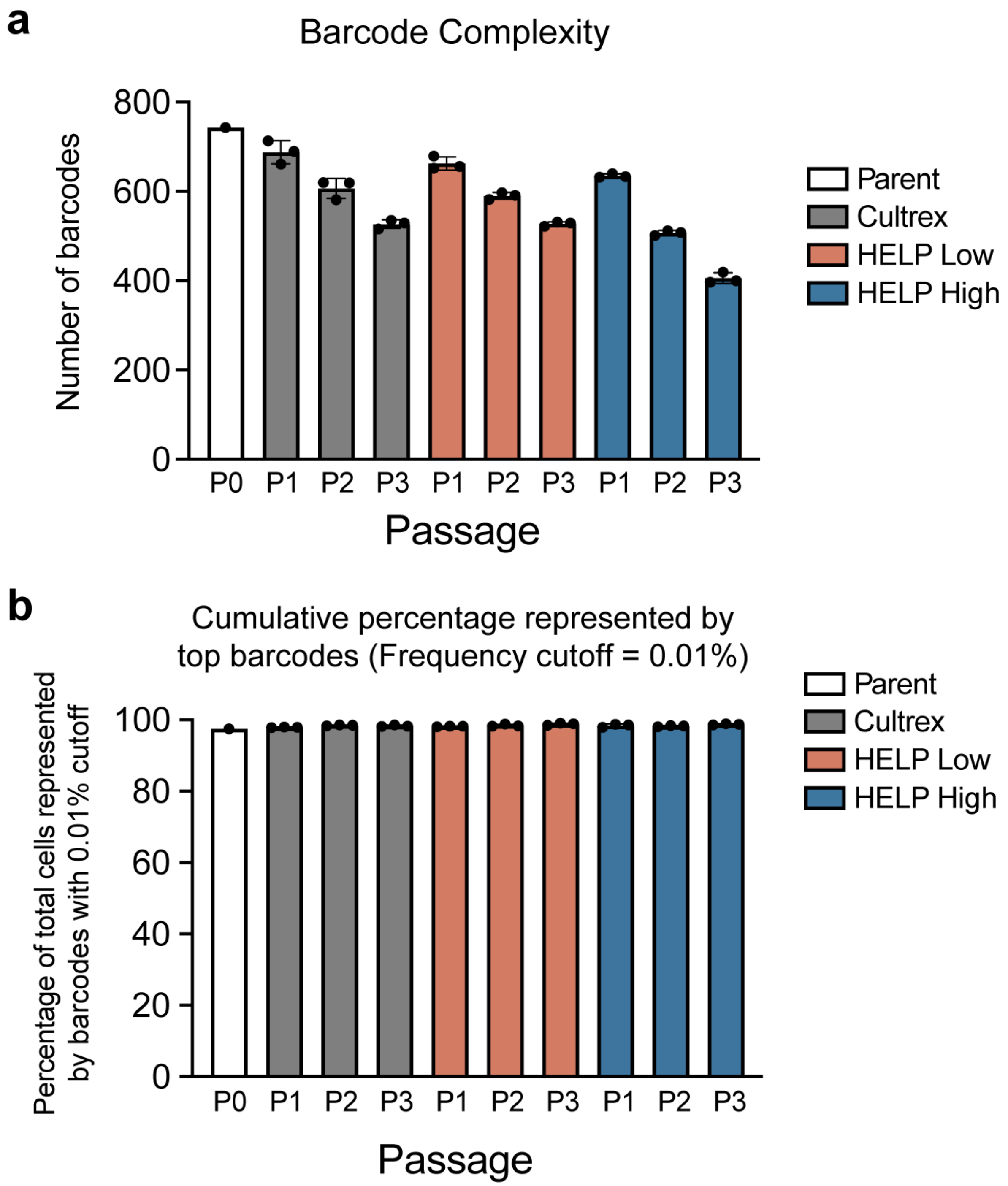


**Supplementary Figure 8. High stiffness HELP matrices impart steeper clonal selection pressure on PDAC organoids. a**, Number of unique barcodes present within PDAC organoid populations expanded within Cultrex, HELP Low, or HELP High matrices for one to three passages (N=3, mean ± SD). The Parent population was the same for all matrices and is indicated by the white bar (N=1). Only barcodes with a population frequency >0.01% were included. Biological replicates were kept separate throughout all passages. **b**, Cumulative frequencies of the unique barcodes (cutoff: >0.01%) for each matrix at each timepoint (N=3, mean ± SD).

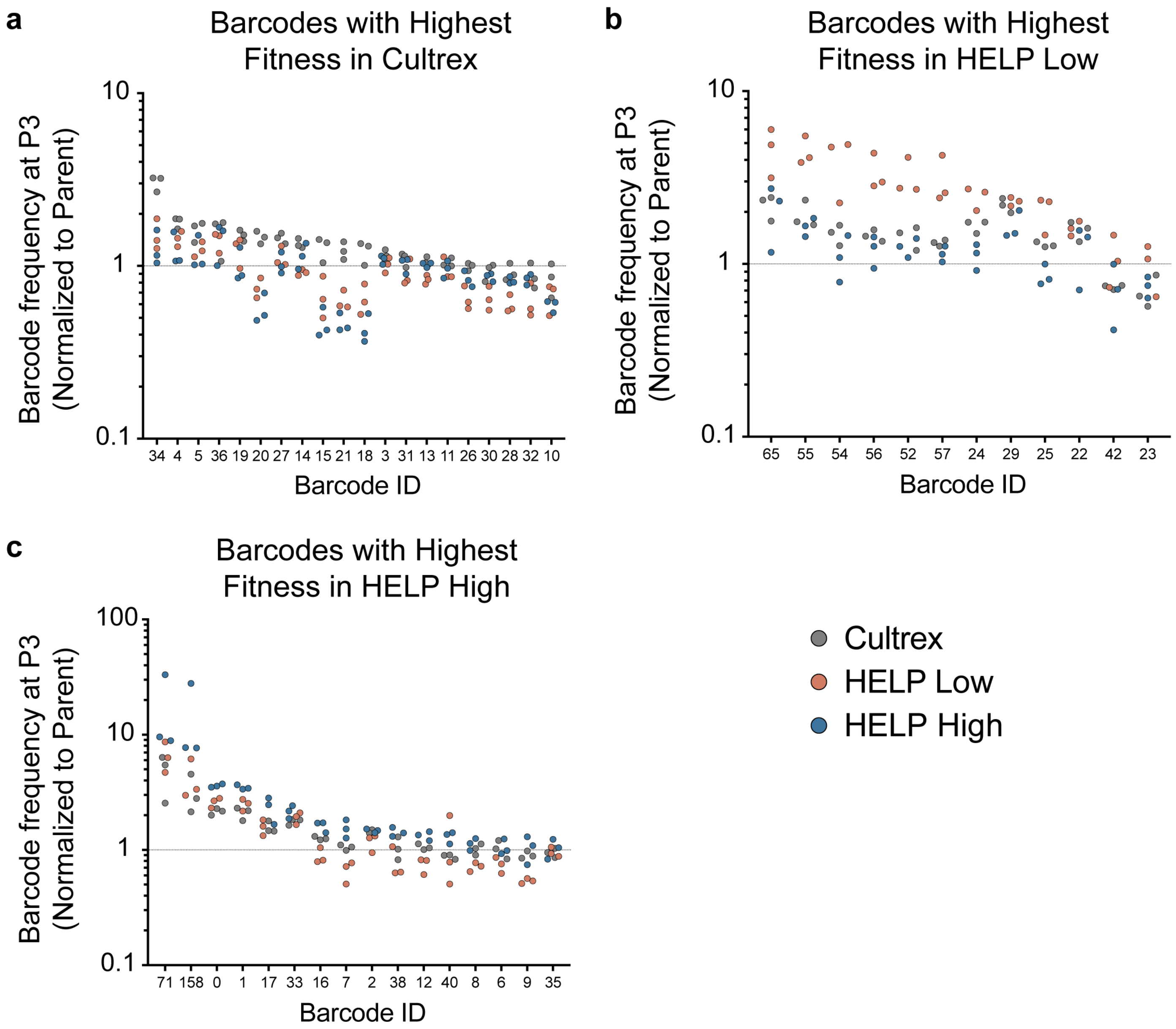


**Supplementary Figure 9. Unique PDAC organoid subclones have different fitness within distinct matrix environments.** For samples at passage three, the 25 most frequent barcodes present within Cultrex, HELP Low, or HELP High were identified and combined into a single list of top barcodes, then duplicates were removed, yielding a total of 48 barcodes in the superset. For each barcode in the superset, its frequency at passage three within each matrix was normalized by its initial frequency in the Parent population, and its average normalized frequency across three biological replicates was compared across organoids expanded within each matrix. The normalized frequencies of barcodes with the highest average fitness in Cultrex (**a**), HELP Low (**b**), or HELP High (**c**) were plotted together in rank order, alongside the normalized barcode frequencies from the other two matrices. For example, Barcode 34 had the highest average normalized frequency within Cultrex matrices compared to HELP Low and High, and thus is included in panel **a**. The dashed line at 1 refers to the barcode frequency in the initial Parent population (i.e. barcodes with normalized frequencies >1 were enriched within that given matrix, while barcodes with normalized frequencies <1 were diminished within that given matrix). For all data, only barcodes present across all biological replicates were considered. Each data point represents 1 biological replicate.

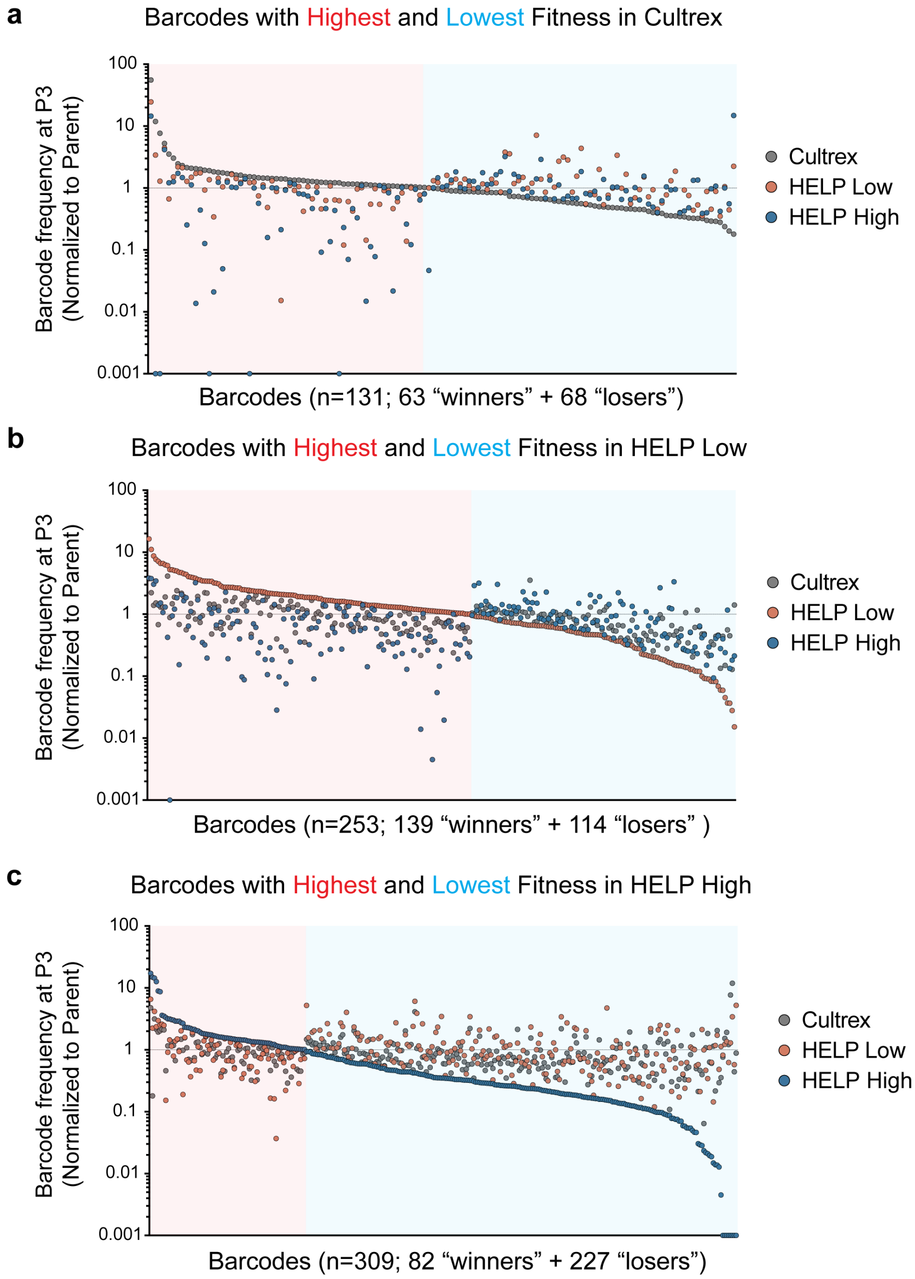


**Supplementary Figure 10. Unique PDAC organoid subclones have different fitness within distinct matrix environments.** For samples at passage three, the barcodes with frequencies greater than 0.01% within Cultrex, HELP Low, or HELP High were identified and combined into a single list of top barcodes, then duplicates were removed, yielding a total of 477 barcodes in the superset. For each barcode in the superset, its frequency at passage three in each matrix was normalized by its initial frequency in the Parent population, and its average normalized frequency across three biological replicates was compared across organoids expanded within each matrix. The normalized average frequencies of barcodes with the highest or lowest fitness within Cultrex (**a**), HELP Low (**b**), or HELP High (**c**) were plotted together in rank order, alongside the normalized average barcode frequencies from the other two matrices. For example, a barcode with the highest average normalized frequency within Cultrex matrices compared to HELP Low and High would be included in panel **a** within the red shaded region. Likewise, a barcode with the lowest normalized average fitness within Cultrex would be included in panel **a** within the blue shaded region. The dashed line at 1 refers to the barcode frequency in the initial Parent population (i.e. barcodes with normalized frequencies >1 were enriched within that given matrix, while barcodes with normalized frequencies <1 were diminished within that given matrix). For all data, only barcodes present across all biological replicates were considered. Each data point represents the average of 3 biological replicates.

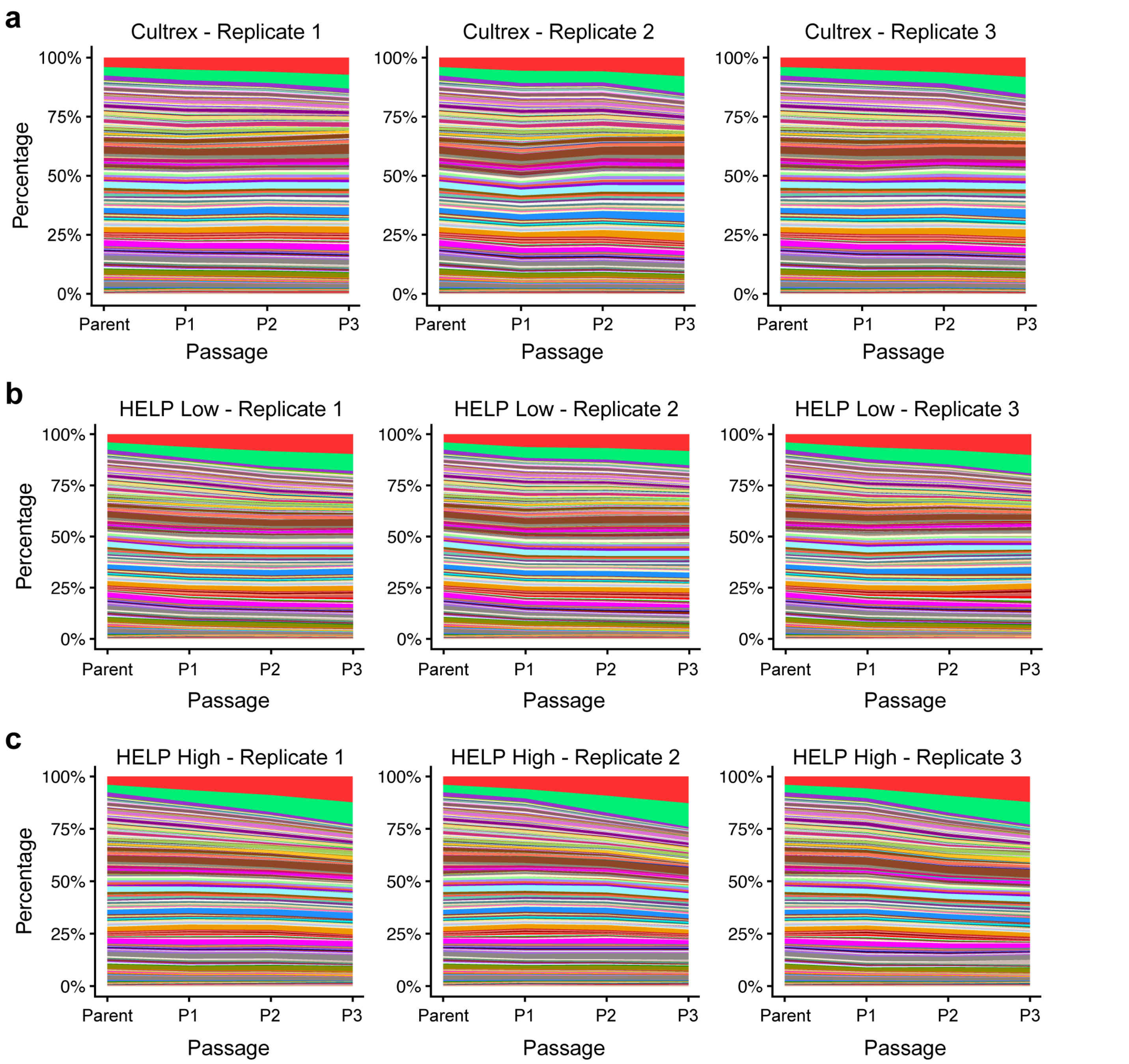


**Supplementary Figure 11. PDAC clonal heterogeneity is broadly maintained during expansion within Cultrex and HELP matrices.** Replicate Muller plots of barcode frequencies from PDAC organoids expanded within Cultrex (**a**), HELP Low (**b**), and HELP High (**c**) matrices for three passages. Colors corresponding to unique barcodes are maintained across replicates and matrix types. For example, the red colored band for Cultrex Replicate 1 represents the same barcoded subclone as the red colored band in HELP High Replicate 3.

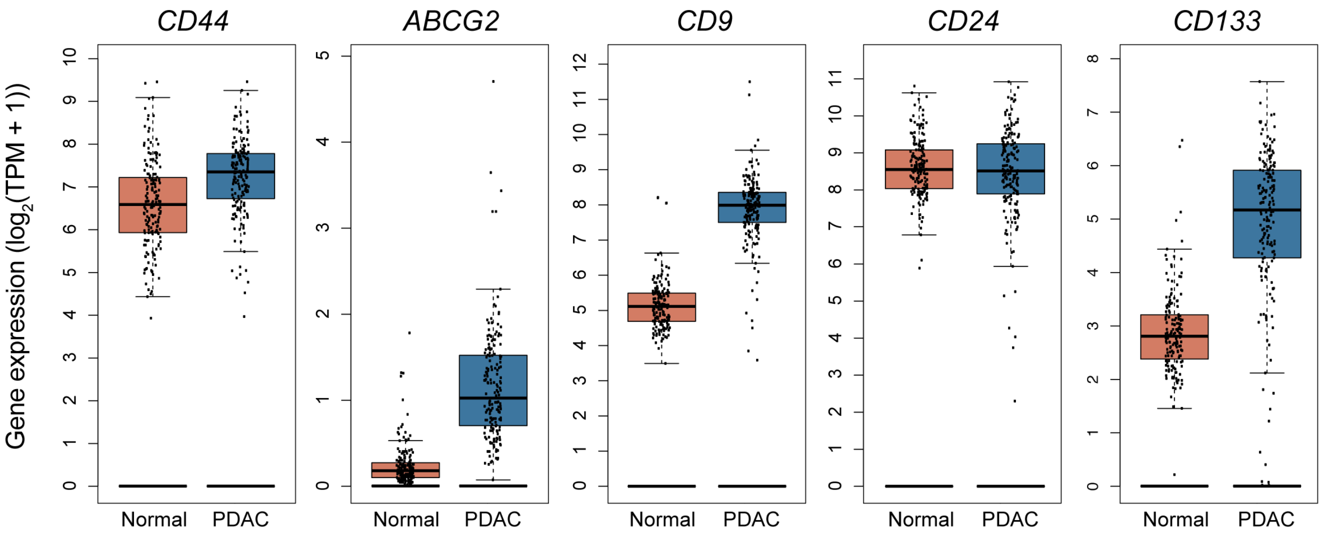


**Supplementary Figure 12. Human normal pancreas and PDAC tumor tissue bulk RNA expression of cancer stem cell markers.** Bulk RNA-sequencing analysis of PDAC and normal pancreatic tissue samples for cancer stem cell markers (N=179 for PDAC, N=171 for Normal).

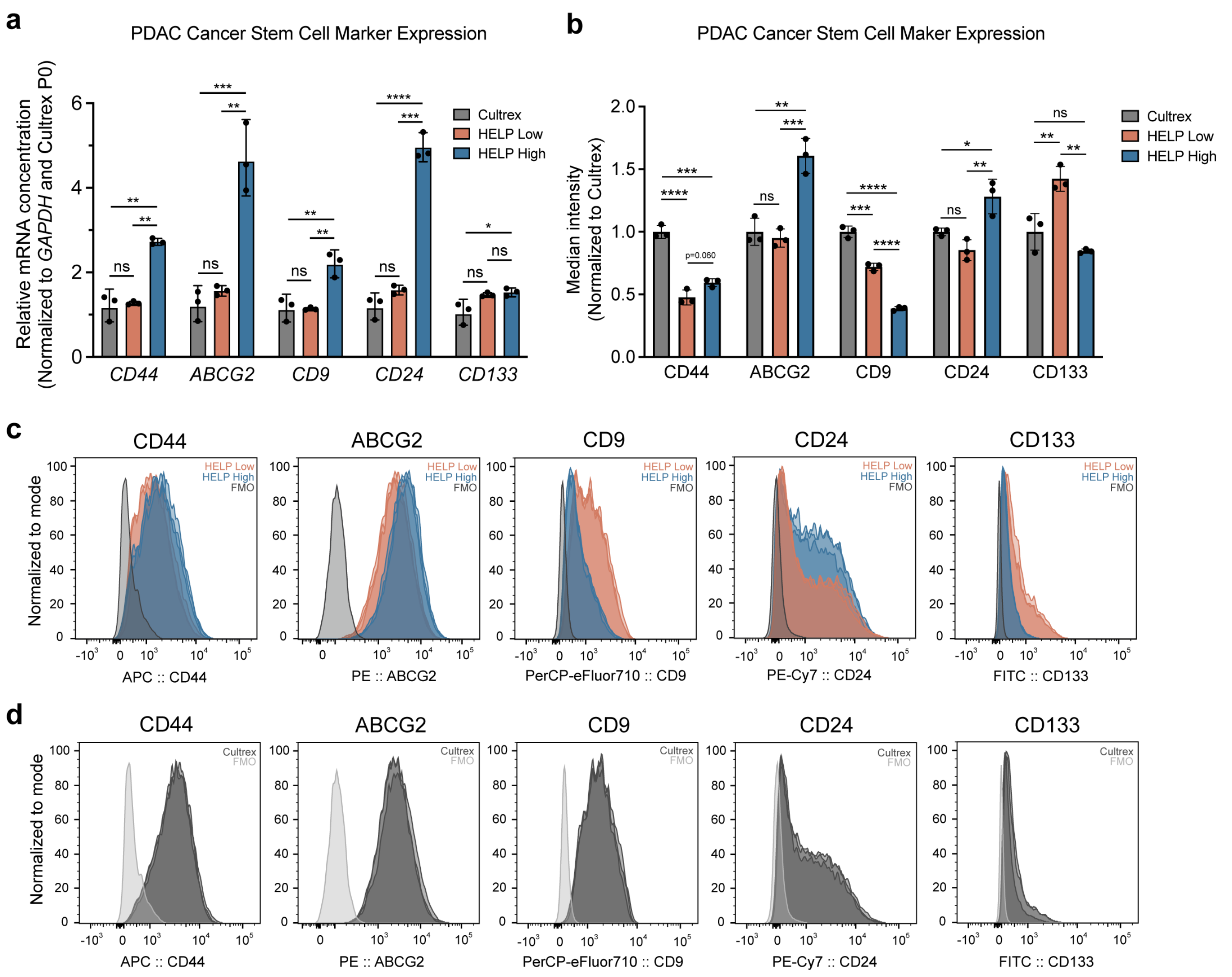


**Supplementary Figure 13. Cancer stem cell markers are upregulated within high stiffness matrices.** **a**, qPCR quantification of mRNA-level CSC marker expression in PDAC organoids expanded within Cultrex, HELP Low, or HELP High matrices for four passages (N=3, mean ± 95% confidence interval). Data are normalized to *GAPDH* gene expression and respective marker expression in the PDAC organoid parent population cultured within Cultrex prior to expansion within HELP (i.e. Cultrex P0). **b**, Median intensity of CSC marker expression measured via flow cytometry of PDAC organoids expanded within Cultrex, HELP Low, or HELP High matrices for four passages (N=3, mean ± SD). Data are normalized to PDAC organoids expanded within Cultrex. In **a** and **b**, statistical analysis was performed using an ordinary one-way ANOVA with Tukey multiple comparisons correction (*p<0.05, **p<0.01, ***p<0.001, ****p<0.0001, ns = not significant). **c,d**, Flow cytometry analysis of CSC marker expression in PDAC organoids expanded within HELP Low and HELP High (**c**) and Cultrex (**d**) matrices (N=3). Fluorescence minus one (FMO) controls are included for each marker in light grey.

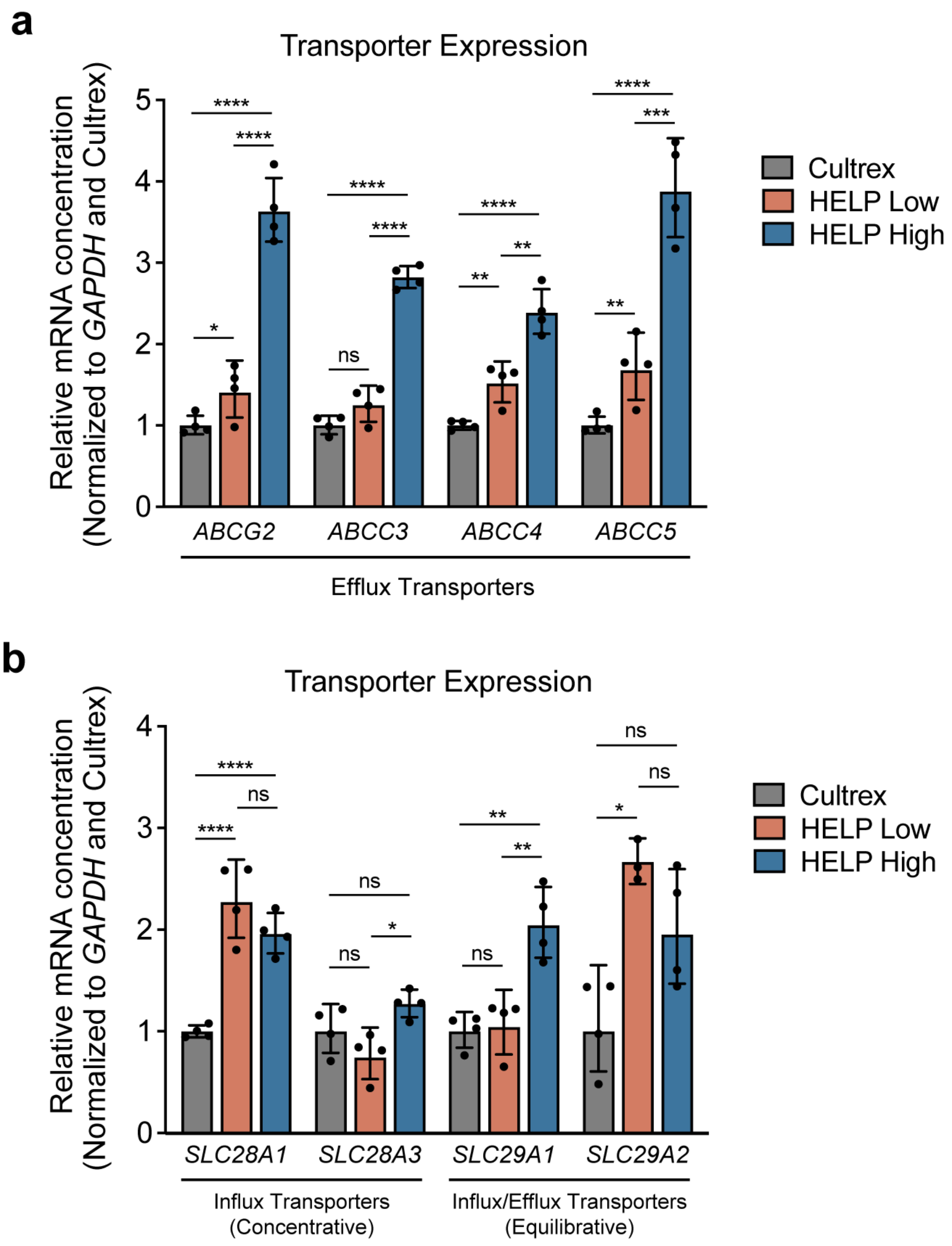


**Supplementary Figure 14. PDAC organoids alter their drug transporter expression within high stiffness matrices. a**, qPCR quantification of mRNA-level ATP-binding cassette (ABC) family drug efflux transporter expression in PDAC organoids expanded within Cultrex or HELP matrices for four passages (N=4, mean ± 95% confidence interval). **b**, qPCR quantification of mRNA-level solute carrier (SLC) family drug transporter expression in PDAC organoids expanded within Cultrex or HELP matrices for four passages (N=4, mean ± 95% confidence interval). In **a** and **b**, statistical analysis was performed using an ordinary one-way ANOVA with Tukey multiple comparisons correction (*p<0.05, **p<0.01, ***p<0.001, ****p<0.0001, ns = not significant). All data are normalized to *GAPDH* gene expression and respective marker expression in PDAC organoids expanded within Cultrex.

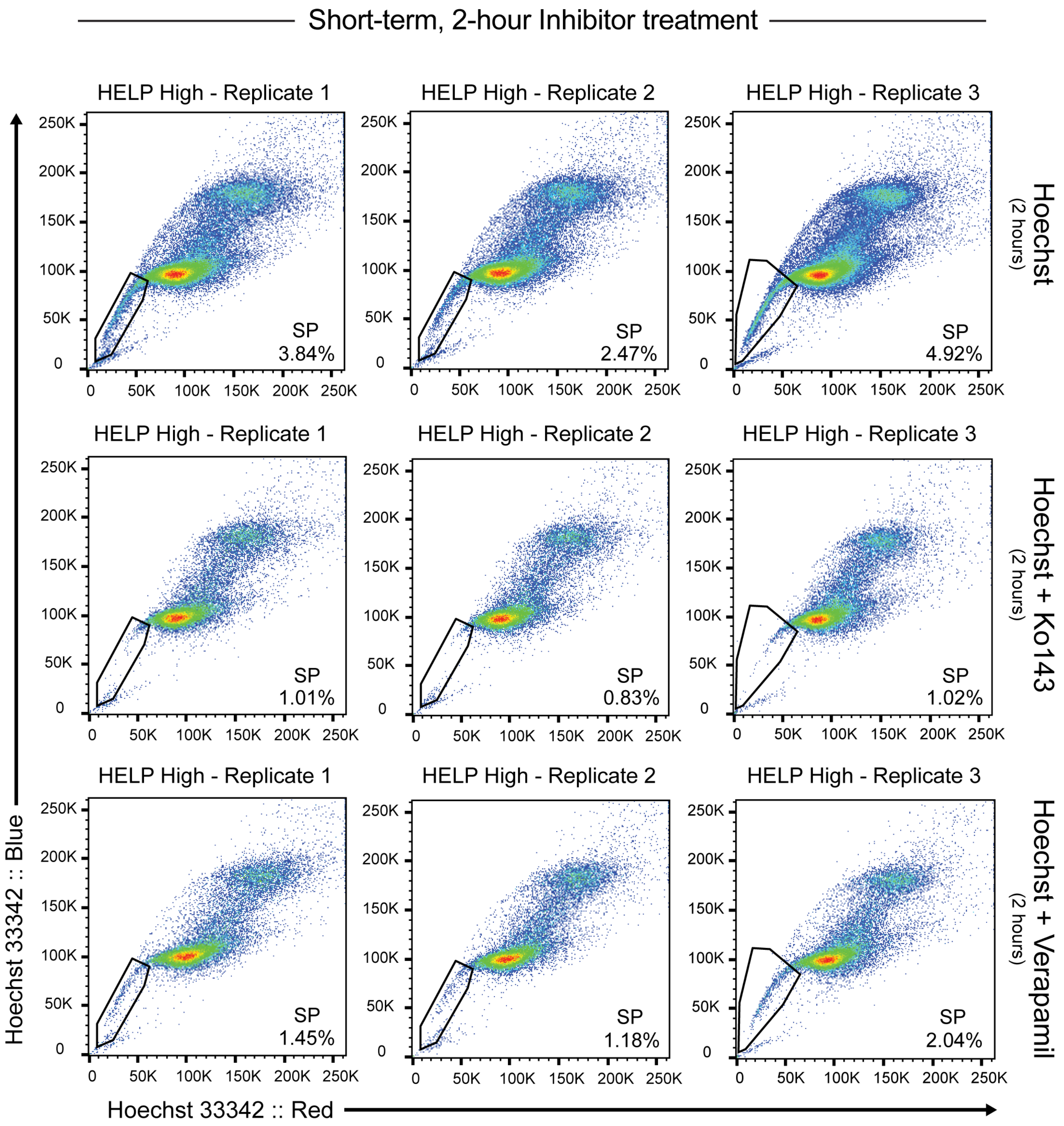


**Supplementary Figure 15. Side population analysis of PDAC organoids expanded within HELP High matrices.** Replicate side population (SP) flow cytometry analysis of PDAC organoids expanded within HELP High matrices for four passages. Prior to measurement, PDAC organoids were dissociated into single cells and treated for two hours with Hoechst (top), Hoechst + Ko143 (ABCG2 efflux transporter inhibitor; middle), or Hoechst + Verapamil (non-specific efflux transporter inhibitor; bottom). Percent of cells within the SP is labeled for each graph. Verapamil treated samples were used to confirm gating strategy for identification of the SP.

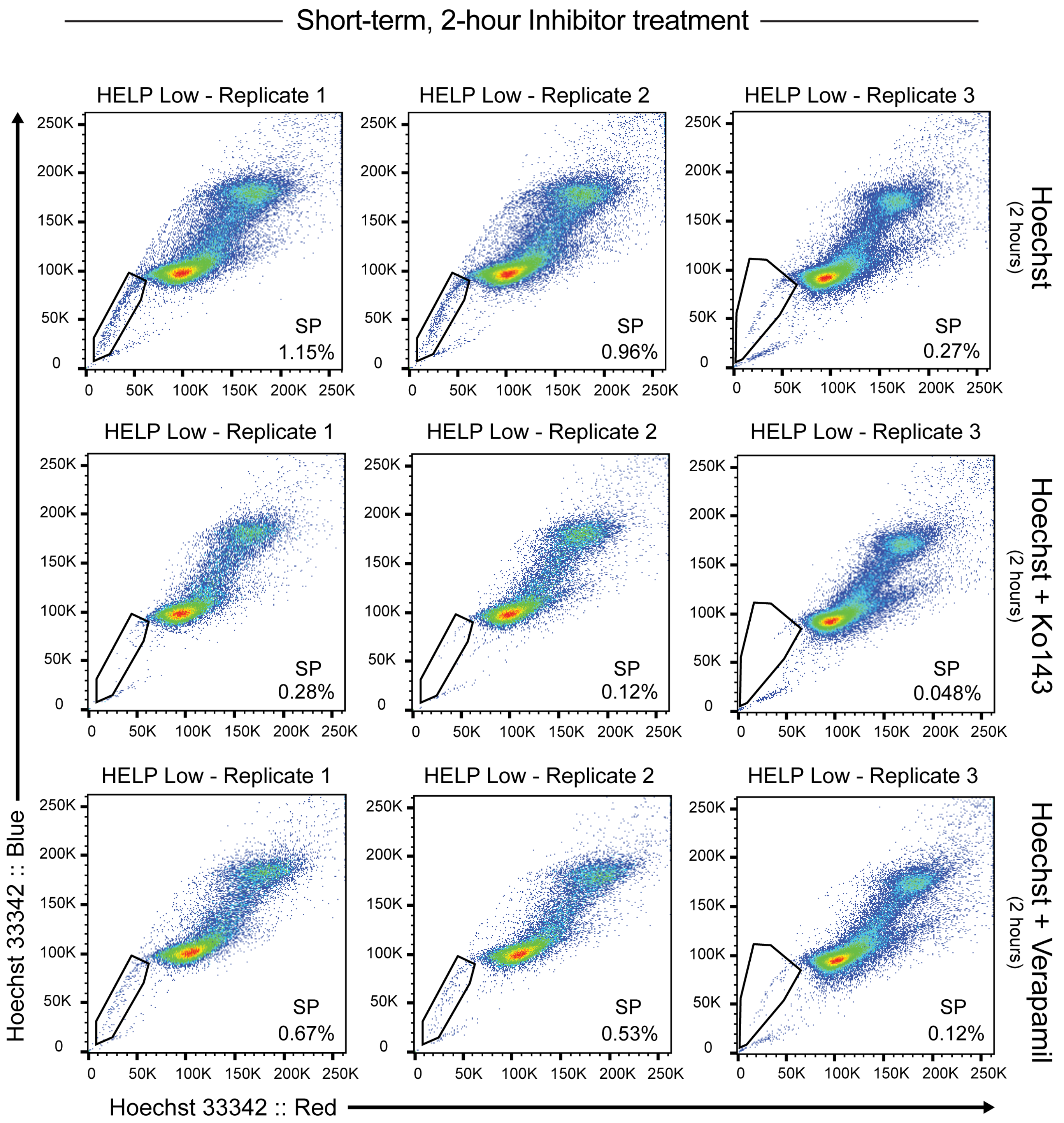


**Supplementary Figure 16. Side population analysis of PDAC organoids expanded within HELP Low matrices.** Replicate side population (SP) flow cytometry analysis of PDAC organoids expanded within HELP Low matrices for four passages. Prior to measurement, PDAC organoids were dissociated into single cells and treated for two hours with Hoechst (top), Hoechst + Ko143 (ABCG2 efflux transporter inhibitor; middle), or Hoechst + Verapamil (non-specific efflux transporter inhibitor; bottom). Percent of cells within the SP is labeled for each graph. Verapamil treated samples were used to confirm gating strategy for identification of the SP.

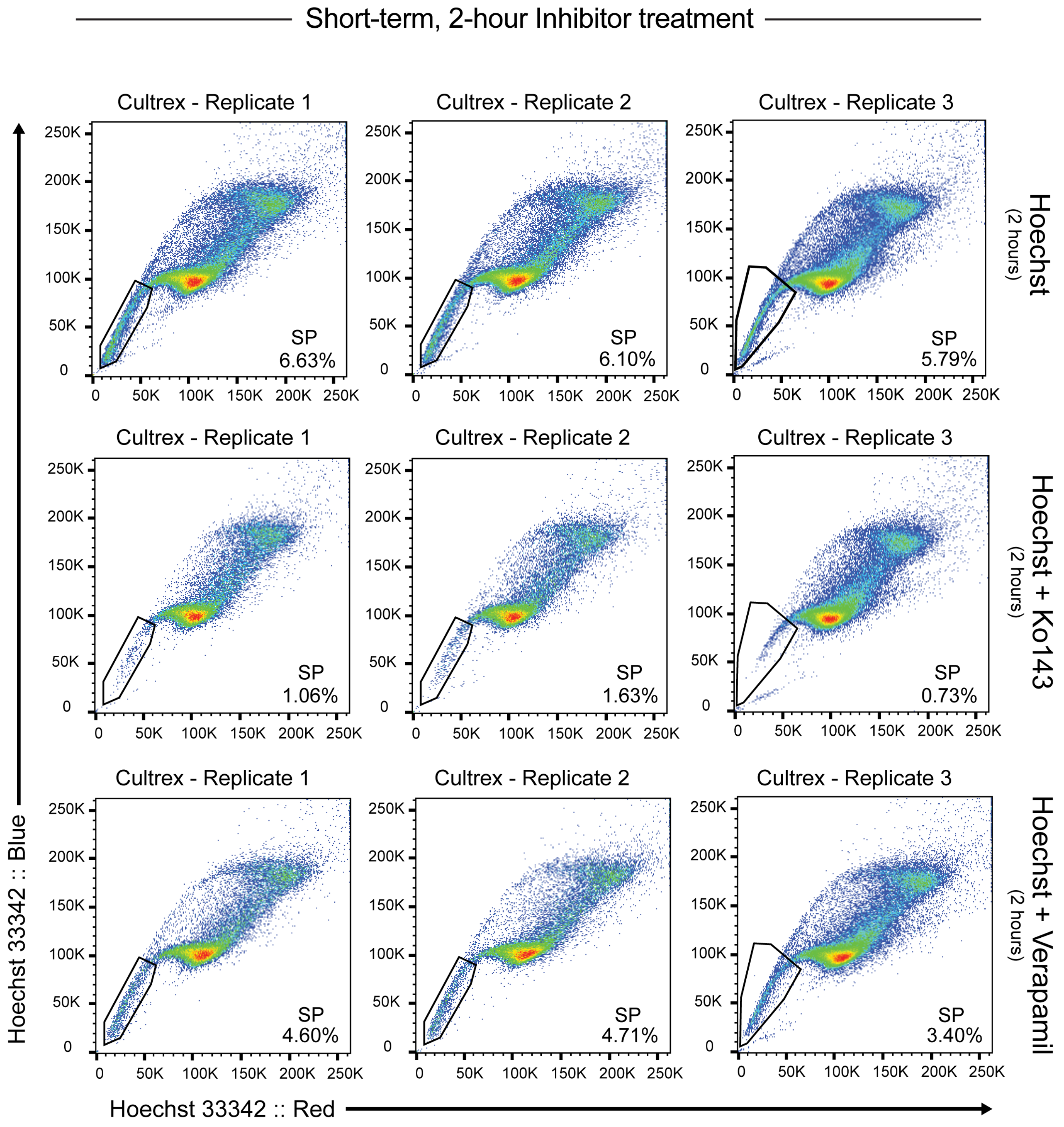


**Supplementary Figure 17. Side population analysis of PDAC organoids expanded within Cultrex matrices.** Replicate side population (SP) flow cytometry analysis of PDAC organoids expanded within Cultrex matrices for four passages. Prior to measurement, PDAC organoids were dissociated into single cells and treated for two hours with Hoechst (top), Hoechst + Ko143 (ABCG2 efflux transporter inhibitor; middle), or Hoechst + Verapamil (non-specific efflux transporter inhibitor; bottom). Percent of cells within the SP is labeled for each graph. Verapamil treated samples were used to confirm gating strategy for identification of the SP.

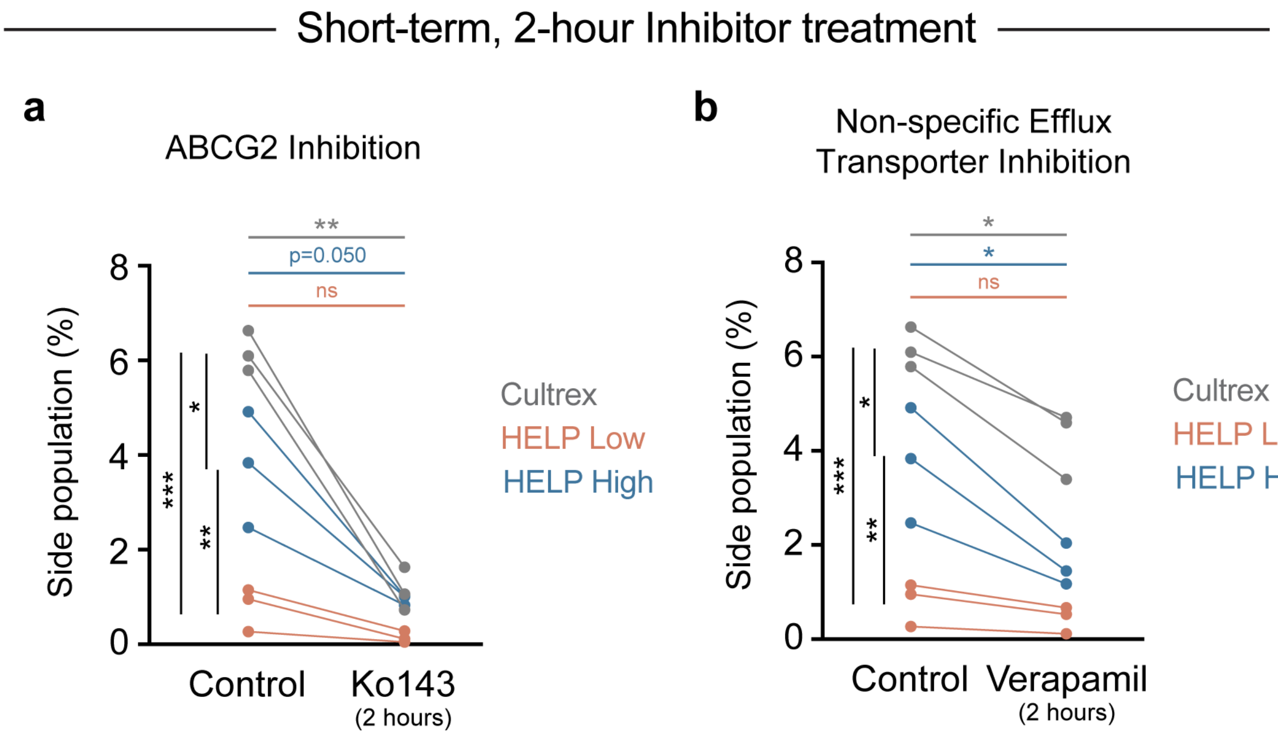


**Supplementary Figure 18. Short-term inhibition of drug efflux transporters decreases PDAC organoid side population. a**, Quantification of side population (SP) cells for PDAC organoids expanded within Cultrex or HELP matrices for four passages and treated with no inhibitor (control) or with Ko143 (ABCG2 efflux transporter inhibitor) for 2 hours (N=3). **b**, Quantification of SP cells for PDAC organoids expanded within Cultrex or HELP matrices for four passages and treated with no inhibitor (control) or with Verapamil (non-specific efflux transporter inhibitor) for 2 hours (N=3). In **a** and **b**, each data point represents 1 biological replicate and data points connected by a line are from the same population of cells. Statistical analysis individually comparing "control" vs. "inhibitor" for each matrix was performed using a paired t-test (color-matched statistical bars, *$p<0.05$, ***$p<0.001$, ns = not significant). Statistical analysis comparing percentage of SP cells in control samples across matrices was performed using an ordinary one-way ANOVA with Tukey multiple comparisons correction (black statistical bars, *$p<0.05$, **$p<0.01$, ***$p<0.001$, ns = not significant).

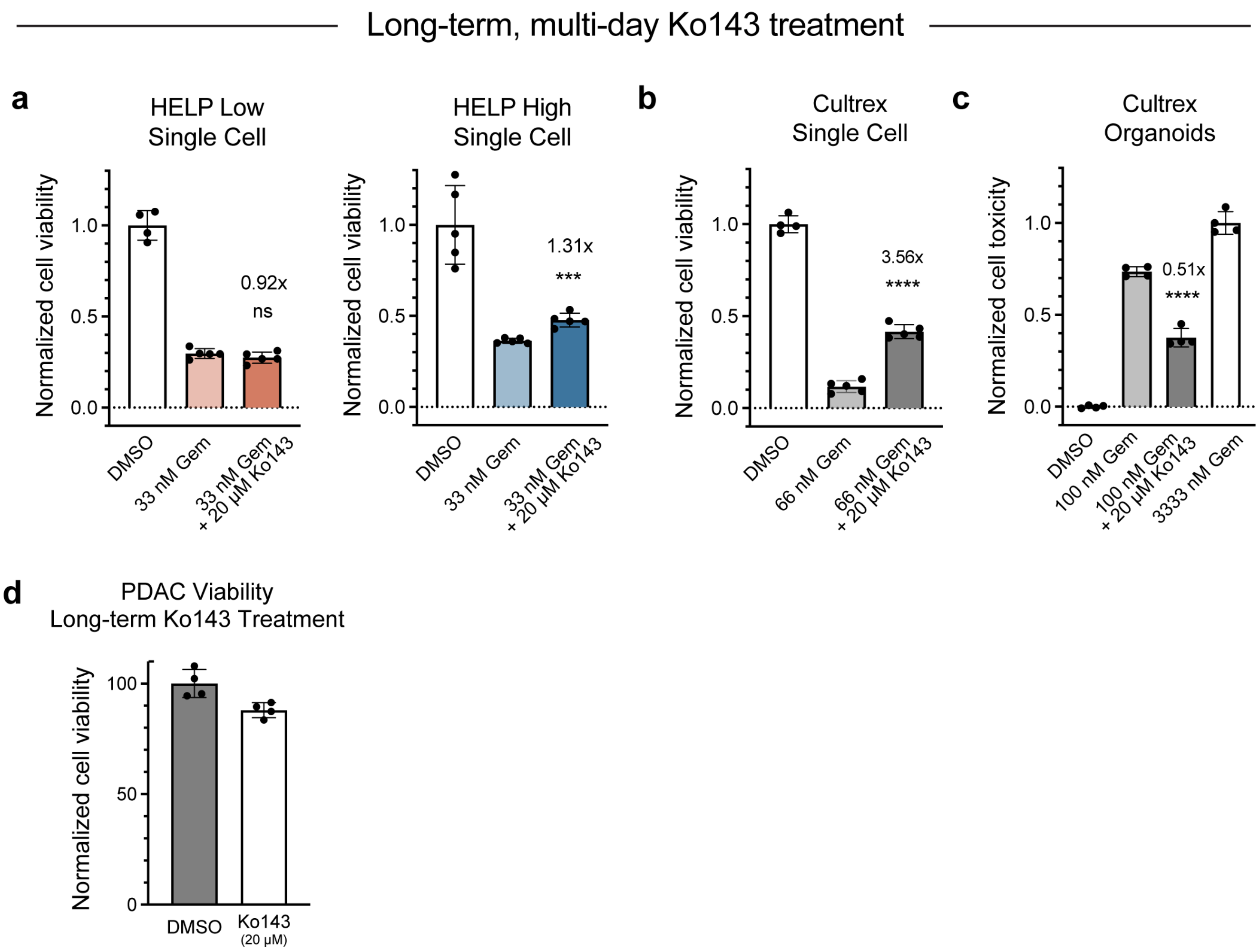


**Supplementary Figure 19. Long-term inhibition of drug efflux transporters promotes gemcitabine resistance in PDAC organoids.** **a**, PDAC viability following treatment with DMSO (control, normalized to 1), 33 nM gemcitabine (Gem), or 33 nM gemcitabine + 20 μM Ko143 (N=4-5, mean ± SD). PDAC organoids were expanded for four passages within HELP Low (left) or HELP High (right) prior to drug (+ inhibitor) treatment during log-phase growth of single cells for six days. **b**, PDAC viability following treatment with DMSO (control, normalized to 1), 66 nM gemcitabine, or 66 nM gemcitabine + 20 μM Ko143 (N=4-5, mean ± SD). PDAC organoids were expanded for four passages within Cultrex prior to drug (+ inhibitor) treatment for six days during log-phase growth of single cells. **c**, PDAC toxicity following treatment with DMSO (control, normalized to 0), 100 nM gemcitabine, 100 nM gemcitabine + 20 μM Ko143, or 3333 nM gemcitabine (positive control, normalized to 1) (N=4, mean ± SD). PDAC organoids were expanded for four passages within Cultrex prior to drug (+ inhibitor) treatment for three days following the formation of ~75-μm diameter multicellular organoids. In **a**-**c**, statistical analysis comparing experimental gemcitabine treatment and gemcitabine + Ko143 treatment was performed using an unpaired two-tailed Student's t-test (***$p<0.001$, ****$p<0.0001$, ns = not significant). The fold change in average normalized cell viability/toxicity between samples treated with gemcitabine, with and without Ko143, is reported for each comparison. **d**, PDAC viability following treatment with DMSO (control, normalized to 1) or 20 μM Ko143(N=4, mean ± SD). PDAC organoids were expanded for four passages within Cultrex matrices prior to Ko143 treatment during log-phase growth for six days. In **a** through **d**, Ko143 was added fresh to PDAC media, and media was changed daily.

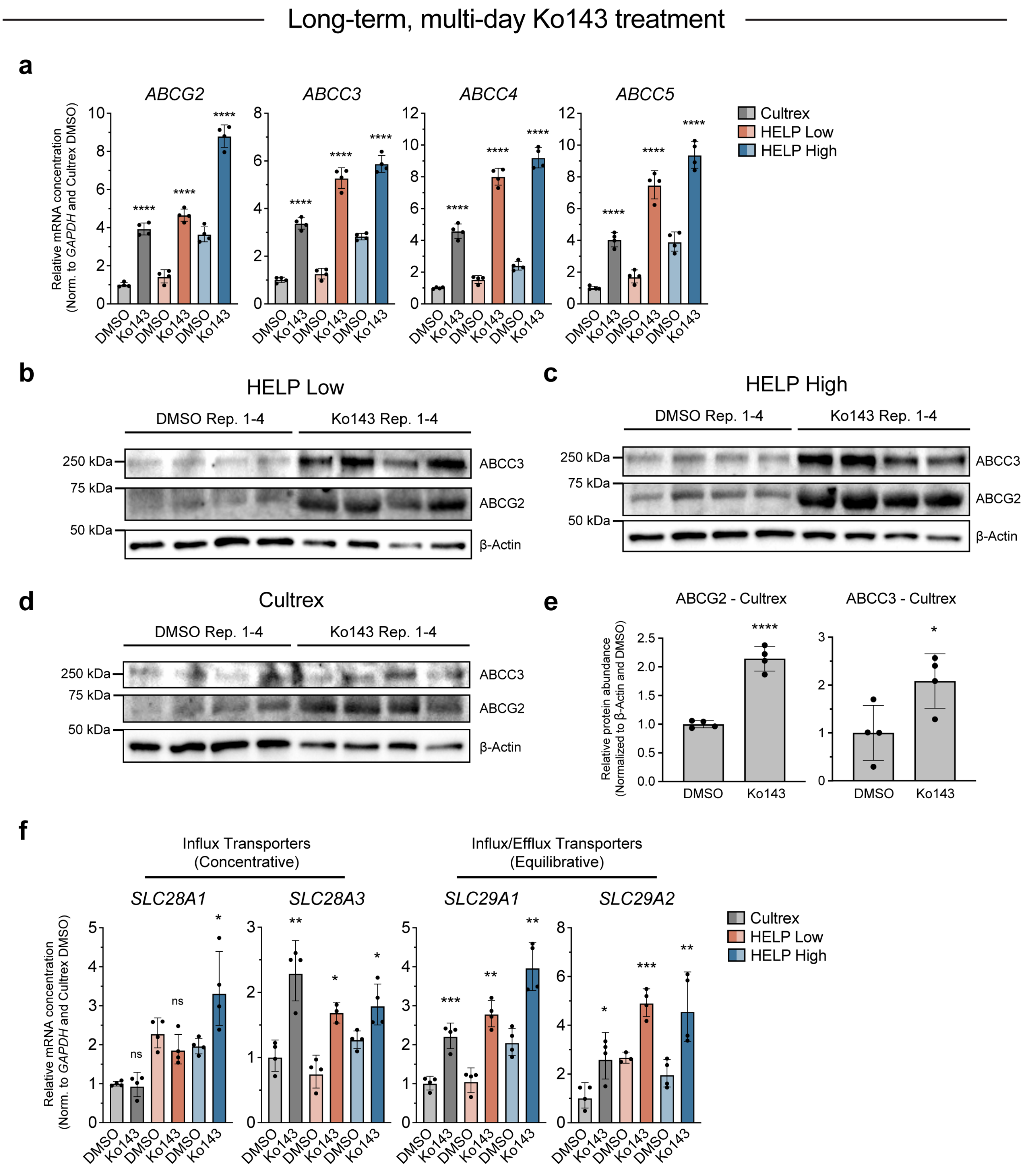

Long-term, multi-day Ko143 treatment
a
ABCG2
ABCC3
ABCC4
ABCC5
Relative mRNA concentration (Norm. to GAPDH and Cultrex DMSO)
Cultrex
HELP Low
HELP High
DMSO
Ko143
****
b
HELP Low
DMSO Rep. 1-4
Ko143 Rep. 1-4
250 kDa
75 kDa
50 kDa
ABCC3
ABCG2
β-Actin
c
HELP High
d
Cultrex
e
ABCG2 - Cultrex
ABCC3 - Cultrex
Relative protein abundance (Normalized to β-Actin and DMSO)
f
Influx Transporters (Concentrative)
Influx/Efflux Transporters (Equilibrative)
SLC28A1
SLC28A3
SLC29A1
SLC29A2
ns

**Supplementary Figure 20. PDAC organoids respond to long-term drug efflux transporter inhibition by increasing expression of a broad range of drug efflux transporters.** **a**, qPCR quantification of mRNA-level ATP-binding cassette (ABC) family drug efflux transporter expression in PDAC organoids expanded within Cultrex or HELP matrices for four passages and treated with either DMSO or 20 μM Ko143 for six days (N=4, mean ± 95% confidence interval). **b-d**, Western blot analysis of ABC-family drug efflux transporter expression in PDAC organoids expanded within HELP Low (**b**), HELP High (**c**), or Cultrex (**d**) matrices for four passages and treated with either DMSO or 20 μM Ko143 for six days (N=4). β-Actin expression was used as a loading control. **e**, Western blot quantification of data in **d** (N=4, mean ± SD). Statistical analysis comparing DMSO vs. Ko143 treatment was performed using an unpaired two-tailed Student's t-test (*$p<0.05$, ***$p<0.001$). Data are normalized to β-actin expression and DMSO samples for each marker. **f**, qPCR quantification of mRNA-level solute carrier (SLC) family drug transporter expression in PDAC organoids expanded within Cultrex or HELP matrices for four passages and treated with either DMSO or 20 μM Ko143 for six days (N=3-4, mean ± 95% confidence interval). In **a** and **f**, statistical analysis individually comparing DMSO vs. Ko143 treatment for each matrix was performed using an unpaired two-tailed Student's t-test (*$p<0.05$, **$p<0.01$, ***$p<0.001$, ****$p<0.0001$, ns = not significant). All data are normalized to *GAPDH* gene expression and respective marker expression when PDAC organoids were treated with DMSO within Cultrex matrices. In **a**-**f**, Ko143 treatment was applied to PDAC cultures throughout single cell log-phase growth for six days.

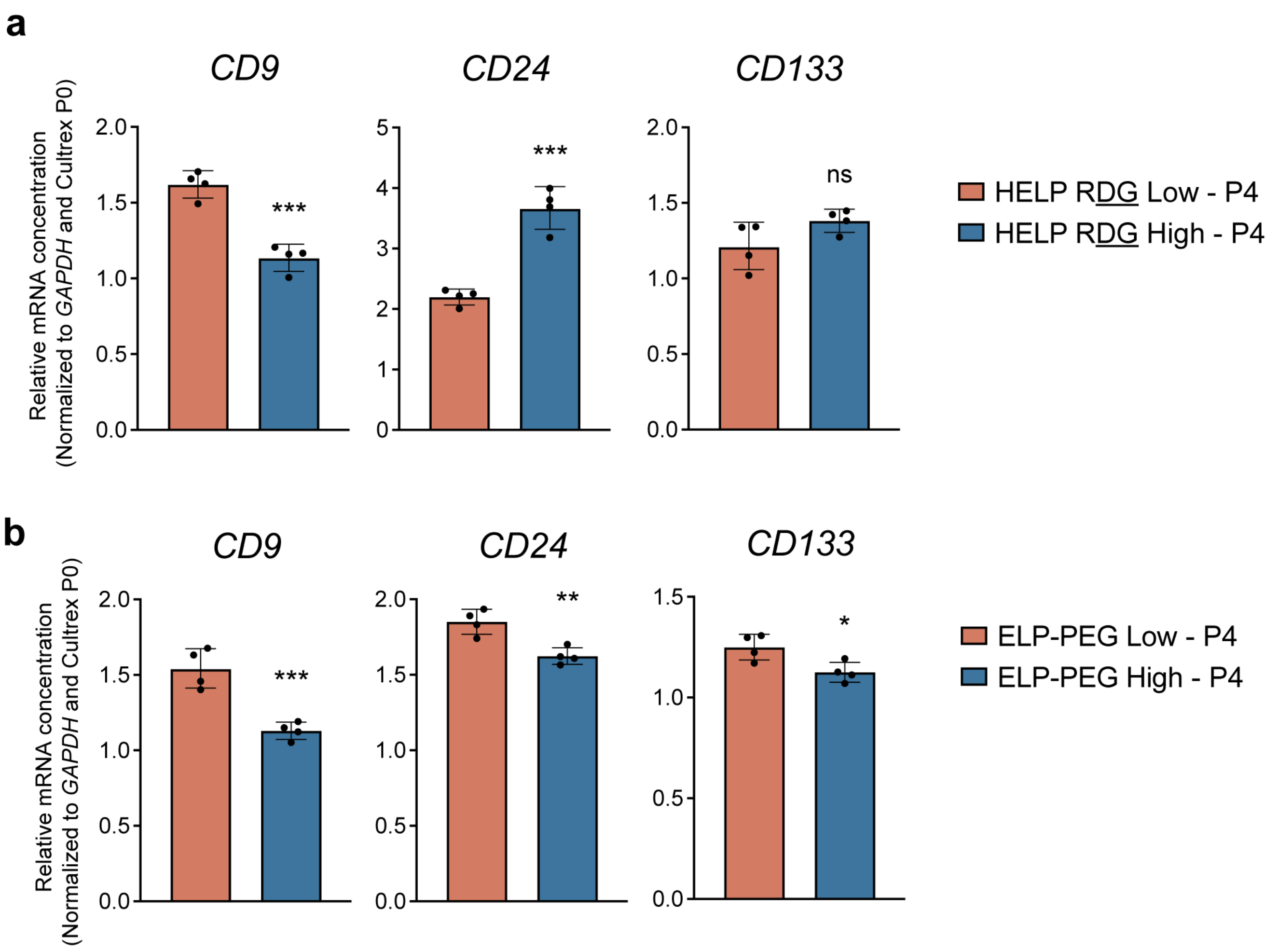


**Supplementary Figure 21. Cancer stem cell marker expression in engineered matrices without RGD ligand or hyaluronan. a**, qPCR quantification of mRNA-level CSC marker expression in PDAC organoids expanded within HELP RDG Low or HELP RDG High matrices for four passages (N=4, mean ± 95% confidence interval). **b**, qPCR quantification of mRNA-level CSC marker expression in PDAC organoids expanded within ELP-PEG Low or ELP-PEG High matrices without hyaluronan for four passages (N=4, mean ± 95% confidence interval). In **a** and **b**, statistical analysis comparing marker expression in Low vs. High matrices was performed using an unpaired two-tailed Student's t-test (*p<0.05, **p<0.01, ***p<0.001, ns = not significant). All data are normalized to *GAPDH* gene expression and respective marker expression in the PDAC organoid parent population cultured within Cultrex prior to expansion in HELP RDG or ELP-PEG (i.e. Cultrex P0).

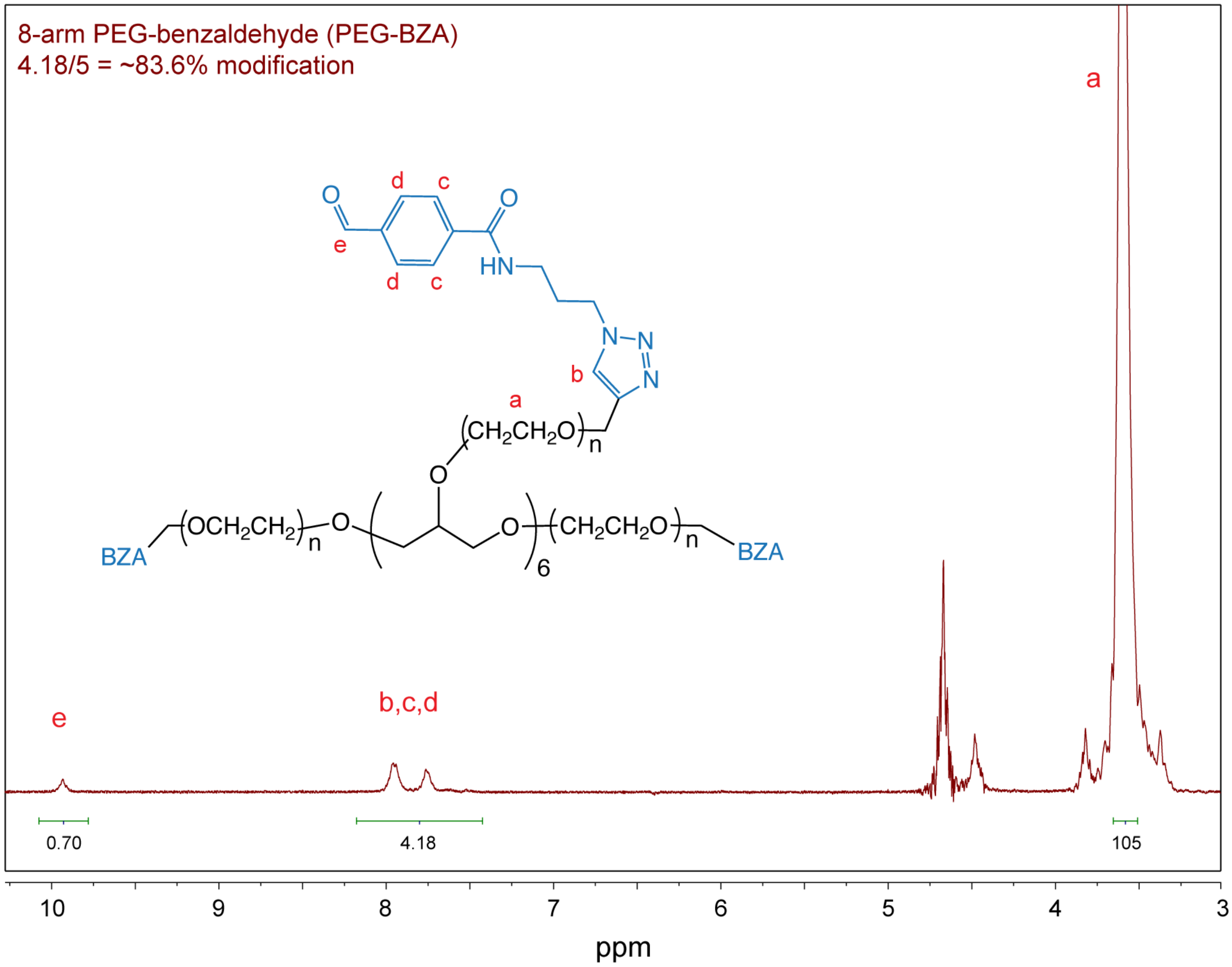


**Supplementary Figure 22. Modification of polyethylene glycol (PEG) with a benzaldehyde functional group.** Representative $^{1}H$ NMR spectrum ($D_2O$ solvent) of a 10-kDa 8-arm star PEG molecule modified with a benzaldehyde functional group.

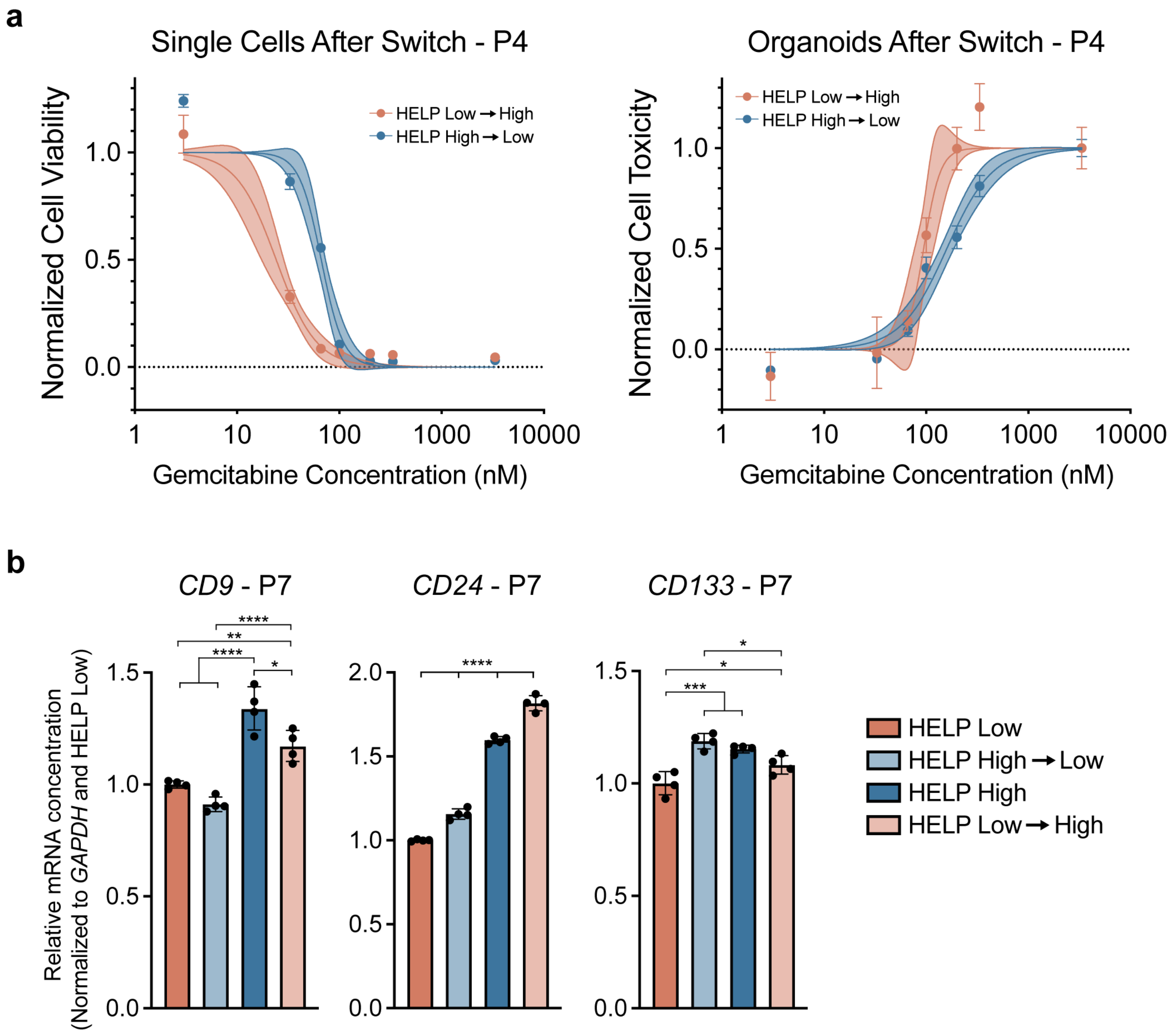


**Supplementary Figure 23. PDAC organoid chemoresistance and CSC marker expression is reversible. a**, Single cell- (left) and organoid-level (right) gemcitabine dose-response curves for PDAC organoids expanded within HELP Low or HELP High matrices for three passages and then switched to the opposite stiffness matrix for an additional passage (i.e. passage 4). Each data point represents the mean ± SEM (N=4, solid center line is nonlinear least squares regression of data; shaded region represents 95% confidence bands of nonlinear fit; data are normalized to positive controls (DMSO for single cells; 3333 nM gemcitabine for organoids)). **b**, qPCR quantification of mRNA-level CSC marker expression in PDAC organoids expanded within HELP Low or HELP High matrices for seven passages, with or without a matrix switch at passage four (N=4, mean ± 95% confidence interval, ordinary one-way ANOVA with Tukey multiple comparisons correction, *p<0.05, **p<0.01, ***p<0.001, ****p<0.0001, unlabeled comparisons are not significantly different). All data are normalized to *GAPDH* gene expression and respective marker expression within HELP Low at passage seven.

**Supplementary Table 1. Extracellular Matrix Formulations**

| Hydrogel Formulation | [ELP-hydrazine] (wt%) | [HA-benzaldehyde] (wt%) | [PEG-benzaldehyde] (wt%) | Stiffness (Pa) ± SD | [RGD] (mM) |
|---|---|---|---|---|---|
| **Cultrex** | | | | | |
| Reduced Growth Factor, Type 2 | - | - | - | 84 ± 27 | - |
| **HELP** | | | | | |
| HELP Low | 1 (RGD) | 1 (~7% modification) | - | 279 ± 68 | 1.05 |
| HELP Medium | 1 (RGD) | 1 (~34% modification) | - | 1253 ± 162 | 1.05 |
| HELP High | 1 (RGD) + 1 (RDG) | 1 (~34% modification) | - | 3040 ± 176 | 1.05 |
| **ELP-PEG** | | | | | |
| ELP-PEG Low | 1 (RGD) + 0.5 (RDG) | - | 1 (10 kDa, 8-arm) | 274 ± 99 | 1.05 |
| ELP-PEG High | 1 (RGD) + 1.5 (RDG) | - | 1.75 (10 kDa, 8-arm) | 2910 ± 142 | 1.05 |

**Supplementary Table 1.** Summary of hydrogel formulations and their relevant parameters used throughout the study.

**Supplementary Table 2. Elastin-like Protein (ELP) Amino Acid Sequences**

| ELP | Amino Acid Sequence |
|---|---|
| RGD ELP | MASMTGGQQMGHHHHHHDDDDKLQLDASTVYAVTG**RGD**SPASSAASAVPGIGVPGIGVP<br>GKGVPGIGVPGIGVPGIGVPGIGVPGKGVPGIGVPGIGVPGIGVPGIGVPGKGVPGIGVPGI<br>GVPLDASTVYAVTG**RGD**SPASSAASAVPGIGVPGIGVPGKGVPGIGVPGIGVPGIGVPGIGV<br>PGKGVPGIGVPGIGVPGIGVPGIGVPGKGVPGIGVPGIGVPLDASTVYAVTG**RGD**SPASSA<br>ASAVPGIGVPGIGVPGKGVPGIGVPGIGVPGIGVPGIGVPGKGVPGIGVPGIGVPGIGVPGIG<br>VPGKGVPGIGVPGIGVPLDASTVYAVTG**RGD**SPASSAASAVPGIGVPGIGVPGKGVPGIGV<br>PGIGVPGIGVPGIGVPGKGVPGIGVPGIGVPGIGVPGIGVPGKGVPGIGVPGIGVPLE* |
| RDG ELP | MASMTGGQQMGHHHHHHDDDDKLQLDASTVYAVTG**RDG**SPASSAASAVPGIGVPGIGVP<br>GKGVPGIGVPGIGVPGIGVPGIGVPGKGVPGIGVPGIGVPGIGVPGIGVPGKGVPGIGVPGI<br>GVPLDASTVYAVTG**RDG**SPASSAASAVPGIGVPGIGVPGKGVPGIGVPGIGVPGIGVPGIGV<br>PGKGVPGIGVPGIGVPGIGVPGIGVPGKGVPGIGVPGIGVPLDASTVYAVTG**RDG**SPASSA<br>ASAVPGIGVPGIGVPGKGVPGIGVPGIGVPGIGVPGIGVPGKGVPGIGVPGIGVPGIGVPGIG<br>VPGKGVPGIGVPGIGVPLDASTVYAVTG**RDG**SPASSAASAVPGIGVPGIGVPGKGVPGIGV<br>PGIGVPGIGVPGIGVPGKGVPGIGVPGIGVPGIGVPGIGVPGKGVPGIGVPGIGVPLE* |

Note: Tag region; Fibronectin-derived integrin binding/non-binding region; Elastin-like repeat region

**Supplementary Table 2.** Amino acid sequences of recombinant elastin-like protein sequences used throughout the study.

**Supplementary Table 3. Primary antibodies**
**ICC = Immunocytochemistry, IHC = Immunohistochemistry, WB = Western blot, FC = Flow cytometry**

| Target | Host species | Supplier | Catalog Number | Dilution |
|---|---|---|---|---|
| **Antibodies** | | | | |
| ABCC3 / MRP3 | Rabbit | Cell Signaling | 14182S | WB: 1:1000 |
| ABCG2 / BCRP1 | Rabbit | Cell Signaling | 42078S | ICC: 1:200 WB: 1:1000 |
| ABCG2 / BCRP1 - PE | Mouse | Invitrogen | 12-8888-42 | FC: 5:100 |
| β-Actin | Rabbit | Cell Signaling | 4970S | WB: 1:1000 |
| CD133 - FITC | Mouse | Invitrogen | 11-1339-42 | FC: 5:100 |
| CD24 | Rabbit | Abcam | ab202073 | ICC: 1:100 |
| CD24 - PE-Cy7 | Mouse | Invitrogen | 25-0247-42 | FC: 2.5:100 |
| CD44 | Mouse | Santa Cruz Biotech | sc7297 | ICC: 1:50 |
| CD44 - APC | Rat | Invitrogen | 17-0441-81 | FC: 0.15:100 |
| CD9 - PerCP-eFluor 710 | Mouse | Invitrogen | 46-0098-42 | FC: 2.5:100 |
| Cleaved caspase-3 | Rabbit | Cell Signaling | 9664S | ICC: 1:400 |
| Cytokeratin 19 | Rabbit | Abcam | ab52625 | IHC: 1:200 ICC: 1:200 |
| Fibronectin | Rabbit | Cell Signaling | 26836S | IHC: 1:200 |
| Ki67 | Rabbit | Cell Signaling | 9129S | ICC: 1:200 |
| Ki67 | Mouse | Cell Signaling | 9449S | ICC: 1:200 |
| **Other** | | | | |
| Biotinylated Hyaluronan Binding Protein (500ug/mL stock in water) | - | MilliporeSigma | 38591150UG | IHC: 1:100 |
| DAPI (5 mg/mL stock in water) | - | MilliporeSigma | D9542 | ICC: 1:2000 IHC: 1:2000 |
| Phalloidin (100 ug/mL stock in DMSO) | - | MilliporeSigma | P1951 / P5282 | ICC: 1:500 |

**Supplementary Table 3.** Primary antibodies and reagents used for imaging, flow cytometry, and Western blot experiments.

**Supplementary Table 4. Secondary antibodies**
**ICC = Immunocytochemistry, IHC = Immunohistochemistry, WB = Western blot**

| Host and target | Fluorescence | Supplier | Catalog Number | Dilution |
|---|---|---|---|---|
| **Antibodies** | | | | |
| goat-anti-mouse | AF488 | Invitrogen | A11029 | IHC/ICC: 1:500 |
| goat-anti-mouse | AF647 | Invitrogen | A21237 | IHC/ICC: 1:500 |
| goat-anti-mouse | AF532 | Invitrogen | A11002 | IHC/ICC: 1:500 |
| goat-anti-rabbit | AF488 | Invitrogen | A11034 | IHC/ICC: 1:500 |
| goat-anti-rabbit | AF546 | Invitrogen | A11071 | IHC/ICC: 1:500 |
| goat-anti-rabbit | AF647 | Invitrogen | A32733 | IHC/ICC: 1:500 |
| HRP-donkey-anti-rabbit | - | JacksonImmuno Research | 711-035-152 | WB: 1:10,000 |
| **Other** | | | | |
| Streptavidin (1 mg/mL in PBS) | AF488 | Invitrogen | S11223 | IHC/ICC: 1:500 |

**Supplementary Table 4.** Secondary antibodies and reagents used for imaging and Western blot experiments.

**Supplementary Table 5. qPCR Primers**

| Target | Forward Primer (5' to 3') | Reverse Primer (5' to 3') |
|---|---|---|
| ABCC3 | TGGGGTGAAGTTTCGTACTGG | CACGTTTGACTGAGTTGGTGATA |
| ABCC4 | AGCTGAGAATGACGCACAGAA | ATATGGGCTGGATTACTTTGGC |
| ABCC5 | AGTCCTGGGTATAGAAGTGTGAG | ATTCCAACGGTCGAGTTCTCC |
| ABCG2 | GGTGGAGGCAAATCTTCGTTA | GAGTGCCCATCACAACATCA |
| CD133 | AGTCGGAAACTGGCAGATAGC | GGTAGTGTTGTACTGGGCCAAT |
| CD24 | GCTCCTACCCACGCAGATT | GGTGGTGGCATTAGTTGGAT |
| CD44 | CTGCCGCTTTGCAGGTGTA | CATTGTGGGCAAGGTGCTATT |
| CD9 | GATATTCGCCATTGAAATAGCTGC | TTGTAGGTGTCCTTGTAAAACTCC |
| GAPDH | CATGAGAAGTATGACAACAGCCT | AGTCCTTCCACGATACCAAAGT |
| SLC28A1 | CCTCACCTGTGTGGTCCTCA | AGACCCCTCTTAAACCAGAGC |
| SLC28A3 | CACAGAGCCCTTCCTCTTTTTG | GCCAGAACCAATGGCTGTTTAG |
| SLC29A1 | TGAGCGGAACTCTCTCAGTG | TGAGGTAGGTGAATAACAGCAGG |
| SLC29A2 | TCAGTGCAGTCCTACAGGG | GGCGTGATAAAGTACCCCAGG |

**Supplementary Table 5.** Primers used for qPCR experiments.
